# Behavioural Investigations of Psilocybin in Animals 1962-2021: A Scoping Review

**DOI:** 10.1101/2024.01.04.574146

**Authors:** Ron Shore, Kat Dobson, Nina Thomson, Nigel Barnim, Hailey Bergman, Katie Rideout, Sandra McKeown, Mary C. Olmstead, Craig Goldie, Eric Dumont

**Affiliations:** University of Groningen, Groningen, the Netherlands; Department of Biomedical and Molecular Sciences, Queen’s University; Department of Psychology, Centre for Neuroscience Studies, Queen’s University; Bracken Health Sciences Library, Queen’s University; Department of Oncology, Queen’s University

**Keywords:** psilocybin, psychedelic, non-human animal research, scoping review, behavioural research, Research Domains Criteria

## Abstract

**Background and Aims:** Psilocybin is a psychedelic drug that may hold promise for a wide range of human health conditions, yet the identification of therapeutic processes and mechanisms of action remains exploratory. We conducted a scoping review on pre-clinical behavioural investigations of psilocybin in non-human animals to help determine the behavioural effects of psilocybin in non-human animals, to identify studies completed, behavioural tests employed, and what dosing modalities had been studied.

**Methods:** A librarian-conducted literature search was performed using predefined key terms and search criteria and additional searching was conducted by reviewers, using electronic databases, grey literature sources, and reference lists of relevant articles or reviews. The final search updated occurred in October, 2021. Studies were reviewed, screened and selected against an a priori protocol using Covidence software by multiple reviewers with results plotted across the Research Domains Criteria construct.

**Results:** From 4124 records identified by database searching, 260 publications were subjected to full-text review with 77 studies included in this scoping review, published between 1962-2021. The preponderance of studies (n=64) investigated behavioural outcomes in rodents. Only 43 studies (55.8%) reported on housing conditions, and seventeen studies (22.1%) failed to report sample size. All studies reported behavioural outcomes following drug administration, with fifty-one studies (66.2%) using psilocybin, thirty studies (42.9%) psilocin, four studies (5.2%) administering whole mushroom extracts (WME), and a further eight studies investigating both psilocybin and psilocin and one study reporting the effects of both psilocin and WME. One hundred and thirty distinct behavioural investigations using fifty different behavioral paradigms were identified. Few adverse events were reported, and even exceedingly high doses were apparently well tolerated.

**Conclusion:** With seventy-seven publications spanning close to sixty years, there is huge variation in study design and quality. Overall psilocybin presents a unique and strong safety profile with no evidence of biological toxicity, is characterized by unique time and dose-dependent effects, and its pattern of drug action is significantly context and training-sensitive. Data suggest putative effects of psilocybin include acute arousal, dose-dependent sedation, reductions in fear conditioning at low doses, reduced aggression, improved valence, acute disruption of working memory, the rescuing of deficits from chronic stress, and improved learning when combined with repeated environmental exposure after resolution of drug effect.

## 1. INTRODUCTION

### 1.1 Psilocybin

Psilocybin (4-phosphoryloxy-*N,N*-dimethyltryptamine) is a naturally occurring tryptamine indolealkylamine and a prodrug to psilocin (4-Hydroxy-N,N-dimethyltryptamine), a serotonin 5HT_2A_ receptor agonist. Along with lysergic acid diethylamide (LSD) and N, N-Dimethyltryptamine (DMT), psilocybin is considered a classic serotonergic psychedelic and is structurally related to the monoamine neurotransmitter serotonin (David E Nichols, 2016; Tylš et al., 2014). With an indole nucleus at their basic structure, indoleamines have high structural similarity with the endogenous tryptamine serotonin (5-hydroxytryptamine; 5-HT), a monoamine neurotransmitter and neuromodulator (synthesized from the essential amino acid tryptophan) with a wide range of biologic functions (Araújo et al., 2015; David E Nichols, 2018).

5-HT receptors regulate a range of processes including contraction of smooth muscle, learning and memory, sleep and wake cycles, thermoregulation, appetite, sexual behaviour, pain, motor activity and aspects of autonomic function (Flanagan & Nichols, 2018). 5-HT_2A_ receptors principally expressed in the cortex are implicated in the novel patterns of global neurological connectivity (Robin L. Carhart-Harris et al., 2012), entropy (Robin L. Carhart-Harris et al., 2014) or cortical desynchronization (Muthukumaraswamy et al., 2013) produced by psilocybin. When combined with the appropriate clinical preparation and setting, psilocybin may result in a subsequent and beneficial “rewiring” of brain networks away from previous pathological patterns and more similar to pre-disease states (Robin L. Carhart-Harris et al., 2017; Robin L. Carhart-Harris & Goodwin, 2017; D. E. Nichols et al., 2017) characterized by positive mood and sustained improvements in mental health (Calvey & Howells, 2018; de Gregorio et al., 2021; De Gregorio et al., 2018).

The therapeutic effects of classical psychedelics have been variously proposed to be psychological mechanisms such as ego dissolution (Smigielski et al., 2019), mysticism (Griffiths et al., 2016), spirituality (Lafrance et al., 2021), existential intelligence (Tupper, 2002), or relaxed expectations and beliefs (R. L. Carhart-Harris & Friston, 2019). Various underlying neurobiological mechanisms have been identified in previous research, including alterations to thalamic gating (Vollenweider, 2001), reintegrated cortical to sub-cortical brain network connectivity (Doss et al., 2021), and neuroplasticity (de Vos et al., 2021). Psilocybin has been trialed for therapeutic effects in a range of human health conditions, including depression, substance use disorder, obsessive compulsive disorder, anxiety and distress in advanced cancer patients, demoralization, and migraine headaches (Shore, 2019).

Psilocybin is rapidly converted to psilocin *in vivo* (Horita & Weber, 1962). Equimolar doses of psilocin and psilocybin are believed to yield comparable behavioural effects (Davis & Walters, 1977). The relative potency of psilocin to psilocybin is estimated to be 1:1.4 based on the ratio of their molecular weights: 0.71 mg/kg psilocin is equivalent to 1 mg/kg psilocybin and believed to result in similar behavioural effects (Wolbach et al., 1962). The relative potency of whole mushroom extracts is unclear, as psilocybin-containing mushrooms typically contain a variety of other potentially bioactive alkaloids in addition to psilocybin, and psilocin in varying concentrations resulting from strain as well as growing and drying conditions (Wieczorek et al., 2015). When reporting behavioural outcomes in this review, studies using psilocybin and psilocin are compared by converting the treatment dose of one compound to its equivalent dose in the other.

### 1.2 Non-human Animal Studies as Translational Models for Psychedelic Research

Behavioural models of animal study are of use in psychopharmacology as simulations of more complex psychiatric states and as screening tests in the development of novel treatments (Willner & Belzung, 2015). Various psychedelic compounds produce markedly similar effects in humans and non-human animals, demonstrating cross-tolerance, while sharing metabolic pathways and affecting common anatomical brain regions (Calvey & Howells, 2018; de Gregorio et al., 2021; Halberstadt, 2015). The 5-HT_2A_ receptor contributes to behavioral effects of psychedelics in humans and animals (Calvey & Howells, 2018; Glennon et al., 1984; López-Giménez, Juan F., González-Maeso, 2018). Pre-clinical studies demonstrating this effects include drug discrimination, tolerance, and behavioural assays such as head-twitch response (HTR), pre-pulse inhibition (PPI) of startle, interval timing, locomotor response and exploration s (de Gregorio et al., 2021; Halberstadt et al., 2017; Hanks & González-Maeso, 2013). Drug toxicity studies on non-human animals became legislated as a requirement for new drug development under food and drug regulatory frameworks in the late 1930s, and the requirement of animal research before human trials was written into both the Nuremberg code (1946) and the Helsinki Declaration (Van Norman, 2019).

Animal research has continued despite criticisms based on ethics (Barré-Sinoussi & Montagutelli, 2015), lack of translational success (Leenaars et al., 2019), bias towards anthropomorphism (Anderzhanova et al., 2017) and methodological concerns such as lack of replicability and inter-species variability (Herzog et al., 2018) as well as the limitations implicit in generalizing results from the adolescent-aged, in-bred rodent strains standard to animal investigations (Duque et al., 2021). Living in controlled experimental conditions results in chronic stress on study animals with resultant effects on neuromodulators and self-regulatory systems (Alexander et al., 1981).

Animal studies are inconsistent predictors of pharmacological toxicity in humans (Van Norman, 2019). Animal models of complex, multi-factorial psychiatric conditions such as depression, anxiety or substance use disorder remain a challenge limited by significant differences in language capacity, perceptual and sensory systems (Hanks & González-Maeso, 2013). Interpreting any single animal behaviour as reflective of human depressive-like symptoms lacks constructive validity, especially without the communication required to understand internal states of feeling (Commons et al., 2017; Slattery & Cryan, 2012). There is yet no clear “gold-standard” protocol for investigating the behavioral effects of psychedelics in non-human animals and the value of translational models of animal research remains somewhat controversial (De Gregorio et al., 2018).

Animal paradigms may be useful in that they can elicit observable, biological phenomenon of organisms not entirely dissimilar from humans and identify areas of possible consistency between animal and human behaviour. Animal research, particularly in areas of neuropsychology, can provide bottom-up information for the identification of underlying mechanisms and are useful in stimulating new conceptual paradigms. Similarly, animal paradigms can be of value in identifying the pharmacological effects of psilocybin outside of the psychological supports included in human trials (Meinhardt et al., 2020).

Generalizing results from animal to human trials is problematic. Animal research does not in itself create a pipeline of new drug discovery for clinical application, but does help provoke new understanding of mechanisms, as has been the case with neuroplasticity literature (Macpherson & Hikida, 2019). The study of the molecular biology of relatively simple *Aplysia* sea slug contributed to contemporary understandings of neuronal plasticity and implicit memory (Kandel et al., 2014).

### 1.3 Challenges in Knowledge Translation from Non-human Animal Studies

As translational research, animal models are assessed for their validity in application to humans by way of *construct validity*, *face validity* and *predictive validity* (Willner, 1984). Construct validity is met if there is coherence in the concept being tested, face validity requires the identification of common or similar underlying physiological mechanisms between the two, and predictive validity is the capacity to predict findings (Herzog et al., 2018; McOmish et al., 2014). Building on Wilner’s three primary criteria for translational validity, Belzung and Lemoine present a more nuanced and comprehensive framework for quality of validity assessment (Belzung & Lemoine, 2011). While there is partial overlap of criteria, the latter present five categories of validity in comparison to Willner’s two, disentangling different dimensions while redefining face and predictive validity: homological, pathogenic, mechanistic validity, face and predictive validity.

Quality standards for the conduct and reporting of scientific animal studies have been developed. The ARRIVE guidelines, Animal Research: Reporting In Vivo Experiments, were published in 2010 (Physiol, 2010), identifying the minimum standards of reporting necessary to transparently describe in vivo experiments. Revised in 2020, the guidelines have seen limited uptake, and many animal model publications fail to report on the information necessary to interpret the validity of the investigation (du Sert et al., 2020; Smith, 2020). A validated Risk of Bias (RoB) assessment tool for animal intervention studies (the SYRCLE’s RoB tool) has been developed (Hooijmans et al., 2014). Based on the Cochrane RoB tool, the SYRCLE RoB tool has been adjusted to the differences between randomized controlled human trials and animal studies. The tool assesses the risk for biases such as selection bias, performance bias, detection bias, attrition and reporting biases.

While it may be difficult to fully model a complex human disorder such as depression in animal models, diagnostic disorders and their distinctive symptomologies or traits can themselves be understood to have associated endophenotypes. Endophenotypes are measurable, trait-related deficits which can be assessed by laboratory study (Braff et al., 2007). Major depressive disorder, for example, is characterized by several distinct endophenotypes such as amotivation, anhedonia, and impaired cognition (Higgins et al., 2021). Behavioural assays such as exploratory and approach-avoidance conflict paradigms, forced swim test, tail suspension test and associative learning paradigms have been developed in animal research to capture elements of human psychiatric conditions (Teixeira & Quevedo, 2013; Willner & Belzung, 2015).

Animal studies which model such endophenotypes may have value in translation to more complex human dynamics as contributing to our understanding of the composite characteristics of mental health diagnoses (Slattery & Cryan, 2012). Phenotype measures have the additional value of being more proximal to actual genetic expression, physiology, and behaviour than the more abstract structures of mental health diagnostic criteria (Iacono, 2018) and may be more translatable across the preclinical to clinical spectrum with better predictive and construct validity (Day et al., 2008; Markou et al., 2009). While animal investigations may not be able to completely model a pathology, they can simulate certain aspects as reflected in their behavioural endpoints (Willner & Belzung, 2015). To improve validity, animal investigations have increasingly adopted domain-based inclusion criteria and moved away from the adoption of disorder-based nosology (Herzog et al., 2018).

### 1.4 The Research Domains Criteria Framework of Translational Research

The Research Domain Criteria (RDoC) is a framework for endophenotype-oriented translational neuroscience and presents a new framework for categorizing psychopathologies based on dimensions of observable behavior and neurobiological measures (Cuthbert & Insel, 2013). The RDoC matrix provides a dimensional and trans-diagnostic epistemic framework by which to understand pre-clinical and clinical sciences in the service of knowledge translation. In doing so, it provides an alternative nosology for the understanding of mental health, presenting as a distinct, more nuanced, and biologically valid framework than the definitions and categorizations of mental and emotional health as described in the DSM-V, the Diagnostic and Statistical Manual of the American Psychiatric Association. Under the RDoC framework, traditional psychiatric symptoms are understood as functions of a domain of biological activity, and dysfunctions can span both multiple domains and multiple disorders (Macpherson & Hikida, 2019).

The RDoC framework has recently been applied to psychedelic-assisted psychotherapy in a review of transdiagnostic effects (Kelly et al., 2021). By its focus on smaller units of analysis, RDoC proposes an interdisciplinary science of psychopathologies, identifying basic mechanisms which cut across diagnostic boundaries (Zoellner & Foa, 2016). The domains of analysis within RDoC serve as a matrix for model validation; homologies between human and experimental non-human animals in the domains justifies the validity, reliably and translatability of animal models appearing as *endophenotypes* of negative and positive affect, social interaction and general arousal/modulatory systems, while the complexity of the RDoC cognitive behavioural domain requires ongoing clarification (Anderzhanova et al., 2017).

### 1.5 Research Question

This scoping review was guided by a distinct research question: What are the observable neurological and behavioural effects of psilocybin in non-human animals? The scope of this study was limited to behavioural investigations and the observable effects of psilocybin s. From these studies, pharmacological effects of psilocybin can be mapped, and the studies were scoped to identify variables found to influence drug effects (such as pre and post conditioning manipulations). Within the larger scope of this research into the clinical potential of psilocybin, findings from the non-human animal research could complement and add additional insight to findings from human clinical trials – in part because they are not bound by the specificities of mental health diagnostics and clinical trial design.

## 2. METHODS

Scoping review methodology was selected to identify, summarize and map the literature on pre-clinical behavioural studies of non-human animals using psilocybin, psilocin and *Psilocybe* mushrooms. Scoping reviews utilize a rigorous, transparent and replicable methodology to comprehensively explore, identify and analyze all relevant literature pertaining to a research area (Arksey & O’Malley, 2005; Pham et al., 2014). Scoping reviews provide a basis in evidence for the evolution of both research and clinical practice, in this case by understanding the behavioural outcomes in translational models of animal research.

The scoping review was guided by an *a priori* protocol (see **Appendix** A). The objective was: To determine the effects of psilocybin (PSI) in animal studies across behavioural task clusters and neurological measures and to chart: what studies have been done, what behavioural tests have been used, what neurological measures have been implemented and which dosing modalities have been used and to what effect. Through librarian-conducted literature searches (see **Appendix** B) and defined inclusion criteria, the relevant number of internationally published, peer-reviewed academic articles in addition to grey area literature was identified, and the findings charted to identify the current state of psilocybin animal study research.

As a preliminary stage, we completed multiple preliminary reviews, literature research, scanned abstracts for keywords, identified pilot inclusion criteria, developed a pilot data extraction tool, and conducted background research into the history of psilocybin in animal models prior to conducting the actual scoping review. The study protocol was published in QSpace (https://qspace.library.queensu.ca/), an open access repository for scholarship and research produced at Queen’s University, under School of Kinesiology and Health Studies Faculty Publications. Reporting has been completed in accordance with the PRISMA-ScR Reporting Guidelines.

### 2.1 Search Strategy

A comprehensive search approach was employed by an experienced health services librarian to locate published studies and conference materials. A preliminary search was conducted in Ovid Embase using a combination of keywords and subject headings, followed by an analysis of relevant citations to identify other relevant keywords and subject headings. The optimized Ovid Embase search strategy was then adapted for Ovid MEDLINE, Ovid EBM Reviews – Cochrane Central Register of Controlled Trials, Ovid PsycINFO, Web of Science Core Collection, and BIOSIS Previews. All databases were searched from inception up to October 2019, with a search update performed in October 2021. No publication date limits were applied. The complete search strategies for all databases are presented in **Appendix** B. The reference lists of all eligible studies were screened to identify any additional studies. A librarian-conducted literature search was performed through MEDLINE, Embase and PsycINFO, using predefined key terms and search criteria (SM – Queen’s Bracken Health Sciences Library). Additional searching was conducted by reviewers, using electronic databases, grey literature sources, and reference lists of relevant articles or reviews. The first search was conducted in Fall of 2019, the second in the Spring of 2021 and finally October 2021.

### 2.2 Citation Management

All citations were imported into Covidence, duplicate citations were removed.

### 2.3 Eligibility Criteria

Studies were included which reported behavioural parameters in an animal model following the administration of psilocybin, psilocin or whole mushroom/whole mushroom extracts (WME). Only included primary research investigating behavioural effects in mammals was included in the scope of this review.

We excluded review articles (meta-analyses, etc.), but referred to their references to scope for primary research that may fit the inclusion criteria. Duplicates and studies that were not reported in English were excluded. We excluded publications that did not have an adequate control group, to which changes in behaviour could be compared. These included studies that only investigated interactive effects of PSI treatment and another compound (for example, in a competitive binding paradigm) or drug discrimination studies (where the behaviour is dependent on reinforcement trained by a separate compound), as we wanted to investigate the effects of PSI treatment alone on behaviour. Multiple studies investigated the effects of multiple drugs, so we included those studies if there was an experimental paradigm that included a control and reporting of behavioural outcomes following psilocybin administration without other pharmacological interventions.

### 2.4 Study Selection and Screening

Articles were included through a two-pass review system. In the first pass, two reviewers evaluated the abstracts in Covidence for compliance with basic inclusion criteria. Each reviewer opted to a) exclude the study as it was already evident it did not meet at least one of the predetermined criteria (NO), b) push the study forward as it remained unclear whether or not all criteria were met and the article required further review – or perhaps not enough information was available to make a judgement (MAYBE), or c) accept the study, to pass through to the full read (YES). The second pass included full-paper scanning to ensure all inclusion criteria were met.

### 2.5 Data Extraction

The included studies were divided between reviewers (RS, KD, NB, NT, KR) with reviewers extracting relevant information to a shared extraction spreadsheet. Data extraction was corroborated by a second reviewer, and a third reviewer went through all final exactions to ensure accuracy. Information in the summary table included article characteristics (year of publication, country of institutions), details of the study design and animal model used (number of experimental groups, type of experimental groups, total number of animals included; strain, sex, age, weight, and housing conditions of animals), dosing regimen (specific substance, route of administration, number and concentration of doses, control substance, time between drug administration and testing), behavioural test administered (including type of test, which behavioural parameters were recorded, setting of tests, training period), and outcomes (behaviour results, other mechanisms recorded, adverse outcomes). Following data extraction, the main outcomes were grouped by behavioural category.

### 2.6 Data Summary, Synthesis, and Charting

Given the broad scope of the review, outcomes were grouped by the type of behavioural test administered and clustered under the construct domains of the Research Domains Criteria (RDoC) of the National Institute of Mental Health. Mapping outcomes depended on the type of behavioural test used; results were further categorized by parameters that may contribute to behavioural outcome, including dose or time between administration and testing. To provide dose equivalencies, the 0.71:1 equimolar psilocin to psilocin conversion is used; the predominantly used compound is used as the base comparison in each table. This does not account for differences that may result from metabolism, or other variables, but serves as a rough comparison across different study drug formulations.

From records identified through database screening and using Covidence online software, duplications were removed, records screened, assessed for eligibility and included or excluded from full-text review (**Figure** 3.1) Reviewers (RS, KD, NB, NT, KR) read the full text of selected studies and each extracted data using an *a priori* data extraction spreadsheet. Reviewers cross-checked for homogenous process quality control and consensus was necessary when conflicts arose.

**Figure 3.1:**
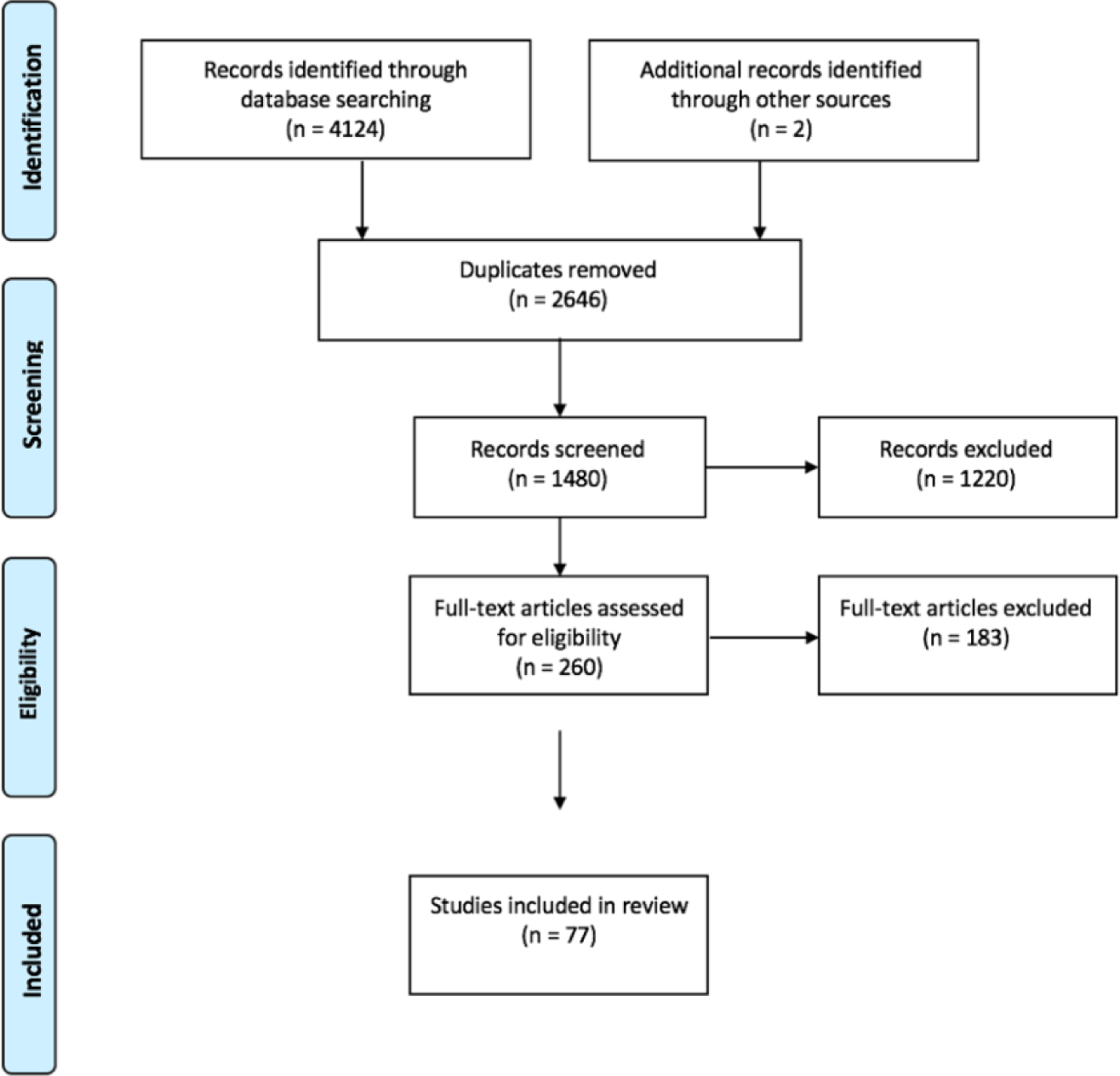
Prisma Flow Diagram, BIPA Scoping Review (p.12)

### 2.7 Classification of Results by RDoC Matrix

Results were allocated across the RDoC framework (see **Table** 2.1). Our team assigned the various individual animal assays to one of the domains after a thorough review of each study design, consultation with the larger body of literature specific to the paradigms (Angoa-Pérez et al., 2013; Commons et al., 2017; Kalueff et al., 2016; Kelly et al., 2021; Pittenger et al., 2019; Scheggi et al., 2018; Slattery & Cryan, 2012; Willner & Belzung, 2015) and by consensus within the team. Each of the seventy-seven individual publications was then located within the domain classifications. Using the RDoC framework in this way allowed for the presentation of our scoping review results within an established and valid epistemic framework of health research and knowledge translation.

**Table 2.1:**
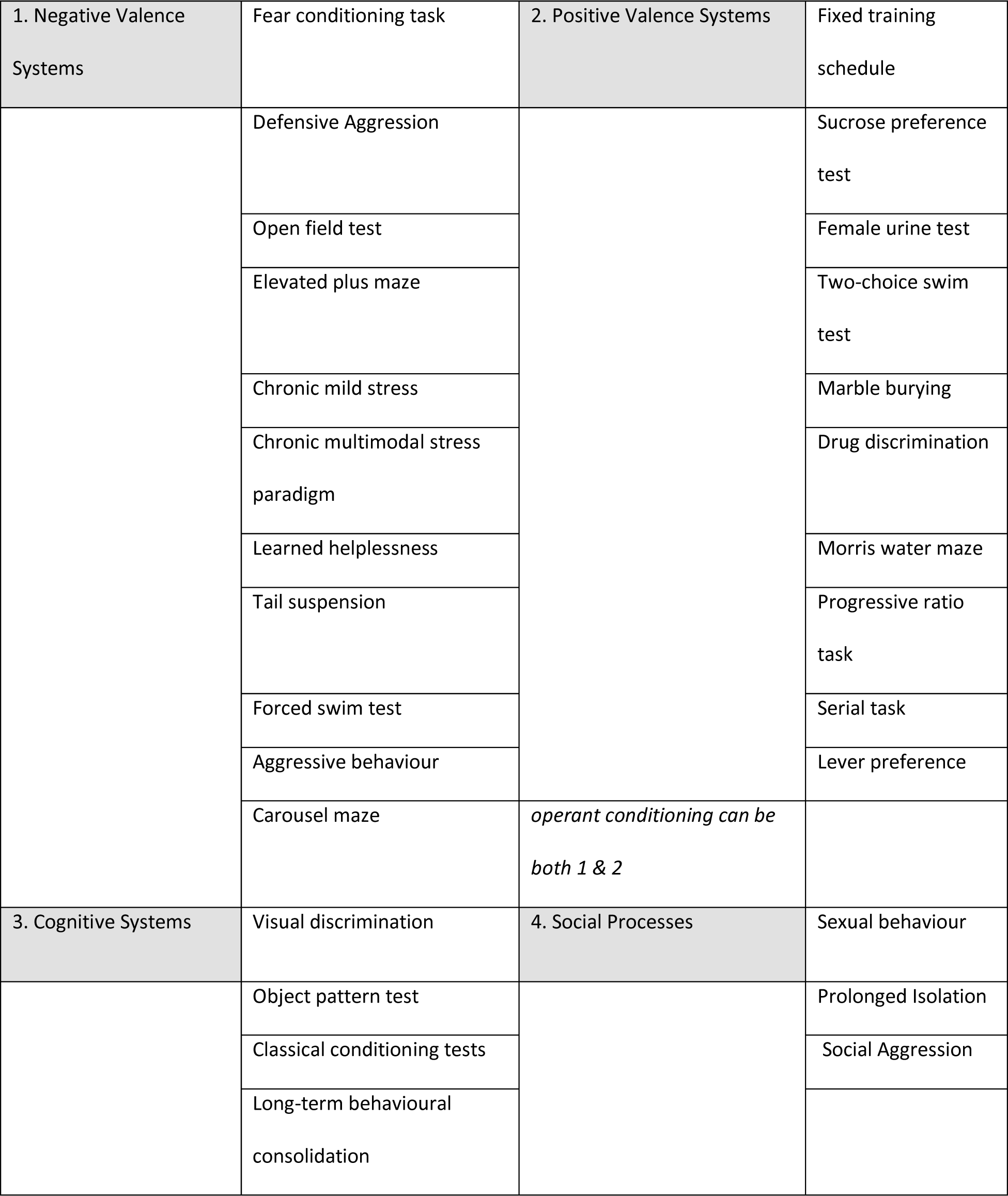
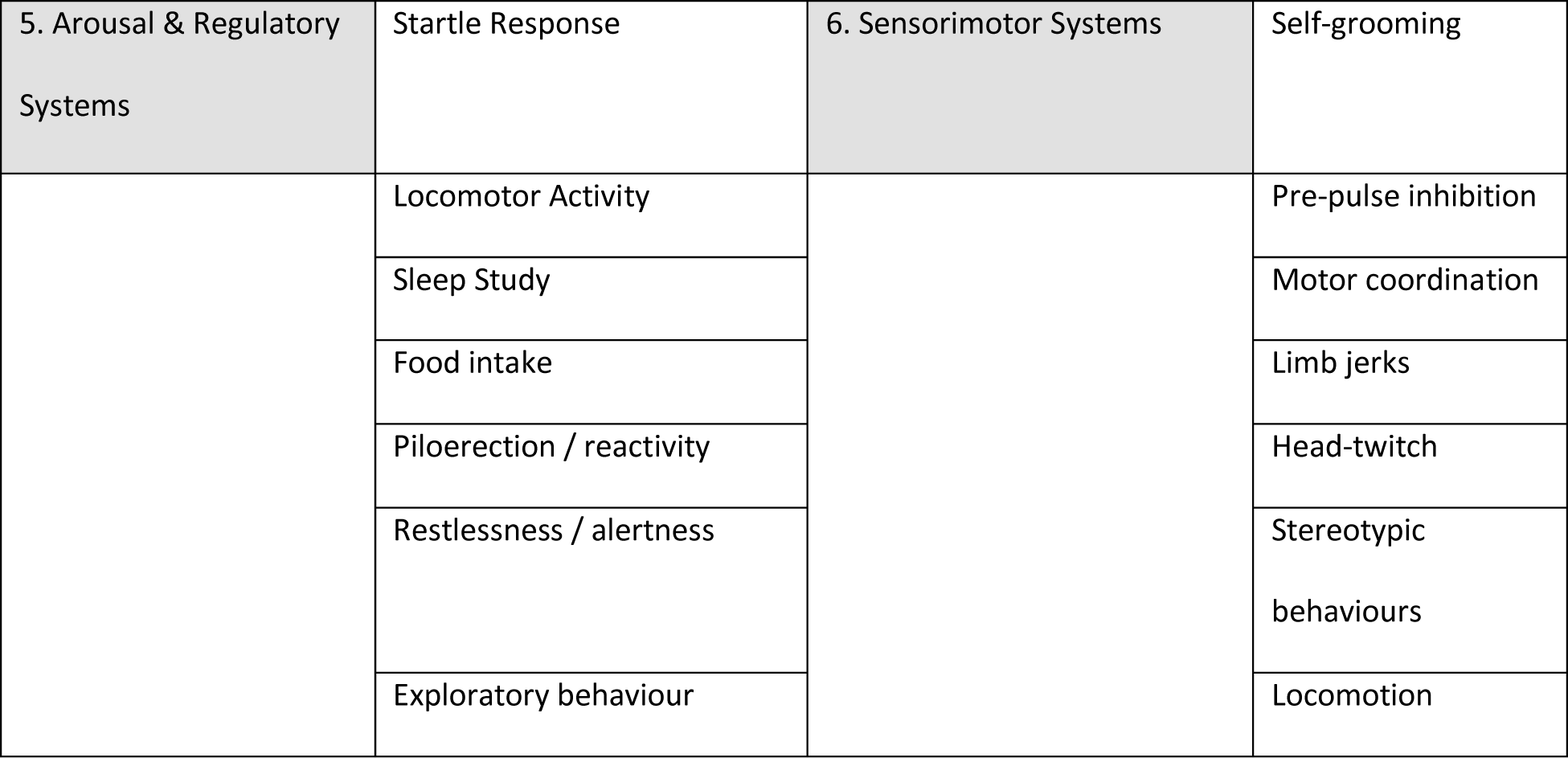
Animal Paradigms by RDoC Domain Classification (p.13)

The development of this table is a novel and significant contribution to knowledge translation. By assigning known animal paradigms to the six domains of the RDoC framework, we were able to group or cluster the results of the individual studies within specific process-oriented higher-order domains of function. Study results were mapped directly onto this epistemological framework as part of an overall scoping of psilocybin pre-clinical health research using animal models. This adds to the translation value of this scoping review by mapping results onto this recognized and validated framework. Applying the RDoC criteria in this manner is intended to strengthen the interpretation and presentation of findings, increasing validity by situating the study findings within more specific, localized data points and providing an epistemic framework for the interpretation of study results. In this way, the diagnostic validity of DSM-V or ICD-10 definitions of such things as depression are less relevant, and study findings can more meaningfully be nested within areas of behavioural function and known biological systems.

## 3. RESULTS

Seventy-seven publications met eligibility criteria and are included in this scoping review. From 4124 records identified through database searching, 2646 duplicate records were removed and two additional hand-sorted studies meeting eligibility criteria were added. From 1480 records screened, 1220 records were excluded for not meeting eligibility criteria established in our a priori protocol (see **Appendix** A). 260 articles were subject to full-text review, and finally 77 studies were included in this scoping review (**Figure** 3.1).

### 3.1 Study Characteristics

Publications are primarily scientific journal articles (n = 63), but given the wide purview of scoping reviews, some conference abstracts are also included (n = 14) (**Table** 3.1), including abstracts which may then have been followed by full publications on the same research. All included studies investigate the behavioural effects of psilocybin in non-human animals, and all investigations were with non-human mammals. This review captures research published over six decades between 1962 and 2021. Forty-six studies were published in or before 1987, thirty-one studies were published in 2005 or later, and interestingly, no study publications were found for the time-period 1988 to 2004 (**Figure** 3.2). The decade with the most publications was the 1970s. Institutional affiliations of Primary Investigators span 14 different countries, with more than half of the studies based in the United States (n = 43) (**Table** 3.1).

**Table 3.1:** Characteristics of Included Studies: BIPA (p.14)

| Publication Information | Number (n<br>= 77) | Percentage |
| --- | --- | --- |
| **Publication Type** |  |  |
| Journal articles | 63 | 81.8% |
| Conference abstracts | 14 | 18.2% |
| **Publication Year by Decade** |  |  |
| 1961 – 1970 | 11 | 14.3% |
| 1971 – 1980 | 25 | 32.5% |
| 1981 – 1990 | 10 | 13.0% |
| 1991 – 2000 | - | - |
| 2001 – 2010 | 3 | 3.9% |
| 2011 – 2020 | 22 | 28.6% |
| 2021 | 6 | 7.8% |
| **Country of Author Affiliation\*** |  |  |
| United States | 43 | 55.8% |
| United Kingdom | 11 | 14.3% |
| Czech Republic | 7 | 9.1% |
| Brazil | 3 | 3.9% |
| Canada | 3 | 3.9% |
| Denmark | 3 | 3.9% |
| Japan | 3 | 3.9% |
| Poland | 2 | 2.6% |
| Sweden | 2 | 2.6% |
| Other** | 5 | 6.5% |
| **Species\*** |  |  |
| Non-human primates | 9 | 11.7% |
| Pigs | 1 | 1.3% |
| Dogs | 2 | 2.6% |
| Cats | 5 | 6.5% |
| Rabbits | 2 | 2.6% |
| Guinea pigs | 2 | 2.6% |
| Rats | 41 | 53.2% |
| Mice | 19 | 24.7% |
| **PSI Treatment Formulation\*** |  |  |
| Psilocin | 33 | 42.9% |
| Psilocybin | 51 | 66.2% |
| Whole mushroom extract | 4 | 5.2% |
| **Sex of Research Animals** |  |  |
| Male | 40 | 52.0% |
| Female | 13 | 16.9% |
| Both | 11 | 14.3% |
| Not reported | 13 | 16.9% |

\* Values represent the number of studies that include the country affiliation, animal species, or treatment compound, with some studies including multiple affiliations, species, or treatments. Therefore, total number is greater than 77 and total percentage is greater than 100%.
\*\* Other reported countries of author affiliation, including those of co-authors, are France, Iran, Israel, Italy, and Senegal.

**Figure 3.2:**
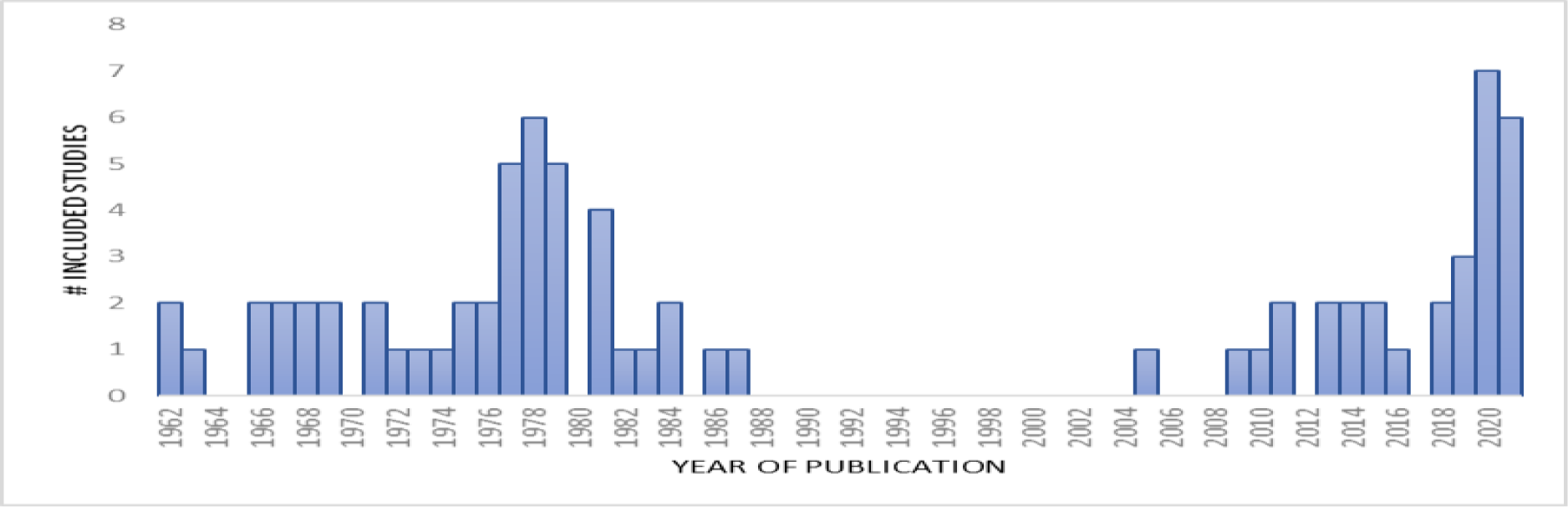
Year of Publication of Included Studies (p.14)

#### 3.2 Research Animals

The preponderance of studies investigate behavioural outcomes in rodents (see **Table** 3.1)., specifically rats (n = 41) or mice (n = 19) and four studies report behavioural outcomes in both rats and mice (Bourn et al., 1978, 1979; Kostowski et al., 1972; Thompson, 2019). Fewer studies use non-human primates (NHPs; n = 9) or other larger mammals, like cats (n = 5), dogs (n = 2), or pigs (n = 1). Rabbits (n = 2) and guinea pigs (n = 2) are also studied. Studies published in 2005 or later exclusively investigate the effects of Psi treatment in rats or mice, except for only one experiment with pigs (Donovan et al., 2020).

More than half of the included studies investigate the effects of Psi treatment in rats (n=41, or 53.2%). Various strains are used, including Sprague-Dawley (n = 14), Wistar (n = 12), Wistar-Kyoto (WKY) (n = 3), Long-Evans (n = 3), Flinders Sensitive (FSL) and Flinders Resistant Lines (FRL) (n = 1). Despite differential outcomes between rat strain performance, several studies (n = 8) failed to report strain. We report which strain was investigated when available.

Reporting of animal housing conditions provides context for the interpretation of behavioural outcomes and is an indication of study quality. Animal Research: Reporting of *In Vivo* Experiments (ARRIVE) guidelines (2010) outline essential information preferentially included in animal research publications to provide context for the interpretation of experimental outcomes and help determine reliability of study findings (du Sert et al., 2020). Evaluating all studies for their inclusion of essential ARRIVE information was beyond the scope of this review, and most were published prior to the ARRIVE guidelines; however, study inclusion of some relevant parameters (such as reporting of animal housing conditions) is noted by us as a proxy measure of methodological quality by modern standards. Behavioural responses of research animals are sensitive to differing environmental factors and to the housing conditions of the study. We noted where studies provide information about housing condition, length of animal training and reports of biological sex as well as the developmental age of animal. All are variables known to effect outcomes.

Only 43 of the included studies (55.8%) report any degree of housing conditions, describing the primary housing enclosure, food, or light schedule, or whether animals are housed individually or together in groups. Conference abstracts are also included in this review but typically do not report all study details. Two of fourteen (14.3%) conference abstracts report housing conditions, compared to forty-one of sixty-three journal articles (65.1%). The reporting of housing conditions in journal articles varies over time (see **Table** 3.2). 53.5% (23 / 43) of journal articles published from 1962 to 1987 report housing conditions, compared to 90.0% (18 / 20) of journal articles published from 2005 to 2021, reflecting a trend towards improved study quality over time.

**Table 3.2:** Housing Conditions Reported in Journal Articles, by Year of Publication (p.16)

| Date Range | Reported Housing Conditions | No Information |
| --- | --- | --- |
| 1961 – 1970 | 4 (36.4%) | 7 (63.6%) |
| 1971 – 1980 | 14 (58.3%) | 10 (41.7%) |
| 1981 – 1990 | 5 (62.5%) | 3 (37.5%) |
| 1991 – 2000 | - | - |
| 2001 – 2010 | 2 (100%) | 0 (0%) |
| 2011 – 2020 | 10 (83.3%) | 2 (16.7%) |
| 2021 | 6 (100%) | 0 (0%) |
| **Total** | **41 (65.1%)** | **22 (34.9%)** |

*The number of included journal articles that report housing conditions or provide no information, reported with percentage per years of publication.*

Sixty studies (77.9%) report the number of non-human animals included in experiments, leaving seventeen studies (22.1%) failing to report study sample size. Of the publications that do not report study sample size, eight are peer-reviewed academic journal articles and nine are conference abstract presentations. The number of non-human animals included in each study varies from as few as two rats (Cameron & Appel, 1976) to groups of twenty per treatment condition (Collins et al., 1966)

#### 3.3 Trial Characteristics and Experiment Design

All studies investigate behavioural outcomes following drug administration: fifty-one studies (66.2%) use psilocybin, thirty studies (42.9%) administer its active metabolite psilocin (42.9%), and four studies (5.2%) administer extracts from whole mushrooms containing both psilocybin and psilocin as well as other active alkaloids which together may create a synergistic entourage effect (**Table** 3.1). Eight studies investigate the effects of both psilocin and psilocybin, one study reports the effects of both psilocin and whole mushroom extracts (WME) (Zhuk et al., 2015) and one study investigates behavioural outcomes following administration of psilocin, psilocybin and WME (Matsushima et al., 2009). Four studies investigate the alkaloid composition, toxicity, and behavioural effects of full mushroom extracts. Extracts are prepared from *Psilocybe cubensis* (Kirsten & Bernardi, 2010; Mahmoudi et al., 2018), *Psilocybe argentipes* (Matsushima et al., 2009), *Psilocybe semilanceata* and *Pholiotina cyanopus* species (Zhuk et al., 2015).

The most common route of administration for Psi treatment is intraperitoneal injection (n = 49) (see **Table** 3.3). Intraperitoneal injection (IP) is the most common method of administration for rats (n = 26), mice (n = 15), and cats (n = 4). Subcutaneous injections (SC) are also sometimes used in rats (n = 9). Intravenous injections (IV) were used in non-human primates (n = 3), rabbits (n = 3), and pigs (n = 1). Two studies administer Psi orally to mice (Collins et al., 1966; Matsushima et al., 2009). Two studies administer Psi via intramuscular injection (IM) to non-human primates (Schlemmer & Davis, 1986; Sink et al., 1983) and in one study, psilocybin was administered intraventricularly to macaque monkeys (Wada, 1962).

Experimental doses used range from 0.01 to greater than 100 mg/kg psilocybin for a single administration. Most included studies (58 / 77; 75.3%) investigate the effects of more than one treatment dose (**Table** 3.3). Many studies report dose-dependent as well as time-dependent effects of Psi treatment; both effects are significant and are separately reported below. Despite the range of treatment doses investigated, most studies include doses that are equal to or less than 1.0 mg/kg psilocybin (n = 56). Some studies (n=17) only include doses greater than 1 mg/kg. The translational value of these doses may be less clear as they far exceed doses tested in clinical or research use in humans. However, administration of these exceedingly high doses -- 100 mg/kg psilocybin (Horita & Weber, 1962) and 120 mg/kg psilocybin (Schneider, 1968) – occurs with no reported biological adverse effects, providing evidence toward the strong biological safety profile of Psi.

**Table 3.3:** Characteristics of Psilocybin Treatment (p.17)

| Dosing Information | Number (n<br>= 77) | Percentage |
| --- | --- | --- |
| **Route of Administration\*** |  |  |
| Intraperitoneal injection | 49 | 63.6% |
| Subcutaneous injection | 11 | 14.3% |
| Intravenous injection | 7 | 9.1% |
| Intramuscular injection | 2 | 2.6% |
| Oral administration | 2 | 2.6% |
| Intraventricular injection | 1 | 1.3% |
| Not reported | 7 | 9.1% |
| **Number of Doses** |  |  |
| Only 1 | 19 | 24.7% |
| More than 1 | 58 | 75.3% |

*\* Values represent the number of studies that use the type of administration method, with some studies including multiple methods (thus cumulative percentage is greater than 100%).*

Studies vary widley in methodology for investigation of drug effect on behavioural outcome, in the number or frequency of dosing, and/or in the timing between treatment administration and behavioural measures. Some studies investigate the acute effects of a single Psi treatment and assess behaviour within the minutes-to-hours following dose administration, time-points to the understanding of the various phases of acute drug effect. Other studies investigate the persisting effects of single or repeated psilocybin doses, with behavioural assessments occurring weeks after drug administration. Some studies administered psilocybin repeatedly on dosing regimens reported as “microdosing”.

Some studies investigate the effects of multiple compounds, comparing the effects of psilocybin with other serotonergic hallucinogens or stimulants; with these we report only the behavioural outcomes specific to psilocybin. In some studies, the effects of multiple compounds were tested in a single subject, with reported wash-out periods or intervals between drug administration ranging from 3 days (Harris et al., 1981) to 2 weeks (Roberts & Bradley, 1967). Since cross-tolerance has been demonstrated across various serotonergic psychedelics, in some instances there may be interference in behavioural measures when study drugs are administered within shorter wash-out periods.

Of the studies which investigate acute behavioural effects, there is large variability across studies in the time between Psi administration and behavioural testing. Behavioural testing may begin immediately following injection or up to 90 minutes following administration (Kostowski et al., 1972). The duration of behavioural measurement varies as well; some assays have a four-minute testing period (as with FST, TST) while many measure behaviours across a longer time-course of acute drug behavioural effect.

Studies used a variety of experimental controls. Some employed separate control groups while others used the same animal as its own control. This is of note due to evidence of persisting effects for up to one-month and the need for sufficient wash-out periods between doses. The behavioural effects are documented along a time course of drug effect, but it is not entirely clear when an animal has entirely shed the downstream secondary effects of drug metabolism within this period lasting weeks. In experiments using the same animal as their own control, the time between active and sham drug administration varied. Careful consideration should be given to interpretation of these experiments. Further, most studies investigated only one test of behavior. In assessing trial quality, the assessment of multiple behavioural reflects positively; multiple assessments provide more and richer information, allowing the ability to compare, contrast information, and to better identify potential underlying mechanisms of action while giving a better behavioural profile of the animal.

#### 3.4 Sex as a Biological Variable

Forty studies (52%) included in the scope of investigate the effects of psilocybin treatment on behaviour in only males, while 24 studies include females (31.2%) including studies with only females and studies with both sexes (**Table** 3.1). Thirteen studies (16.5%) fail to report the sex of research animals included in experimentation. Of the 11 studies that investigate both sexes, five studies did not report behavioural outcomes separately per sex (Martin et al., 1978; Roberts & Bradley, 1967; Steiner & Sulman, 1963; Trulson et al., 1984; Vaupel et al., 1979) and one did not incorporate sex as a factor in the analysis of behavioural outcomes (Everitt & Fuxe, 1977). Four studies (5.2%) incorporate sex as a variable in study design and analysis, and of those, two studies report sex-specific differences in behavioural outcomes.

Halberstadt et al. report no significant effect of sex on head-twitch response in mice following treatment with psilocin and report no significant interaction between sex and other variables or behavioural outcomes, collapsing data across sexes (Halberstadt et al., 2011). Conversely, mice treated with whole mushroom extract had significant sex differences in wet-dog shakes, gnawing behaviour and locomotion (Kirsten & Bernardi, 2010). Tylš et al. report a significant effect of sex on locomotor activity, rearing behaviour, and wet-dog shakes; female rats appear less affected by psilocybin than male rats except for wet-dog shakes (Tylš et al., 2016). Shao reports more pronounced increased spine formation rate and density among female rats compared to males because of Psi but does not separately report behavioural outcomes (Shao et al., 2021). Kirsten & Bernardi report sex differences in self-grooming behaviour after psilocybin, with increases in self-grooming noted among females after dosing and reduced self-grooming among males compared to controls (Kirsten & Bernardi, 2010). One study separates female rats into separate pre-estrus, estrus, and metaestrus, and diestrus groups (Tylš et al., 2016).

#### 3.4 Behavioural Research Domains

We report behavioural outcomes under the six domain constructs of the Research Domain Criteria (RDoC) framework (**Table** 3.4). Within each domain, experimental outcomes are reported by the test administered. Where trial numbers justify, we further grouped and summarized results by sub-construct or behavioural assay. Sensorimotor Systems is the RDoC domain most studied when measured by the number of distinct behavioural tests reported (n=47), followed by Negative Valence (n=34), Arousal and Regulatory Systems (n=26), Positive Valence (n=14), Social Processes (n=6) and Cognitive Systems (n=3). One-hundred and thirty distinct behavioural investigations into the effects of psilocybin in non-human animals are captured in these seventy-seven studies, and we identified fifty different behavioural paradigms used in this body of research.

**Table 3.4.:** Animal Behavioural Paradigms by RDoC Framework Domains (p.21)

|  | Overview | Included Studies |
| --- | --- | --- |
| 1. Negative Valence Systems | N=34 |  |
| Fear Conditioning | N = 11<br>Low doses of Psi decrease response latency, perhaps indicating heightened sensitivity.<br>Doses of 5 mg/kg demonstrate dynamic time-dependent, biphasic effects with a corresponding decrease then increase in avoidance proficiency. | Bourn et al., 1978<br>Bourn et al., 1979<br>Calil, 1978<br>Catlow et al., 2013<br>Collins et al., 1966<br>Davis & Hatoum, 1987<br>Hagsater et al., 2021<br>Marquis et al., 1973<br>Rambousek et al., 2014<br>Shao et al., 2021<br>Sugrue, 1969 |
| Forced Swim Test (FST) | N = 6<br>Three studies show no effect of Psi on FST behavior, and three studies. report decreased time spent immobile following treatment (indicating less despair). Peak behavioural effect noted at 35 days post Psi. Persisting, more than acute effects, demonstrated. Psi reverses effects of chronic stress. | Hesselgrave et al., 2021<br>Hibicke et al., 2019<br>Hibicke et al., 2020<br>Jefsen et al., 2019<br>Mahmoudi et al., 2018<br>Nichols & Hibicke, 2020 |
| Tail Suspension Test (TST) | N = 1 No effect noted. | Mahmoudi et al., 2018 |
| Defensive Aggression | N = 2<br>Both studies demonstrate dose-dependent inhibition of aggressive behaviour, with strongest effect 30 minutes post Psi administration. | Sbordone et al., 1979<br>Uyeno, 1967 |
| Elevated Plus Maze (EPM) | N = 3<br>Increased time in open arms at 2 and at 41 days.<br>Microdoses elicited a mild increased entry to and time spent open arms of maze. | Horsley et al., 2018<br>Hibicke et al., 2019<br>Hibicke et al., 2020 |
| Open Field Test (OFT) | N = 11<br>Time, dose, and sex-dependent effects on behavior.<br>Decreased locomotion, decreased time in OF. | Collins et al., 1966<br>Cunha & Masur, 1978<br>Hesselgrave et al., 2021<br>Hibicke et al., 2019<br>Jefsen et al., 2010<br>Kirsten & Bernardi, 2010<br>Mahmoudi et al., 2018<br>Palenicek et al., 2005<br>Tyls et al., 2011<br>Tyls et al., 2014<br>Tyls et al., 2016 |
| 2. Positive Valence Systems<br>Five-choice task, binge-like feeding, drug as discriminative stimulus, drug-induced conditioned suppression, female urine sniffing test, intra-cranial self-stimulation, Y-water maze, lever preference, marble-burying task, Morris water maze, progressive ratio task, serial task, sucrose preference test, two-choice swim test | N=14<br>Psi reduces binge-eating behaviour at several doses. Psi inhibits compulsive marble-burying behaviour throughout in a dose-dependent manner, with most effect evident at the medium-high but not highest dose range. Psi exerts rapid antianhedonic actions in chronically stressed mice.<br>Psilocybin demonstrates a biphasic dose and time response, with decreases in behavioural response evidenced early followed by subsequent increases in hedonic | Cameron & Appel, 1976<br>Castellano, 1978<br>Harris et al., 1981<br>Hesselgrave et al., 2021<br>Hesselgrave et al., 2021b<br>Higgins et al., 2021<br>Hurley et al., 2020<br>Koerner & Appel, 1982<br>Marquis et al., 1973<br>Matsushima et al., 2009<br>Rambousek et al., 2014<br>Rech et al., 1975<br>Sakloth et al., 2019<br>Thompson, 2019 |
|  | behaviours. At lower doses Psi demonstrates discriminative properties with fewer grossly disruptive effects on behaviour |  |
| 3. Cognitive Systems<br>Visual discrimination tasks, object pattern separation | N = 3<br>Psi treatment dose-dependently disrupts size discrimination and visual accuracy. Psi's persisting effects rescue chronic stress-induced visual system impairments. | Nichols & Hibicke, 2020<br>Roberts & Bradley, 1967<br>Uyeno, 1969 |
| 4. Systems for Social Processes<br>Female sexual receptivity, and social isolation-induced aggression paradigms | N = 6<br>Acute decrease (reduced social grooming, increased distancing) in affiliative social behaviour in NHPs.<br>Biphasic, dose-dependent effect on sexual receptivity among rats, with lower doses significantly increasing sexual receptivity and higher doses having a non-significant effect. Psi treatment is consistent in reducing aggressive behaviour, with greater inhibition of aggressive behaviour | Everitt & Fuxe, 1977<br>Schlemmer & Davis, 1986<br>Sink et al., 1983<br>Uyeno, 1966<br>Meldrum & Naquet, 1971<br>Kostowski et al., 1972 |
|  | as a function of increasing dose,<br>and with greater treatment effect<br>the longer the pre-testing isolation<br>period. |  |
| 5. Arousal and Regulatory Systems | N=26 |  |
| General Arousal and Physiological<br>Responses | N = 12<br>Acute changes in general arousal<br>are observed, accompanied by<br>mydriasis, piloerection, elevated<br>heart rate and respiratory rate, and<br>decreased appetite. | Guest & Consroe, 1979<br>Horibe, 1974<br>Horita & Weber, 1962<br>Martin et al., 1978<br>Meldrum & Naquet, 1971<br>Schlemmer & Davis, 1986<br>Sink et al., 1983<br>Steiner & Sulman, 1963<br>Sugrue, 1969<br>Vaupel et al., 1979<br>Wada, 1962<br>Wilson et al., 1981 |
| Sleep-Wake Behaviour | N = 1 | Thomas et al., 2020 |
|  | Psi is acutely wake-promoting, at the expense of REM sleep, with no persisting changes in sleep-wake behaviour observed. |  |
| Locomotor Activity | N=4<br>Mixed results. Among some animals and in some tests at lower doses Psi increased total activity while reducing in other tests. Evidence of reduced locomotor activities at higher doses, and some stimulation of activity at lower doses. | Bert et al., 1968<br>Bourn et al., 1978<br>Bourn et al., 1979<br>Collins et al., 1966 |
| Startle Response | N = 2<br>Dose-dependent effect of Psi on startle response. Low doses increase startle amplitude or have no effect while higher doses depress startle response | Davis & Walters, 1977<br>Geyer et al., 1978 |
| Investigatory/ Exploratory Behaviour | N=7 | Geyer et al., 1979<br>Halberstadt et al., 2011<br>Jacobs et al., 1977(a)<br>Jacobs et al., 1977(b)<br>Sakashita et al., 2015<br>Trulson et al., 1981 |
|  |  | Trulson et al., 1984 |
| 6. Sensorimotor Systems | N=47 |  |
| Stereotyped Behaviour | N=27<br>Stereotyped behaviours such as head-twitch response are reliably induced in animals following Psi treatment, once the dose crosses a threshold of efficacy, and are potentially suppressed at higher doses. Extremely high doses also induce myoclonus or head rotation in a dose-dependent manner. | Donovan et al., 1977<br>Everitt & Fuxe, 1977<br>Guest & Consroe, 1979<br>Halberstadt et al., 2011<br>Hesselgrave et al., 2020<br>Hesselgrave et al., 2021<br>Higgins et al., 2021<br>Jacobs et al., 1976<br>Jacobs et al., 1977(a)<br>Jacobs et al., 1977(b)<br>Jenner et al., 1981<br>Kiilerich et al., 2019<br>Kirsten & Bernardi, 2010<br>Klein et al., 2021<br>Luscombe et al., 1984<br>Martin et al., 1978<br>Meldrum & Naquet, 1971<br>Sakashita et al., 2015<br>Schlemmer & Davis, 1986<br>Schneider, 1968<br>Shao et al., 2021<br>Sherwood et al., 2020 |
|  |  | Sink et al., 1983<br>Trulson et al., 1981<br>Trulson et al., 1984<br>Tyls et al., 2016<br>Zhuk et al., 2015 |
| Grooming | N = 10<br>Biphasic effects, with no reductions in grooming at lower doses but disruptions evident at standard doses and above. Effects on grooming may be related to the time-course of the drug effect Sex-dependent differences are reported in mice. | Cunha & Masur, 1978<br>Guest & Consroe, 1979<br>Higgins et al., 2021<br>Kiilerich et al., 2019<br>Kirsten & Bernardi, 2010<br>Schlemmer & Davis, 1986<br>Sink et al., 1983<br>Trulson et al., 1981<br>Trulson et al., 1984<br>Tyls et al., 2016 |
| Motor Coordination | N = 5<br>Dose-dependent effect on discoordination with no effect at lower doses and discoordination at very doses. | Bourn et al., 1978<br>Bourn et al., 1979<br>Horibe, 1974<br>Schneider, 1968<br>Trulson et al., 1977 |
| Prepulse Inhibition | N = 5<br>Psi effectively disrupts PPI in rats.<br>This effect may be dose-dependent and have sex-specific effects. 1 | Kiilerich et al., 2019<br>Palenicek et al., 2005<br>Tyls et al., 2011<br>Tyls et al., 2013 |
|  | mg/kg psilocin (IP) effectively disrupts PPI ASR in rats, but lower doses (0.25 mg/kg) have no effect and higher doses (4 mg/kg) have variable effects. | Tyls et al., 2016 |

#### 3.5 Adverse Events

Few adverse events were reported in the scope of this review. We identified seven studies as reporting adverse events (**Table** 3.5). In some cases, these events were communicated in passing and not identified as adverse events in the publication itself. Overall, across the seventy-seven publications and one hundred and thirty paradigms reported, the safety profile of psilocybin at a biological level appears exceedingly safe. As reported earlier, some of the physiological effects reported (increased heart rate, for example) could be considered adverse as could some of the stereotyped behaviours (such as head-twitch or limb flick). Many trials report sedation at high doses if that can be considered adverse. Of the seven trials reporting adverse effects, none are correlated with psilocybin drug-effect, but rather, related to animal housing conditions. Respiratory infections were reported, surgical infections, as well as one “premature” death with no reference to causation or relation to drug effect. Several studies mention deaths of study animals during pre-drug training schedules prior to Psi administration.

**Table 3.5.**
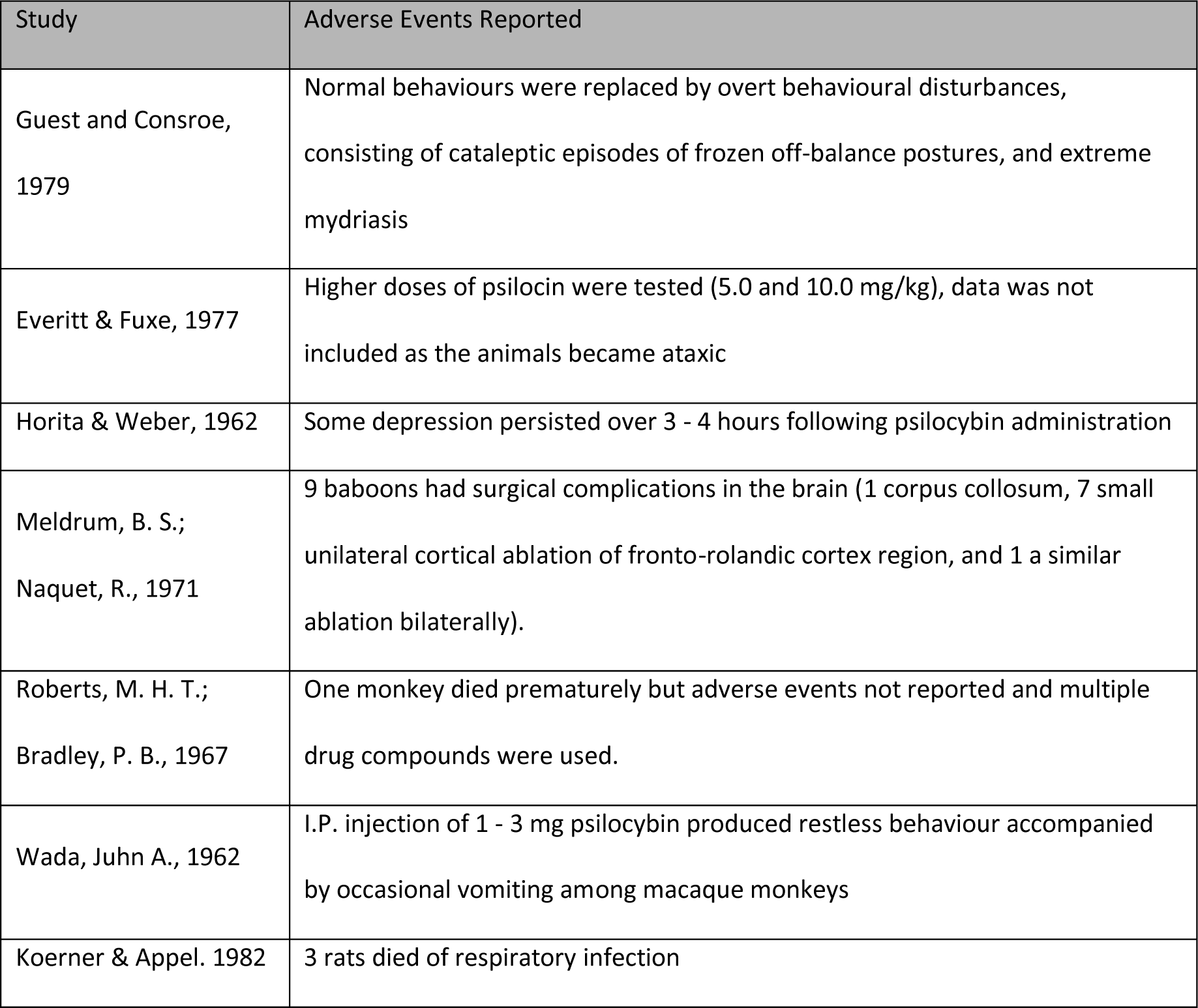
Studies Reporting Trial-related Adverse Events (p.21)

| Study | Adverse Events Reported |
| --- | --- |
| Guest and Consroe, 1979 | Normal behaviours were replaced by overt behavioural disturbances, consisting of cataleptic episodes of frozen off-balance postures, and extreme mydriasis |
| Everitt & Fuxe, 1977 | Higher doses of psilocin were tested (5.0 and 10.0 mg/kg), data was not included as the animals became ataxic |
| Horita & Weber, 1962 | Some depression persisted over 3 - 4 hours following psilocybin administration |
| Meldrum, B. S.;<br>Naquet, R., 1971 | 9 baboons had surgical complications in the brain (1 corpus callosum, 7 small unilateral cortical ablation of fronto-rolandic cortex region, and 1 a similar ablation bilaterally). |
| Roberts, M. H. T.;<br>Bradley, P. B., 1967 | One monkey died prematurely but adverse events not reported and multiple drug compounds were used. |
| Wada, Juhn A., 1962 | I.P. injection of 1 - 3 mg psilocybin produced restless behaviour accompanied by occasional vomiting among macaque monkeys |
| Koerner & Appel. 1982 | 3 rats died of respiratory infection |

Exceedingly high doses of psilocin, psilocybin and whole mushroom extracts were tolerated. No toxic effects were reported following intraperitoneal administration of 180 – 250 mg/kg psilocin in mice (Zhuk et al., 2015). However, the reported range of lethal effect is narrow, with the median lethal dose (LD_50_) of psilocin only a slightly higher concentration (293.07 ± 1.02 mg/kg). LD_50_ value for whole mushrooms extracts was greater than those of pure psilocin and the application of psilocin exhibited the highest toxicity compared to the extracts. Other investigations with high doses were tolerated. Subcutaneous administration of 80 – 120 mg/kg psilocybin to rats induced ataxia and unusual behavioural effects as evidenced by motor discoordination (Schneider, 1968). Intraperitoneal injection of 100 mg/kg psilocybin and 72 mg/kg psilocin in mice led to piloerection, exophthalmos, and hind-leg ataxia, with effects completely reversible and animals appearing completely normal within three to four hours following treatment, except for some depression of activity (Horita & Weber, 1962).

In evaluating the acute toxicity of an aqueous *Psilocybe cubensis* extract, increased gnawing behaviour and stereotyped shaking was reported, but no other harmful physical effects or mortality (Kirsten & Bernardi, 2010). Vomiting and restless behaviour was reported among macaque monkeys, adverse events common to humans. This study investigating electrocortical activity with concomitant electrical stimulation reported that intraperitoneal injection of psilocybin (1 – 3 mg) in macaque monkeys (weighing 5 – 6 kg) produced occasional vomiting (Wada, 1962).

Overall, few studies report adverse events following Psi treatment, even with exceedingly high doses. However, some doses were assessed in pilot studies and not used in the experiment as the doses impaired movement in such a way that would interfere with the behavioural task. Higher doses of psilocin (5 and 10 mg/kg) were not included in a study of investigatory behaviour in rats, as the animals became ataxic (Geyer et al., 1979). Dose-dependent responses were investigated in pigs (Donovan et al., 2021). Doses of 0.04 and 0.08 mg/kg psilocybin had an observable effect on behaviour, while 0.16 mg//kg led to the animal lying down with its eyes closed and not moving. The lower dose (0.08 mg/kg) was selected for further behavioural testing.

### 4. DISCUSSION

With seventy-seven publications spanning greater than half-a-century, there is huge variation in study design and quality across the articles reported in this review. The validity of the individual studies has not been systematically evaluated according to current ARRIVE guidelines for health research; this review assessed trial quality by assigning several variables as proxy measures providing some reflection of study quality. These metrics include the reporting of housing conditions, sex of animals and the number of behavioural assays conducted per publication. Many studies failed to report important study variables such as housing, sex of the animals and pre-training protocols.

Laboratory research on non-human animals is conducted in stressful and artificial environments. Serotonin is particularly context sensitive; comparable elevations in brain serotonin in rats are found in both prolonged forced activity (such as forced 15-minute cold swim assays) and from mild handling of the animal alone, resulting in the clear possibility of confounding results due to environmental stressors (Collins et al., 1966).

#### 4.1 Therapeutic Dose Range of Psilocybin

It is not entirely clear how treatment doses used in animal studies compare to those investigated in healthy human volunteers or clinical trials, or if humans and the various non-human animals investigated demonstrate comparable behavioural responses. Care should be taken when comparing dose-dependent effects across non-human animal species and in considering translation of results to humans. In human clinical trials, psilocybin doses typically fall in the 20 – 30 mg/ 70 kg (0.29 – 0.43 mg/kg) dose range (Garcia-Romeu et al., 2021). Animal doses should not be converted into human equivalent doses; rather, *allometric* approaches consider the differences in body surface area when extrapolating doses of therapeutic agents (Nair & Jacob, 2016). Body surface area (BSA) normalization correlates well across several mammalian species and factors in several biological parameters including oxygen utilization, caloric expenditure, basal metabolism, blood volume and plasma proteins as well as renal function (Reagan-Shaw et al., 2008). One study in our review determined a comparable dose range between pigs and humans by assessing characteristic behavioural effects, cerebral 5-HT_2A_ receptor occupancy, and plasma psilocin levels following psilocybin injection (Donovan et al., 2020). Dose conversion estimates are different for mice, rats, and other species. What is clear is that even at doses far exceeding those administered to humans, psilocybin displays a strong biological safety profile.

#### 4.2 Sex as a Biological Variable

Sex as a biological variable is critical to research design, analysis, and reporting. To address the overrepresentation of males in biomedical research, the U.S. National Institute of Health instituted a policy in 2015 requiring investigators to consider sex as a biological variable in all vertebrate animal and human research ^1^. Female mammals have been underrepresented, often under the mistaken belief that non-human female animals display excessive variability (Beery & Zucker, 2011). One influential review conducted across ten biological fields in 2009 found male bias evident and most prominent in neuroscience, with single-sex studies outnumbering those of females by a factor of 5.5:1 (Beery & Zucker, 2011). A follow up study in 2019 searched thirty-four journals across nine biological disciplines, finding a significant increase in the proportion of studies that included both sexes but with minor change at all in the proportion of studies which reported data analyzed by sex (Woitowich et al., 2020).

Forty of the 77 studies in this review investigated exclusively males, this body of animal research is characteristic of the larger body of biased literature in preclinical neuroscience and pharmacology. One study noted their rationale for exclusion of female rats by citing the lack of previous research investigating baseline behavioural parameters in Wistar-Kyoto rats (Hibicke et al., 2020a) while another concludes that further studies are required for evaluating anti-depressant life effects of psilocybin in female mice, in particular the identification of robust stress-sensitive endpoints (Hesselgrave et al., 2021).

Greater understanding of sex and gender differences in psilocybin, psilocin and whole-mushroom effects are needed. Many studies in this review did identify differential effects between male and female study animals. Among humans, subjective effects of psychedelic compounds may differ between sexes (Liechti & Vollenweider, 2001); female sex is identified as a risk factor for adverse drug reactions (Bale & Epperson, 2017) and for challenging experiences under psychedelics (Bienemann et al., 2020).

#### 4.3 Time of Day and Circadian Rhythms

Time of day of PSI dose administration was rarely reported. Like human trials, which commonly administered drug doses early in the day, animal studies are conducted in daytime, and animal subjects are commonly housed in conditions of artificial light with a twelve-hour light and dark cycle. Animal models not covered in this review have demonstrated the significance of timing for psychedelic drug administration; in crickets LSD disrupted locomotor activity when administered early in the light phase but not later suggesting that serotonin participates in the regulation of circadian rhythm of locomotor activity (Cymborowski, 1963) while DOI induced wet dog shakes in rats were found to peak late in the light phase (Nagayama & Lu, 1996). Time of dose administration has variable effects on human behavioural outcomes including sleep (Schindler et al., 2018). In addition to differences among species, between sexes, and different routes of administration and drug formulation, the methods of measuring time points and duration must be considered in the effects of psychedelics.

#### 4.4 Research Domains Criteria Matrix Construct

We mapped the results of behavioural investigations of psilocybin in non-human animals within the units of analysis provided by the RDoC framework, providing our results with an additional level of model validation. The RDoC behavioural domains provide basic construct, network and phenomenological homologies across human and non-human experimental animals; while the Cognitive Systems requires further clarification, animal models in the domains of Negative and Positive Valence, Social Processes and Arousal Systems as well as general biological regulation are considered valid, reliable and translatable as phenotypes of human function and pathology (Anderzhanova et al., 2017).

##### 4.4.1 Negative Valence Systems

Psychedelics have been reported elsewhere to modulate fear and threat responses, decrease emotional avoidance, lessen rumination, and reduce rejection sensitivity (Kelly et al., 2021). In the area of fear conditioning, this review found low doses to decrease response latency, indicating heightened sensitivity, while larger doses demonstrate time-dependent effects. While three studies show no effect of Psi on FST behavior, and three studies report decreased time spent immobile following treatment (indicating less despair), peak behavioural effects are noted at 35 days post Psi demonstrating time-dependent persisting therapeutic effects. The FST is considered a better measure of stress coping strategies than depression-type symptoms (Commons et al., 2017). Psi reversed the effects of chronic stress in adolescent rodents while demonstrating acute dose-dependent inhibition of defensive aggressive behaviours. Animals demonstrated increased time in open arms in post-acute and in persisting effect time periods in the elevated plus maze but decreased exploratory time, along with decreased locomotion, in the open field test during acute drug effects. Time, dose and sex-dependent variables all affect the behaviour of experimental animals in negative valence constructs.

Psilocybin attenuates acute and sustained fear or threat responses. Human studies demonstrate the manner in which psilocybin, in a dose and time-dependent manner reduces hyper-reactivity of the amygdala, associated with negative affect and attentional bias to negatively valanced stimuli in the time period after Psi administration for up to one-month (Barrett et al., 2020) though increases have been noted in the day immediately following (Roseman et al., 2018). Low doses of Psi have been found to enhance the extinction of fear conditioned responses more rapidly than high doses or in control groups, indicating that Psi produces alterations in hippocampal neurogenesis (Catlow et al., 2013).

Psi has been found to acutely increase plasma corticosterone and anxiety-type behaviours in the OFT with acute anxiogenic effects correlated with post-acute anxiolytic effects; in this way Psi administration may be protective against future stressful events (Shao et al., 2021). Hibicke and colleagues conjecture that Psi administration may open a period of subsequent behavioural flexibility, noting that a single treatment with Psi produced context-dependent, long-lasting antidepressant-like and anxiolytic effects in male WKY rats in both FST and EPM assays (Hibicke et al., 2019, 2020b; C. Nichols & Hibicke, 2020). In providing evidence that Psi ameliorates stress-related behavioral deficits in mice, Shao et al., further establish that Psi increases spine density and spine size in frontal cortical pyramidal cells, evoking structural remodeling which persistent for at least 1 month and that this documented dendritic rewiring is accompanied by elevated excitatory neurotransmission (Shao et al., 2021). In pigs, a single dose of Psi resulted in increased levels of hippocampal synaptic vesicle protein 2A and lowered hippocampal and prefrontal cortex 5-HT_2A_ receptors density (Raval et al., 2020).

##### 4.4.2 Positive Valence Systems

Psychedelic compounds have been reported to increase responsiveness to reward, approach motivation and reward learning, suggesting that psychedelic therapies may help to recalibrate negative hyper-responsivity and increase positive valence across a range of neuropsychiatric disorders by altering maladaptive signaling in mesolimbic reward circuitry (Kelly et al., 2021). Studies included in this review demonstrate that Psi reduces binge-eating behaviour at several doses. Psi inhibits compulsive marble-burying behaviour in a dose-dependent manner, with most effects evident at the medium-high but not highest dose range. Psi exerts rapid antianhedonic actions in chronically stressed mice. Psilocybin demonstrates a biphasic dose and time response, with decreases in behavioural response evidenced early followed by subsequent increases in hedonic behaviours. At lower doses Psi demonstrates discriminative properties with fewer grossly disruptive effects on behaviour. While one investigation into alcohol relapse behavior among rodents included in this study found only sub-chronic Psi treatment to have a short-lasting anti-relapse effect (Meinhardt et al., 2020), a separate study found psilocybin could restore mGluR2 expression and reduce alcoholic relapse behavior (Meinhardt et al., 2021). A rodent food-reward model found low dose Psi enhanced motivation and attention while reducing impulsivity in low performing rats (Higgins et al., 2021).

##### 4.4.3 Cognitive Systems

The Cognitive Systems domain includes subconstructs such as attention, perception, working memory, declarative memory, language behavior and cognitive control. Dysfunctions in cognitive control are common across multiple psychopathologies, including depression, addiction, and obsessive-compulsive disorder. Investigations of cognitive flexibility are captured across multiple domains, including assays of behavioural flexibility. The animal paradigms reported here consist solely of tests of the visual system Psi treatment dose-dependently disrupts size discrimination and visual accuracy. Psi’s persisting effects rescue chronic stress-induced visual system impairments.

The results found in animal paradigms mirror those from human trials. 5-HT-2A receptors are richly expressed in the visual cortex. Classical psychedelics such as psilocybin have significant acute effects on the visual system, including contraction of nearby visual space (Fischer et al., 1970) and a reduced rate of binocular rivalry, a visual process thought to be linked to levels of arousal and attention (Carter et al., 2007). Psi reduces coherent motion sensitivity but does not significantly affect local motion contrast sensitivity, meaning study subjects were still capable of attending to stimuli and accurately reporting perceptions while under psilocybin (Carter et al., 2005). The closed-eye visual perceptions which occur under psychedelics occur as the early visual system of the brain behaves as if it were perceiving spatially localized visual inputs (Roseman et al., 2016), a dynamic consistent with reduced top-down inhibition of visual constraints and relaxed thalamic gating.

The impact of psilocybin on visual processes may be key to its therapeutic effects. Visual processes are considered ancient and pre-verbal cognitive processes; the top-down inhibition affected by psychedelics may induce psychological changes while overwriting stress-related neuroepigenetic coding of past fears, and as such, the visual effects of psychedelics may be associated with their demonstrated effects in resetting fear-responses (Császár-Nagy, 2019). The visual cortex is itself highly associated with working memory. Memory is an essential cognitive process necessary for learning and therefore critical to the adaptability of an organism.

Psilocybin produces dose-dependent impairments in memory task performance while increasing the vividness of autobiographical memories (Healy, 2021). Psi impairs high-level but not low-level motor perception (Carter et al., 2005). Consistently, the experience of listening to music while under psychedelics increases flow of information from the parahippocampus to the visual cortex, a phenomenon consistent with increases in bottom-up signaling and reduced top-down cognitive control under both Psi and LSD (R. L. Carhart-Harris & Friston, 2019; Kaelen, 2017). Impairments in visual tasks are associated with deficits in working memory. Mechanistically, basal ganglia function is also implicit in working memory, acting to enhance focus on a target while suppressing irrelevant distractors during working memory tasks, a function critical to the initial encoding of memory (Tyng et al., 2017). Working memory lays the foundation for later, and other cognitive controls.

The constructs of cognitive and behavioural flexibility are intertwined (Uddin, 2021). While behavioural flexibility refers to adaptive changes in behaviour in response to changes in the environment as demonstrated in animal assays mapped across other RDoC domains, cognitive flexibility is understood as the ability to switch between thinking about different concepts in a shared context. Cognitive task switching and salience processing are associated with functioning of the claustrum, situated between the putamen and the insular cortex, and known to hold dense supplies of serotonergic receptors and glutamatergic connectivity to areas of the cerebral cortex. Psi acutely reduces claustrum activity and alters connectivity to the default mode network (DMN) and frontoparietal task control network (FPTC) (Barrett et al., 2020). The claustrum has been identified as a hub in a “two-hit” model of psychedelic action modulating sub-cortical activity (Doss et al., 2021); disruptions in the prefrontal cortex account for acute effects, and the claustrum mediates connectivity to the thalamus, striatum and parahippocampal gyrus to produce downstream effects. In this way, psilocybin may assist in pausing, or disrupting established negatively-valenced habits and in potentiating revived emotion and improved learning within.

##### 4.4.4 Systems for Social Processes

Our review found that psilocybin produces acute decreases (reduced social grooming, increased distancing) in affiliative social behaviour in non-human primates. Biphasic, dose-dependent effect on sexual receptivity among rats were also found, with lower doses significantly increasing sexual receptivity and higher doses having a non-significant effect. Psilocybin treatment is consistent in reducing aggressive behaviour, with greater inhibition of aggressive behaviour as a function of increasing dose, and with greater treatment effects the longer the pre-testing isolation period of study animals.

In human trials, psilocybin induced changes in social processing systems; specifically, social reward processing (Kelly et al., 2021). The administration of Psi is associated with post treatment increases in openness (Erritzoe et al., 2018), connectedness (R. L. Carhart-Harris et al., 2018; Watts & Luoma, 2020) and nature relatedness (Lyons & Carhart-Harris, 2018). Psychedelics, including Psi, have been found to have the potential to modulate social processing, increasing emotional empathy and altruistic behaviours while reducing negative emotional recognition and sensitivity to social rejection (Preller & Vollenweider, 2019). Increases in interpersonal closeness have been sustained for up to fourteen months after Psi administration (Griffiths et al., 2011). A recent animal study in pre-publication found psilocybin reduces social deficits characteristic of autism (Mollinedo-Gajate, 2020).

Social cognition falls under the Social Processes construct of the RDoC framework but is also related to visual, cognitive, threat processing. Psilocybin decreases threat sensitivity in the visual cortex during emotional processing, suggesting the therapeutic potential of psilocybin may again be found in its ability to shift emotional biases away from negative and towards positive valence, enhancing the processing of positive stimuli (Kraehenmann et al., 2016) and again suggesting the implication of limbic system components such as the amygdala as an associated with the underlying biological mechanisms of action. In both the Negative Valence and Social Processes domains, Psi has been shown to reduce the effects of fear conditioning, indicating therapeutic potential in the treatment response to psychopathologies related to fear, chronic threat (anxiety) and chronic stress. Psi has been trialed in health conditions of anxiety rooted in fear or loss (advanced cancer diagnosis and demoralization) but given its acute effects in producing anxiety-type physiological responses and the potential for challenging experiences, psilocybin should be considered carefully before its use in therapy for Generalized Anxiety Disorder (GAD).

##### 4.4.5 Arousal Systems

Our review found acute changes in general arousal because of psilocybin administration, accompanied by mydriasis, piloerection, elevated heart rate and respiratory rate, and decreased appetite. Psilocybin is acutely wake-promoting, at the expense of REM sleep, with no persisting changes in sleep-wake behaviour. Among some animals and in some tests at lower doses Psi increased total activity while reducing in other tests. Evidence of reduced locomotor activities at higher doses is clear, as is some stimulation of activity at lower doses. Psilocybin demonstrates dose-dependent effects on startle response. Low doses increase startle amplitude or have no effect while higher doses acutely depress startle response.

Psychedelics have been found to acutely modulate the Autonomic Nervous System, the neuroendocrine and immune systems and do so in part by modulating activities of the corticolimbic networks and reducing inflammation (Flanagan & Nichols, 2018; Kelly et al., 2021; Kuypers, 2019; Schindler et al., 2018). Psilocybin acutely activates components of the sympathetic nervous system across animal species, including humans, producing increases in heart rate, blood pressure, pupillary dilation, and body temperature. Psilocybin also acutely stimulates the endocrine system, resulting in acute increases in cortisol and corticotropin (Woody, 2015). Psychedelics are also believed to have immunomodulatory and anti-inflammatory effects (Thompson & Szabo, 2020) due to via 5-HT_1_, 5-HT_2_, and sigma-1 receptor activity (Flanagan & Nichols, 2018).

Given the established sensitivity to context found with psilocybin and other classical psychedelics, arousal is key to the understanding of therapeutic effect. Arousal is considered a continuum of sensitivity of the organism to both internal and external stimuli, facilitates interaction with the environment in a context-specific manner, can be modulated by the significance of the stimuli, is distinct from motivation, and is also regulated by basic biological homeostatic drives such as hunger, sleep, thirst, and sex. Disruptions of the cortico-limbic circuit are associated with levels of heart rate variability, reflecting the physiological capacity for flexible emotional regulation in response to stress (Woody, 2015) and suggesting heart rate variability (HRV) as a relevant biomarker in assessing any potential therapeutic improvements after psilocybin administration as would cerebral spinal fluid markers of neuroinflammation. The safety and efficacy demonstrated in the psilocybin migraine trial is of note, not simply due to that trial being the sole Psi trial for a health condition not considered a psychiatric condition, and because it highlights the impact of Psi on neuroendocrine and immunomodulatory processes which may underlie its therapeutic effects.

##### 4.4.6 Sensorimotor Systems

Our review of animal studies found that stereotyped behaviours such as head-twitch response are reliably induced in non-human animals following Psi treatment, once the dose crosses a threshold of efficacy, and are potentially suppressed at higher doses. Extremely high doses also induce myoclonus or head rotation in a dose-dependent manner. Other stereotyped motor behaviours have been observed following Psi administration in rodents, cats, and non-human primates, including myoclonic spasm and limb flicking. Biphasic effects, with no reductions in grooming at lower doses but disruptions evident at standard doses and above. Effects on grooming may be related to the time-course of the drug effect. Sex-dependent differences are reported in mice. Dose-dependent effect on discoordination with no effect at lower doses and discoordination at very doses are also reported. Psi effectively disrupts PPI in rats. This effect may be dose-dependent and have sex-specific effects. 1 mg/kg psilocin (IP) effectively disrupts PPI ASR in rats, but lower doses (0.25 mg/kg) have no effect and higher doses (4 mg/kg) have variable effects.

Sensorimotor systems are responsible for the control and execution of motor behaviours and include processes involved in the parametrization of action plans and programs based on the integration of internal and external information and on the modeling of body dynamics; they are continually refined through new sensory information and the reinforcement of information via reward learning (Kelly et al., 2021). Given the way psilocybin increases sensory awareness while reducing top-down cognitive processes associated with rumination, further investigation into sensorimotor systems under psychedelics would be beneficial. Pertinent to our hypothesis is that psilocybin recalibrates global brain networks to value cortical and subcortical networks including sensory, sensorimotor, and limbic systems, sensorimotor experience lays the groundwork for visual thought. Sensorimotor structures are derived from repeated patterns of organism-environment interaction, becoming encoded in neural processes as representations of external objects to the degree that even imagined motor actions engage the same networks as actual physical movements, a phenomenon comparable to the closed-eye visual processes evidenced under psychedelics and perhaps associated with the workings of mirror neurons in the brain (Winkelman, 2017). If symbolic values are rooted in primarily sensorimotor processes, then a globally reintegrated brain with greater valuation of attentional processes will look to continued sensorimotor activity in the update, revision and improvement of predictive mental constructs and behavioural routines.

#### 4.5 Study limitations

There is a clear possibility we have missed relevant studies and given the rapid pace of new publications in psychedelic sciences including recent animal investigations, there are certainly studies we would have missed since our most recent database search of the literature. We excluded studies which were not available in English, and which were not available to our reviewers within a reasonable timeframe. There is also a clear balance we struck between the breadth of the literature and the depth of our analysis. Several conference abstracts were included and may report duplicate results.

While we did report on housing conditions, sex as a biological variable and the number of assays per publication as proxy measurements of trial quality, there was no critical analysis of the quality of the literature included in this review and we did not assess if individual trials met modern ARRIVE Guidelines for animal research. While this resulted in more comprehensive coverage of the literature pertaining to the clinical application of psilocybin, it does limit the translation of findings into suggested policies and practice. The a priori protocol guiding this scoping review was posted in a timely manner on Queen’s University open-access repository (QSpace) but was not registered or published elsewhere.

Multiple reviewers extracted data from study publications, raising the possibility of inconsistency in addition to human error. This scoping review took years to complete as part of a program of doctoral research, raising the possibility of inconsistency in analysis over time. Given the large volume of articles identified in this review, providing an accurate and comprehensive synthesis of the literature has been challenging. Lack of a clear methodological apparatus to analyze data extraction also meant that author bias is present in the identification of core themes and conclusions. Our sense of shortcoming in this area in a previous scoping review, Mapping Psilocybin-assisted Therapies: A Scoping Review (Shore, 2019), prompted the use of the Research Domain Criteria framework as an epistemic map to organize results in this review. Further, translation from animal studies to human trials or to the understanding of human health is plagued with challenges. We have done our best to identify common biological systems across species to aid in this act of knowledge translation.

Behavioural studies are characterized by their own limitations, and behavioural studies of non-human animals are particularly limited by lack of ability to self-report subjective experiences. As subjective experiences are considered critical to psychedelic therapies and given the clear effects of psilocybin on unique aspects of human consciousness, behavioural studies in animal paradigms can help in our understanding of psilocybin effects but must be taken in context alongside human trial and naturalistic or population-health level data. The RDoC framework itself emphasizes the importance of integrating multiple levels of analysis, including genes, molecules, cells, circuits, physiology, behavior, self-report, and investigational paradigms.

### 5. CONCLUSIONS

The scoped literature on behavioural investigations of psilocybin in non-human animals is marked by heterogeneity in quality and study design with variations in findings over a period of close to sixty years. Overall, psilocybin presents a unique and strong safety profile with no evidence of biological toxicity even at massive doses which far exceed those studied in humans. Psilocybin is characterized by unique time and dose-dependent effects which can be divided into distinct units of analysis: acute effects, sub-acute and persisting effects. Mapped against the RDoC domain constructs, what emerges from our scoping review is a pattern of drug action which is significantly context and training sensitive. Problems with study quality, such as widespread failure to investigate sex differences or report housing conditions, limit the generalizability of results but the body of trial publications trend towards improved quality of research design and of reporting over time.

Psilocybin acutely disrupts cognition, arousal, and sensorimotor dynamics in a dose-dependent manner in investigatory animals. Psilocybin’s acute physiological effects resolve, but studies which measure outcomes as later time-points provide a second layer of data pertaining to sub-acute and persisting effects. Demonstrated effects of Psi include heightened acute arousal, dose-dependent sedation, reductions in fear conditioning at low doses, reduced aggression, improved valence, acute disruption of working memory, the rescuing of deficits resulting from chronic stress, and improved learning when Psi is combined with subsequent repeated environmental exposure after the resolution of drug effect. The “psychedelic pause” was first identified through an animal paradigm and captures the disruption of repetitive, conditioned associated behavioural patterns stored in memory. Psilocybin is best understood to have biphasic effects, with inhibition of past conditioning evident under acute effects, and stimulation of new behaviours in the follow-up time periods persisting for several weeks after psilocybin.

Together, these data provide credence to a proposed “two-hit” model of psychedelic action, with acute effects primarily mediated by 5-HT agonism, and downstream effects persisting after the resolution of acute drug effect and 5-HT receptor agonism. Psilocybin is best understood as dose-dependent, time-dependent, and variable in its effects depending on pre-dose training and post-effect environmental exposures.

Behavioural research in non-human animals allows for the investigation of psilocybin in dimensions and domains which far exceed those of strict DSM-V nosology. Going beyond such diagnostics as depression and substance use disorder allows for a much more nuanced understanding of psilocybin’s biologic effects across multiple dimensions as captured in the RDoC framework. The results of this study of behavioural investigations of psilocybin in non-human animals provided evidence for a temporal model of psilocybin, highlighted psilocybin’s effects on past conditioning and the importance of additional conditioning during the time periods of persisting effect. These non-human animal studies revealed behavioural phenomenology of drug effect not captured in human clinical trials, providing a biological and behavioural basis for understanding psilocybin’s potential therapeutic effect beyond the narrow and static conceptions of DSM-V based diagnostic categories.

