## Appendices for "Behavioural Investigations of Psilocybin in Animals 1962-2021: A Scoping Review"

**Appendix:**
Supporting Documents -- Behavioural Investigations of Psilocybin in Animals 1962-2021: A Scoping Review

#### Appendix A: Protocol: Scoping Review: Mapping Psilocybin Animal Research

October 25, 2019

**Team Members:**

Ron Shore, PhD Student, School of Kinesiology & Health Studies, Queen's University, Kingston, ON
Nigel Barnim, Undergraduate Psychology Student, Queen's University
Katrina Dobson, MSc Student, Research School of Behavioural and Cognitive Neurosciences, University of Groningen
Sandra McKeown, MLS, Health Sciences Librarian, Queen's University
Dr. Eric Dumont (supervisor), Department of Biomedical and Molecular Sciences, Queen's University
Dr. Craig Goldie (supervisor), Department of Medicine, Queen's University

**Objective:**

To determine the effects of psilocybin in animal studies across behavioural task clusters and neurological measures and to chart:

1. What studies have been done.
2. What behavioural tests have been used.
3. What neurological measures have been implemented.
4. What dosing modalities have been used.

Further, we wish to assess the quality of research to date in line with the ARRIVE animal research quality guidelines (Kilkenny, Browne, Cuthill, Emerson, & Altman, 2010) and to indicate areas for future research.

Through librarian-conducted literature searches and defined inclusion criteria, the relevant number of internationally published, peer-reviewed academic articles in addition to grey area literature will be identified, and the findings charted to identify the current state of psilocybin animal study research.

**Stages of Scoping Review**

A scoping review methodology was selected given the heterogeneous and emergent nature of the application of psilocybin in animal research. Following the methodological framework proposed by Arksey & O'Malley (2005) and further discussed by Levac, Colquhoun, & O'Brien (2010), this scoping review has 6 specific stages.

1. Development and formulation of research question
2. Identifcation of relevant studies
3. Study selection
4. Charting the data
5. Collating, summarizing and reporting results
6. Consultation with relevant stakeholders

**Research Question:**

What are the neurological and behavioural effects of psilocybin administered in non-human animal studies?

**Preparation / preliminary work:**

Reviewers (NB , KD & RS) will complete multiple preliminary reviews, literature research, scan abstracts for keywords, pilot inclusion criteria, pilot data extraction tool, and research background and history of psilocybin in animal models prior to conducting the actual scoping review. Reviewers will also familiarize themselves with scoping review methodology, existing reviews, and ARRIVE guidelines (see citations).

**Literature Searches / Methodology:**

A librarian from Queen's Bracken Health Sciences Library (SM) will conduct a preliminary search using MEDLINE, Embase and PsycINFO. A final search for the purpose of this scoping review will include electronic databases, grey literature sources and the reference lists of identified key studies to identify possible for inclusion. Bibliographic data, study and species characteristics as well as other indicators will be collected and analyzed using a data extraction tool developed by the research team.

**Key extraction fields / data extraction tool:**

**Table 2.** Psilocybin Scoping Review Data Extraction Tool Summary Table

| Study Characteristics | Intervention Characteristics | Research Orientation | Implications |
|---|---|---|---|
| citation | drug(s) administered | Domain of research (behavioural vs. neurological) | research question |
| year of publication | dose | outcomes measured | conclusions |
| source/country of origin | route of administration | Statistical significance | limitations |
| journal type | schedule of administration | Effect size | future research |
| sample size | quality (ARRIVE guidelines) | follow-up | |
| study design | tests used | methodology | |
| animals used | | | |

**Review Process:**

**Screening/Identifying Relevant Studies:**

Librarian (SM) to complete literature search with agreed upon keywords, filtering out for duplication.

**Study Selection:**

Two reviewers (NB & KD) to review each article abstract using Covidence (Cochrane) software. All disagreements in study reviewing will be resolved through finding consensus through discussion among the two reviewers. Articles must meet inclusion criteria to be selected.

*Inclusion Criteria:* animal studies, psilocybin, mammals

*Exclusion Criteria*: duplication, comparative study with no baseline control, competitive study, review article, not specific to neurological and/or behavioural effects

### included =

### excluded =

### duplicates=

### studies selected =

**Charting the Data:**

Reviewers to read entirety of selected studies and extract data using original data extraction tool designed in Microsoft Excel. Reviewers to cross-check for homogenous process quality control. Data entered from selected studies in agreed upon categories.

**Collating/Summarizing Data:**

Data to be summarized using standardized methodology. Key findings to be highlighted, and both practice and patient variables to be summarized. Indications for future research to also be identified.

**Consultation with Relevant Stakeholders:**

Draft results to be shared and reviewed by relevant stakeholders, including all team members and associated biostatistician, and recognized experts in the field of psilocybin and animal research, as well as individuals with scoping review research experience.

**Results Mapping / Knowledge Distribution and Reporting:**

Data from the scoping review will be mapped in at least the following manners:

1. clusters of behavioural tests and neurological measures
2. dosing modalities used
3. quality assessment of research conducted
4. chart time distribution of publication dates
5. compare and contrast disciplines of research and program aims/objectives

**Additional Findings:**

1. Gaps in literature to be identified
2. Indications for areas of future research

3. Interpretation of Key Findings

4. Indications for future animal trials

**Authorship:**

Scoping review authorship to include all team members. Publication to be sought within relevant academic journals. Opportunities to present as poster/paper or as conference presentation to be sought. Protocol to be published on Prospero.

**Appendix B: Search Strategy: Behavioural Investigations of Psilocybin in Animals**

*Initial searches executed Oct 28 2019*

**Search methods**

A comprehensive search approach was employed to locate published studies and conference materials. A preliminary search was conducted in Ovid Embase using a combination of keywords and subject headings, followed by an analysis of relevant citations to identify other relevant keywords and subject headings. The optimized Ovid Embase search strategy was then adapted for Ovid MEDLINE, Ovid EBM Reviews – Cochrane Central Register of Controlled Trials, Ovid PsycINFO, Web of Science Core Collection, and BIOSIS Previews. All databases were searched from inception up to February 2021. No publication date limits were applied. The complete search strategies for all databases are presented in Appendices. The reference lists of all eligible studies were screened to identify any additional studies.

| | |
|---|---|
| Embase | 510 |
| Ovid MEDLINE | 254 |
| Ovid EBM Reviews - Cochrane Central Register of Controlled Trials | 9 |
| PsycINFO | 117 |
| Web of Science | 162 |
| BIOSIS Previews | 693 |

Database(s): **Embase Classic+Embase** 1947 to 2019 October 25

Search Strategy:

| # | Searches | Results |
|---|---|---|
| 1 | psilocybine/ | 1676 |
| 2 | psilocybin*.mp. | 1792 |
| 3 | indocybin.mp. | 4 |
| 4 | psilocibin*.mp. | 9 |
| 5 | psilotsibin.mp. | 0 |
| 6 | teonanacatl.mp. | 7 |
| 7 | psilocin.mp. | 468 |
| 8 | baeocystin.mp. | 30 |
| 9 | 1 or 2 or 3 or 4 or 5 or 6 or 7 or 8 | 1980 |
| 10 | exp animal experiment/ | 2475333 |
| 11 | exp experimental animal/ | 679538 |
| 12 | exp animal model/ | 1324678 |
| 13 | animal/ | 1929167 |
| 14 | exp "sponge (Porifera)"/ | 7079 |
| 15 | exp placozoa/ | 196 |
| 16 | exp mesozoa/ | 21 |
| 17 | coelenterate/ | 3738 |
| 18 | Bilateria/ | 211 |
| 19 | Coelomata/ | 16 |

| | | |
|---|---|---|
| 20 | Pseudocoelomata/ | 3 |
| 21 | exp protostomia/ | 670345 |
| 22 | exp ambulacraria/ | 16135 |
| 23 | chordata/ | 1435 |
| 24 | exp cephalochordata/ | 542 |
| 25 | exp hyperotreti/ | 533 |
| 26 | exp urochordata/ | 4486 |
| 27 | vertebrate/ | 26642 |
| 28 | exp fish/ | 224583 |
| 29 | tetrapod/ | 457 |
| 30 | exp amphibia/ | 73349 |
| 31 | amniote/ | 317 |
| 32 | exp sauropsid/ | 309380 |
| 33 | exp reptile/ | 50414 |
| 34 | mammal/ | 93265 |
| 35 | "calf (mammal)"/ | 2285 |
| 36 | exp monotreme/ | 744 |
| 37 | therian/ | 55 |
| 38 | exp marsupial/ | 10740 |
| 39 | placental mammal/ | 189 |

| | | |
|---|---|---|
| 40 | exp afrotheria/ | 3847 |
| 41 | boreoeutheria/ | 18 |
| 42 | exp laurasiatheria/ | 912181 |
| 43 | exp xenarthra/ | 1739 |
| 44 | euarchontoglires/ | 36 |
| 45 | dermoptera/ | 40 |
| 46 | exp glires/ | 3948872 |
| 47 | exp scandentia/ | 1017 |
| 48 | primate/ | 30089 |
| 49 | exp prosimian/ | 3227 |
| 50 | haplorhini/ | 31892 |
| 51 | exp tarsiiform/ | 235 |
| 52 | simian/ | 285 |
| 53 | exp platyrrhini/ | 9343 |
| 54 | catarrhini/ | 118 |
| 55 | exp cercopithecidae/ | 72326 |
| 56 | ape/ | 5313 |
| 57 | exp hylobatidae/ | 711 |
| 58 | hominid/ | 5721 |
| 59 | exp orangutan/ | 541 |

| | | |
|---|---|---|
| 60 | homo neanderthalensis/ | 1038 |
| 61 | exp gorilla/ | 2983 |
| 62 | exp chimpanzee/ | 12004 |
| 63 | (animal or animals or pisces or fish or fishes or catfish or catfishes or sheatfish or silurus or arius or heteropneustes or clarias or gariepinus or fathead minnow or fathead minnows or pimephales or promelas or cichlidae or trout or trouts or char or chars or salvelinus or salmo or oncorhynchus or guppy or guppies or millionfish or poecilia or goldfish or goldfishes or carassius or auratus or mullet or mullets or mugil or curema or shark or sharks or cod or cods or gadus or morhua or carp or carps or cyprinus or carpio or killifish or eel or eels or anguilla or zander or sander or lucioperca or stizostedion or turbot or turbots or psetta or flatfish or flatfishes or plaice or pleuronectes or platessa or tilapia or tilapias or oreochromis or sarotherodon or common sole or dover sole or solea or zebrafish or zebrafishes or danio or rerio or seabass or dicentrarchus or labrax or morone or lamprey or lampreys or petromyzon or pumpkinseed or pumpkinseeds or lepomis or gibbosus or herring or clupea or harengus or amphibia or amphibian or amphibians or anura or salientia or frog or frogs or rana or toad or toads or bufo or xenopus or laevis or bombina or epidalea or calamita or salamander or salamanders or newt or newts or triturus or reptilia or reptile or reptiles or bearded dragon or pogona or vitticeps or iguana or iguanas or lizard or lizards or anguis fragilis or turtle or | 8088061 |

| | |
|---|---|
| | turtles or snakes or snake or aves or bird or birds or quail or quails or coturnix or bobwhite or colinus or virginianus or poultry or poultries or fowl or fowls or chicken or chickens or gallus or zebra finch or taeniopygia or guttata or canary or canaries or serinus or canaria or parakeet or parakeets or grasskeet or parrot or parrots or psittacine or psittacines or shelduck or tadorna or goose or geese or branta or leucopsis or woodlark or lullula or flycatcher or ficedula or hypoleuca or dove or doves or geopelia or cuneata or duck or ducks or greylag or graylag or anser or harrier or circus pygargus or red knot or great knot or calidris or canutus or godwit or limosa or lapponica or meleagris or gallopavo or jackdaw or corvus or monedula or ruff or philomachus or pugnax or lapwing or peewit or plover or vanellus or swan or cygnus or columbianus or bewickii or gull or chroicocephalus or ridibundus or albifrons or great tit or parus or aythya or fuligula or streptopelia or risoria or spoonbill or platalea or leucorodia or blackbird or turdus or merula or blue tit or cyanistes or pigeon or pigeons or columba or pintail or anas or starling or sturnus or owl or athene noctua or pochard or ferina or cockatiel or nymphicus or hollandicus or skylark or alauda or tern or sterna or teal or crecca or oystercatcher or haematopus or ostralegus or shrew or shrews or sorex or araneus or crocidura or russula or european mole or talpa or chiroptera or bat or bats or eptesicus or serotinus or myotis or dasycneme or daubentonii or pipistrelle or pipistrellus or cat or cats or felis or catus or feline or dog or dogs or canis or canine or canines or otter or otters or lutra or badger or badgers or meles or fitchew or fitch or |

| | |
|---|---|
| | foumart or foulmart or ferrets or ferret or polecat or polecats or mustela or putorius or weasel or weasels or fox or foxes or vulpes or common seal or phoca or vitulina or grey seal or halichoerus or horse or horses or equus or equine or equidae or donkey or donkeys or mule or mules or pig or pigs or swine or swines or hog or hogs or boar or boars or porcine or piglet or piglets or sus or scrofa or llama or llamas or lama or glama or deer or deers or cervus or elaphus or cow or cows or bos taurus or bos indicus or bovine or bull or bulls or cattle or bison or bisons or sheep or sheeps or ovis aries or ovine or lamb or lambs or mouflon or mouflons or goat or goats or capra or caprine or chamois or rupicapra or leporidae or lagomorpha or lagomorph or rabbit or rabbits or oryctolagus or cuniculus or laprine or hares or lepus or rodentia or rodent or rodents or murinae or mouse or mice or mus or musculus or murine or woodmouse or apodemus or rat or rats or rattus or norvegicus or guinea pig or guinea pigs or cavia or porcellus or hamster or hamsters or mesocricetus or cricetulus or cricetus or gerbil or gerbils or jird or jirds or meriones or unguiculatus or jerboa or jerboas or jaculus or chinchilla or chinchillas or beaver or beavers or castor fiber or castor canadensis or sciuridae or squirrel or squirrels or sciurus or chipmunk or chipmunks or marmot or marmots or marmota or suslik or susliks or spermophilus or cynomys or cottonrat or cottonrats or sigmodon or vole or voles or microtus or myodes or glareolus or primate or primates or prosimian or prosimians or lemur or lemurs or lemuridae or loris or bush baby or bush babies or |

| | | |
|---|---|---|
| | bushbaby or bushbabies or galago or galagos or anthropoidea or anthropoids or simian or simians or monkey or monkeys or marmoset or marmosets or callithrix or cebuella or tamarin or tamarins or saguinus or leontopithecus or squirrel monkey or squirrel monkeys or saimiri or night monkey or night monkeys or owl monkey or owl monkeys or douroucoulis or aotus or spider monkey or spider monkeys or ateles or baboon or baboons or papio or rhesus monkey or macaque or macaca or mulatta or cynomolgus or fascicularis or green monkey or green monkeys or chlorocebus or vervet or vervets or pygerythrus or hominoidea or ape or apes or hylobatidae or gibbon or gibbons or siamang or siamangs or nomascus or symphalangus or hominidae or orangutan or orangutans or pongo or chimpanzee or chimpanzees or pan troglodytes or bonobo or bonobos or pan paniscus or gorilla or gorillas or troglodytes).mp. [mp=title, abstract, heading word, drug trade name, original title, device manufacturer, drug manufacturer, device trade name, keyword, floating subheading word, candidate term word] | |
| 64 | nonhuman/ | 5999019 |
| 65 | human versus animal comparison/ | 18107 |
| 66 | human versus nonhuman data/ | 1979 |
| 67 | nonhuman*.mp. | 6005349 |
| 68 | non human*.mp. | 19848 |
| 69 | or/10-68 | 10291268 |

| | | |
|---|---|---|
| 70 | 9 and 69 | 643 |
| 71 | limit 70 to editorial | 12 |
| 72 | limit 70 to "review" | 121 |
| 73 | 70 not (71 or 72) | 510 |

Database(s): **Ovid MEDLINE(R) and Epub Ahead of Print, In-Process & Other Non-Indexed Citations, Daily and Versions(R)** 1946 to October 24, 2019

Search Strategy: (Used for Cochrane CENTRAL as well)

| # | Searches | Results |
|---|---|---|
| 1 | Psilocybin/ | 687 |
| 2 | psilocybin*.mp. | 952 |
| 3 | indocybin.mp. | 1 |
| 4 | psilocibin*.mp. | 7 |
| 5 | psilotsibin.mp. | 0 |
| 6 | teonanacatl.mp. | 3 |
| 7 | psilocin.mp. | 183 |
| 8 | baeocystin.mp. | 20 |
| 9 | 1 or 2 or 3 or 4 or 5 or 6 or 7 or 8 | 992 |
| 10 | exp Animal Experimentation/ | 9204 |
| 11 | exp models, animal/ | 548939 |

| | | |
|---|---|---|
| 12 | exp Animal Population Groups/ | 1159640 |
| 13 | exp Chordata, Nonvertebrate/ | 5819 |
| 14 | exp Amphibians/ | 107366 |
| 15 | exp Birds/ | 218464 |
| 16 | exp Fishes/ | 184024 |
| 17 | exp Reptiles/ | 38951 |
| 18 | Hyraxes/ | 146 |
| 19 | exp Insectivora/ | 4899 |
| 20 | exp Marsupialia/ | 10112 |
| 21 | exp Monotremata/ | 668 |
| 22 | exp Proboscidea Mammal/ | 2111 |
| 23 | exp Xenarthra/ | 1602 |
| 24 | Eutheria/ | 46 |
| 25 | Mammals/ | 29233 |
| 26 | Vertebrates/ | 11387 |
| 27 | Chordata/ | 576 |
| 28 | exp Artiodactyla/ | 673890 |
| 29 | exp Carnivora/ | 461474 |
| 30 | exp Cetacea/ | 8874 |
| 31 | Chiroptera/ | 9171 |

| | | |
|---|---|---|
| 32 | exp Lagomorpha/ | 338140 |
| 33 | exp Perissodactyla/ | 70588 |
| 34 | exp Rodentia/ | 3161163 |
| 35 | exp Scandentia/ | 1142 |
| 36 | exp Sirenia/ | 319 |
| 37 | Primates/ | 11880 |
| 38 | exp Strepsirhini/ | 2515 |
| 39 | Haplorhini/ | 46512 |
| 40 | exp Platyrrhini/ | 13744 |
| 41 | exp Tarsii/ | 77 |
| 42 | Catarrhini/ | 93 |
| 43 | exp Cercopithecidae/ | 121236 |
| 44 | exp Hylobatidae/ | 945 |
| 45 | Hominidae/ | 8112 |
| 46 | Gorilla gorilla/ | 1898 |
| 47 | Neanderthals/ | 633 |
| 48 | Pan paniscus/ | 541 |
| 49 | Pan troglodytes/ | 9321 |
| 50 | exp Pongo/ | 1258 |
| 51 | Animals/ | 6500190 |

| | | |
|---|---|---|
| 52 | (animal or animals or pisces or fish or fishes or catfish or catfishes or sheatfish or silurus or arius or heteropneustes or clarias or gariepinus or fathead minnow or fathead minnows or pimephales or promelas or cichlidae or trout or trouts or char or chars or salvelinus or salmo or oncorhynchus or guppy or guppies or millionfish or poecilia or goldfish or goldfishes or carassius or auratus or mullet or mullets or mugil or curema or shark or sharks or cod or cods or gadus or morhua or carp or carps or cyprinus or carpio or killifish or eel or eels or anguilla or zander or sander or lucioperca or stizostedion or turbot or turbots or psetta or flatfish or flatfishes or plaice or pleuronectes or platessa or tilapia or tilapias or oreochromis or sarotherodon or common sole or dover sole or solea or zebrafish or zebrafishes or danio or rerio or seabass or dicentrarchus or labrax or morone or lamprey or lampreys or petromyzon or pumpkinseed or pumpkinseeds or lepomis or gibbosus or herring or clupea or harengus or amphibia or amphibian or amphibians or anura or salientia or frog or frogs or rana or toad or toads or bufo or xenopus or laevis or bombina or epidalea or calamita or salamander or salamanders or newt or newts or triturus or reptilia or reptile or reptiles or bearded dragon or pogona or vitticeps or iguana or iguanas or lizard or lizards or anguis fragilis or turtle or turtles or snakes or snake or aves or bird or birds or quail or quails or coturnix or bobwhite or colinus or virginianus or poultry or poultries or fowl or fowls or chicken or chickens or gallus or zebra finch or taeniopygia or guttata or canary or canaries or serinus or canaria or parakeet or parakeets or grasskeet or parrot or parrots | 7288178 |

| | |
|---|---|
| | or psittacine or psittacines or shelduck or tadorna or goose or geese or branta or leucopsis or woodlark or lullula or flycatcher or ficedula or hypoleuca or dove or doves or geopelia or cuneata or duck or ducks or greylag or graylag or anser or harrier or circus pygargus or red knot or great knot or calidris or canutus or godwit or limosa or lapponica or meleagris or gallopavo or jackdaw or corvus or monedula or ruff or philomachus or pugnax or lapwing or peewit or plover or vanellus or swan or cygnus or columbianus or bewickii or gull or chroicocephalus or ridibundus or albifrons or great tit or parus or aythya or fuligula or streptopelia or risoria or spoonbill or platalea or leucorodia or blackbird or turdus or merula or blue tit or cyanistes or pigeon or pigeons or columba or pintail or anas or starling or sturnus or owl or athene noctua or pochard or ferina or cockatiel or nymphicus or hollandicus or skylark or alauda or tern or sterna or teal or crecca or oystercatcher or haematopus or ostralegus or shrew or shrews or sorex or araneus or crocidura or russula or european mole or talpa or chiroptera or bat or bats or eptesicus or serotinus or myotis or dasycneme or daubentonii or pipistrelle or pipistrellus or cat or cats or felis or catus or feline or dog or dogs or canis or canine or canines or otter or otters or lutra or badger or badgers or meles or fitchew or fitch or foumart or foulmart or ferrets or ferret or polecat or polecats or mustela or putorius or weasel or weasels or fox or foxes or vulpes or common seal or phoca or vitulina or grey seal or halichoerus or horse or horses or equus or equine or equidae or donkey or donkeys or mule or mules or pig or pigs or swine or swines or hog or hogs or |

| | |
|---|---|
| | boar or boars or porcine or piglet or piglets or sus or scrofa or llama or llamas or lama or glama or deer or deers or cervus or elaphus or cow or cows or bos taurus or bos indicus or bovine or bull or bulls or cattle or bison or bisons or sheep or sheeps or ovis aries or ovine or lamb or lambs or mouflon or mouflons or goat or goats or capra or caprine or chamois or rupicapra or leporidae or lagomorpha or lagomorph or rabbit or rabbits or oryctolagus or cuniculus or laprine or hares or lepus or rodentia or rodent or rodents or murinae or mouse or mice or mus or musculus or murine or woodmouse or apodemus or rat or rats or rattus or norvegicus or guinea pig or guinea pigs or cavia or porcellus or hamster or hamsters or mesocricetus or cricetulus or cricetus or gerbil or gerbils or jird or jirds or meriones or unguiculatus or jerboa or jerboas or jaculus or chinchilla or chinchillas or beaver or beavers or castor fiber or castor canadensis or sciuridae or squirrel or squirrels or sciurus or chipmunk or chipmunks or marmot or marmots or marmota or suslik or susliks or spermophilus or cynomys or cottonrat or cottonrats or sigmodon or vole or voles or microtus or myodes or glareolus or primate or primates or prosimian or prosimians or lemur or lemurs or lemuridae or loris or bush baby or bush babies or bushbaby or bushbabies or galago or galagos or anthropoidea or anthropoids or simian or simians or monkey or monkeys or marmoset or marmosets or callithrix or cebuella or tamarin or tamarins or saguinus or leontopithecus or squirrel monkey or squirrel monkeys or saimiri or night monkey or night monkeys or owl monkey or owl monkeys or douroucoulis or |

| | | |
|---|---|---|
| | aotus or spider monkey or spider monkeys or ateles or baboon or baboons or papio or rhesus monkey or macaque or macaca or mulatta or cynomolgus or fascicularis or green monkey or green monkeys or chlorocebus or vervet or vervets or pygerythrus or hominoidea or ape or apes or hylobatidae or gibbon or gibbons or siamang or siamangs or nomascus or symphalangus or hominidae or orangutan or orangutans or pongo or chimpanzee or chimpanzees or pan troglodytes or bonobo or bonobos or pan paniscus or gorilla or gorillas or troglodytes).mp. [mp=title, abstract, original title, name of substance word, subject heading word, floating sub-heading word, keyword heading word, organism supplementary concept word, protocol supplementary concept word, rare disease supplementary concept word, unique identifier, synonyms] | |
| 55 | nonhuman*.mp. | 17013 |
| 54 | non human*.mp. | 13927 |
| 55 | or/10-54 | 7295497 |
| 56 | 9 and 55 | 254 |

Database(s): **PsycINFO** 1806 to October Week 2 2019

Search Strategy:

| # | Searches | Results |
|---|---|---|
| 1 | Psilocybin/ | 209 |
| 2 | psilocybin*.mp. | 506 |

| | | |
|---|---|---|
| 3 | indocybin.mp. | 0 |
| 4 | psilocibin*.mp. | 2 |
| 5 | psilotsibin.mp. | 0 |
| 6 | teonanacatl.mp. | 0 |
| 7 | psilocin.mp. | 35 |
| 8 | baeocystin.mp. | 2 |
| 9 | 1 or 2 or 3 or 4 or 5 or 6 or 7 or 8 | 516 |
| 10 | animal models/ | 32226 |
| 11 | exp Animals/ | 343015 |
| 12 | (animal or animals or pisces or fish or fishes or catfish or catfishes or sheatfish or silurus or arius or heteropneustes or clarias or gariepinus or fathead minnow or fathead minnows or pimephales or promelas or cichlidae or trout or trouts or char or chars or salvelinus or salmo or oncorhynchus or guppy or guppies or millionfish or poecilia or goldfish or goldfishes or carassius or auratus or mullet or mullets or mugil or curema or shark or sharks or cod or cods or gadus or morhua or carp or carps or cyprinus or carpio or killifish or eel or eels or anguilla or zander or sander or lucioperca or stizostedion or turbot or turbots or psetta or flatfish or flatfishes or plaice or pleuronectes or platessa or tilapia or tilapias or oreochromis or sarotherodon or common sole or dover sole or solea or zebrafish or zebrafishes or danio or rerio or seabass or dicentrarchus or labrax or morone or lamprey or lampreys or petromyzon or pumpkinseed or pumpkinseeds or | 502557 |

| | |
|---|---|
| | lepomis or gibbosus or herring or clupea or harengus or amphibia or amphibian or amphibians or anura or salientia or frog or frogs or rana or toad or toads or bufo or xenopus or laevis or bombina or epidalea or calamita or salamander or salamanders or newt or newts or triturus or reptilia or reptile or reptiles or bearded dragon or pogona or vitticeps or iguana or iguanas or lizard or lizards or anguis fragilis or turtle or turtles or snakes or snake or aves or bird or birds or quail or quails or coturnix or bobwhite or colinus or virginianus or poultry or poultries or fowl or fowls or chicken or chickens or gallus or zebra finch or taeniopygia or guttata or canary or canaries or serinus or canaria or parakeet or parakeets or grasskeet or parrot or parrots or psittacine or psittacines or shelduck or tadorna or goose or geese or branta or leucopsis or woodlark or lullula or flycatcher or ficedula or hypoleuca or dove or doves or geopelia or cuneata or duck or ducks or greylag or graylag or anser or harrier or circus pygargus or red knot or great knot or calidris or canutus or godwit or limosa or lapponica or meleagris or gallopavo or jackdaw or corvus or monedula or ruff or philomachus or pugnax or lapwing or peewit or plover or vanellus or swan or cygnus or columbianus or bewickii or gull or chroicocephalus or ridibundus or albifrons or great tit or parus or aythya or fuligula or streptopelia or risoria or spoonbill or platalea or leucorodia or blackbird or turdus or merula or blue tit or cyanistes or pigeon or pigeons or columba or pintail or anas or starling or sturnus or owl or athene noctua or pochard or ferina or cockatiel or nymphicus or hollandicus or skylark or alauda or tern or sterna or teal or crecca or |

| | |
|---|---|
| | oystercatcher or haematopus or ostralegus or shrew or shrews or sorex or araneus or crocidura or russula or european mole or talpa or chiroptera or bat or bats or eptesicus or serotinus or myotis or dasycneme or daubentonii or pipistrelle or pipistrellus or cat or cats or felis or catus or feline or dog or dogs or canis or canine or canines or otter or otters or lutra or badger or badgers or meles or fitchew or fitch or foumart or foulmart or ferrets or ferret or polecat or polecats or mustela or putorius or weasel or weasels or fox or foxes or vulpes or common seal or phoca or vitulina or grey seal or halichoerus or horse or horses or equus or equine or equidae or donkey or donkeys or mule or mules or pig or pigs or swine or swines or hog or hogs or boar or boars or porcine or piglet or piglets or sus or scrofa or llama or llamas or lama or glama or deer or deers or cervus or elaphus or cow or cows or bos taurus or bos indicus or bovine or bull or bulls or cattle or bison or bisons or sheep or sheeps or ovis aries or ovine or lamb or lambs or mouflon or mouflons or goat or goats or capra or caprine or chamois or rupicapra or leporidae or lagomorpha or lagomorph or rabbit or rabbits or oryctolagus or cuniculus or laprine or hares or lepus or rodentia or rodent or rodents or murinae or mouse or mice or mus or musculus or murine or woodmouse or apodemus or rat or rats or rattus or norvegicus or guinea pig or guinea pigs or cavia or porcellus or hamster or hamsters or mesocricetus or cricetulus or cricetus or gerbil or gerbils or jird or jirds or meriones or unguiculatus or jerboa or jerboas or jaculus or chinchilla or chinchillas or beaver or beavers or castor fiber or castor canadensis or sciuridae or squirrel or squirrels |

| | | |
|---|---|---|
| | or sciurus or chipmunk or chipmunks or marmot or marmots or marmota or suslik or susliks or spermophilus or cynomys or cottonrat or cottonrats or sigmodon or vole or voles or microtus or myodes or glareolus or primate or primates or prosimian or prosimians or lemur or lemurs or lemuridae or loris or bush baby or bush babies or bushbaby or bushbabies or galago or galagos or anthropoidea or anthropoids or simian or simians or monkey or monkeys or marmoset or marmosets or callithrix or cebuella or tamarin or tamarins or saguinus or leontopithecus or squirrel monkey or squirrel monkeys or saimiri or night monkey or night monkeys or owl monkey or owl monkeys or douroucoulis or aotus or spider monkey or spider monkeys or ateles or baboon or baboons or papio or rhesus monkey or macaque or macaca or mulatta or cynomolgus or fascicularis or green monkey or green monkeys or chlorocebus or vervet or vervets or pygerythrus or hominoidea or ape or apes or hylobatidae or gibbon or gibbons or siamang or siamangs or nomascus or symphalangus or hominidae or orangutan or orangutans or pongo or chimpanzee or chimpanzees or pan troglodytes or bonobo or bonobos or pan paniscus or gorilla or gorillas or troglodytes).mp. [mp=title, abstract, heading word, table of contents, key concepts, original title, tests & measures, mesh] | |
| 13 | nonhuman*.mp. | 12391 |
| 14 | non human*.mp. | 4498 |
| 15 | 10 or 11 or 12 or 13 or 14 | 508257 |

| | | |
|---|---|---|
| 16 | 9 and 15 | 117 |

Web of Science (used for BIOSIS Previews as well)

**TOPIC:** (psilocybin* or indocybin or psilocibin* or psilotsibin or teonanacatl or psilocin or baeocystin) *AND* **TOPIC:** ((mammal or mammals or nonhuman or non-human or non-humans or nonhumans or animal or animals or pisces or fish or fishes or catfish or catfishes or sheatfish or silurus or arius or heteropneustes or clarias or gariepinus or fathead minnow or fathead minnows or pimephales or promelas or cichlidae or trout or trouts or char or chars or salvelinus or salmo or oncorhynchus or guppy or guppies or millionfish or poecilia or goldfish or goldfishes or carassius or auratus or mullet or mullets or mugil or curema or shark or sharks or cod or cods or gadus or morhua or carp or carps or cyprinus or carpio or killifish or eel or eels or anguilla or zander or sander or lucioperca or stizostedion or turbot or turbots or psetta or flatfish or flatfishes or plaice or pleuronectes or platessa or tilapia or tilapias or oreochromis or sarotherodon or common sole or dover sole or solea or zebrafish or zebrafishes or danio or rerio or seabass or dicentrarchus or labrax or morone or lamprey or lampreys or petromyzon or pumpkinseed or pumpkinseeds or lepomis or gibbosus or herring or clupea or harengus or amphibia or amphibian or amphibians or anura or salientia or frog or frogs or rana or toad or toads or bufo or xenopus or laevis or bombina or epidalea or calamita or salamander or salamanders or newt or newts or triturus or reptilia or reptile or reptiles or bearded dragon or pogona or vitticeps or iguana or iguanas or lizard or lizards or anguis fragilis or turtle or turtles or snakes or snake or aves or bird or birds or quail or quails or coturnix or bobwhite or colinus or virginianus or poultry or poultries or fowl or fowls or chicken or chickens or gallus or zebra finch or taeniopygia or guttata or canary or canaries or serinus or canaria or parakeet or parakeets or grasskeet or parrot or parrots or psittacine or psittacines or shelduck or tadorna or goose or geese or branta or leucopsis or woodlark or lullula or flycatcher or ficedula or hypoleuca or dove or doves or geopelia or cuneata or duck or ducks or greylag or graylag or anser or harrier or circus pygargus or red knot or great knot or calidris or canutus or godwit or limosa or lapponica or meleagris or gallopavo or jackdaw or corvus or monedula or ruff or philomachus or pugnax or lapwing or peewit or plover or vanellus or swan or cygnus or columbianus or bewickii or gull or chroicocephalus or ridibundus or albifrons or great tit or parus or aythya or fuligula or streptopelia or risoria or spoonbill or platalea or leucorodia or blackbird or turdus or merula or blue tit or cyanistes or pigeon or pigeons or columba or pintail or anas or starling or sturnus or owl or athene noctua or pochard or ferina or cockatiel or nymphicus or hollandicus or skylark or alauda or tern or sterna or teal or crecca or oystercatcher or haematopus or ostralegus or shrew or shrews or sorex or araneus or crocidura or russula or european mole or talpa or chiroptera or bat or bats or eptesicus or serotinus or myotis or dasycneme or daubentonii or pipistrelle or pipistrellus or cat or cats or felis or catus or feline or dog or dogs or canis or canine or canines or otter or otters or lutra or badger or badgers or meles or fitchew or fitch or foumart or foulmart or ferrets or ferret or polecat or polecats or mustela or putorius or weasel or weasels or fox or foxes or vulpes or common seal or phoca or vitulina or grey seal or halichoerus or horse or horses or equus or equine or equidae or donkey or donkeys or mule or mules or pig or pigs or swine or swines or hog or hogs or boar or boars or porcine or piglet or piglets or sus or scrofa or llama or llamas or lama or glama or deer or deers or cervus or elaphus or cow or cows or bos taurus or bos indicus or bovine or bull or bulls or cattle or bison or bisons or sheep or sheeps or ovis aries or ovine or lamb or lambs or mouflon or mouflons or goat or goats or capra or caprine or chamois or rupicapra or leporidae or lagomorpha or lagomorph or rabbit or rabbits or oryctolagus or cuniculus or laprine or hares or lepus or rodentia or rodent or rodents or murinae or mouse or mice or mus or musculus or murine or woodmouse or apodemus or rat or rats or rattus or norvegicus or guinea pig or guinea pigs or cavia or porcellus or hamster or hamsters or mesocricetus or cricetulus or cricetus or gerbil or gerbils or jird or jirds or meriones or unguiculatus or jerboa or jerboas or jaculus or chinchilla or chinchillas or beaver or beavers or castor fiber or castor canadensis or sciuridae or squirrel or squirrels or sciurus or chipmunk or chipmunks or marmot or marmots or marmota or suslik or susliks or spermophilus or cynomys or cottonrat or cottonrats or sigmodon or vole or voles or microtus or myodes or glareolus or primate or primates or prosimian or prosimians or lemur or lemurs or lemuridae or loris or bush baby or bush babies or bushbaby or bushbabies or galago or galagos or anthropoidea or anthropoids or simian or simians or monkey or monkeys or marmoset or marmosets or callithrix or cebuella or tamarin or tamarins or saguinus or leontopithecus or squirrel monkey or squirrel monkeys or saimiri or night monkey or night monkeys or owl monkey or owl monkeys or douroucoulis or aotus or spider monkey or spider monkeys or ateles or baboon or baboons or papio or rhesus monkey or macaque or macaca or mulatta or cynomolgus or fascicularis or green monkey or green monkeys or chlorocebus or vervet or vervets or pygerythrus or hominoidea or ape or apes or hylobatidae or gibbon or gibbons or siamang or siamangs or nomascus or symphalangus or hominidae or orangutan or orangutans or pongo or chimpanzee or chimpanzees or pan troglodytes or bonobo or bonobos or pan paniscus or gorilla or gorillas or troglodytes))

*Indexes=SCI-EXPANDED, SSCI, A&HCI, CPCI-S, CPCI-SSH, ESCI Timespan=All years*

**Appendix C**: Behavioural Investigations of Psilocybin In Non-human Animals: List of Included and Excluded Studies

Included Studies by Date

| Title | Authors | Year | Journal |
|---|---|---|---|
| Behavioral and electrographic effects of intraventricular injection of bulbocapnine and other substances in freely moving monkeys. In: Some biological aspects of schizophrenic behavior | Wada, Juhn A. | 1962 | Ann New York Acad Sci |
| Dephosphorylation of psilocybin in the intact mouse | Horita, A.; Weber, L. J. | 1962 | Toxicol and Appl Pharmacol |
| Simultaneous studies of blood sugar, serotonin metabolism, behaviourial changes and EEG of the wake rabbit after the administration of mescaline and psilocybin | Steiner, J. E.; Sulman, F. G. | 1963 | Harokeach Haivri |
| Inhibition of isolation-induced attack behavior of mice by drugs. | Uyeno, E T | 1966 | Journal of pharmaceutical sciences |
| Psilocin: effects on behaviour and brain serotonin in mice. | Collins, R L; Ordy, J M; Samorajski, T | 1966 | Nature |
| Effects of mescaltne and psilocybin on dominance behavior of the rat | Uyeno E.T. | 1967 | Archives internationales de pharmacodynamie et de therapie |
| Studies on the effects of drugs on performance of a delayed discrimination | Roberts, M. H. T.; Bradley, P. B. | 1967 | Physiol Behav |

| | | | |
|---|---|---|---|
| The Effects of Psilocybin Cent Stim Upon the Visual Pathways of Baboons | Bert, J.; Ayats, H.; Bermond, F. | 1968 | Vagtborg, Harold (Edited by). Use of Nonhuman Primates in Drug Evaluation. A Symposium. Xii + 640p. Illus. University of Texas Press: Austin, Tex., U.S.A. And London, England |
| Behavioural effects of some morphine antagonists and hallucinogens in the rat. | Schneider, C | 1968 | Nature |
| A STUDY OF THE ROLE OF NORADRENALINE NOREPINEPHRINE IN BEHAVIORAL CHANGES PRODUCED IN THE RAT BY PSYCHOTOMIMETIC DRUGS LYSERGIC-ACID DI ETHYLAMIDE PSILOCYBIN DITRAN RESERPINE ALPHA METHYL TYROSINE METHYL ESTER HYDRO CHLORIDE METAB | Sugrue, M. F. | 1969 | British Journal of Pharmacology |
| ALTERATION OF A LEARNED RESPONSE OF THE SQUIRREL MONKEY BY HALLUCINOGENS 2 5 DI METHOXY-4-ETHYL AMPHETAMINE 2 5 DI METHOXY-4-METHYL AMPHETAMINE PSILOCYBIN 4 METHOXY-N N-DIMETHYL TRYPTAMINE N N DI METHYL TRYPTAMINE 6 HYDROXY-N N-DIMETHYL TRYPTAMINE CENT STIM | Uyeno, E. T. | 1969 | International Journal of Neuropharmacology |

| | | | |
|---|---|---|---|
| Effects of psilocybin, dimethyltryptamine, mescaline and various lysergic acid derivatives on the EEG and on photically induced epilepsy in the baboon (Papio papio). | Meldrum, B S; Naquet, R | 1971 | Electroencephalography and clinical neurophysiology |
| Relative potency of amphetamine derivatives and N,N-dimethyltryptamines. | Uyeno, E T | 1971 | Psychopharmacologia |
| PART 2 THE EFFECTS OF SOME HALLUCINOGENS ON AGGRESSIVENESS OF MICE AND RATS | Kostowski, W.; Rewerski, W.; Piechocki, T. | 1972 | Pharmacology (Basel) |
| EFFECTS OF AMPHETAMINE LSD PSILOCYBIN AND 2 5 DI METHOXY-4-METHYL AMPHETAMINE ON SCHEDULE CONTROLLED BEHAVIOR IN THE RAT | Marquis, W. J.; Tilson, H. A.; Rech, R. H. | 1973 | Federation Proceedings |
| The effects of psilocybin on EEG and behaviour in monkeys. | Horibe, M | 1974 | Activitas nervosa superior |
| COMPARISON OF DISCRIMINATIVE STIMULUS PROPERTIES OF DELTA9-THC AND PSILOCYBIN IN RATS | GREENBERG, I; KUHN, D; APPEL, JB | 1975 | PHARMACOLOGY BIOCHEMISTRY AND BEHAVIOR |
| Adaptive changes in behavior after repeated administration of various psychoactive drugs. | Rech, R H; Tilson, H A; Marquis, W J | 1975 | Advances in biochemical psychopharmacology |
| An animal behavior model for studying the actions of LSD and related hallucinogens. | Jacobs, B L; Trulson, M E; Stern, W C | 1976 | Science (New York, N.Y.) |
| DRUG INDUCED SUPPRESSION SPECIFICITY DUE TO DRUG EMPLOYED AS UNCONDITIONED STIMULI | Cameron, O. G.; Appel, J. B. | 1976 | Pharmacology Biochemistry and Behavior |

| | | | |
|---|---|---|---|
| Comparative effects of hallucinogenic drugs on rotational behavior in rats with unilateral 6-hydroxydopamine lesions. | Trulson, M E; Stark, A D; Jacobs, B L | 1977 | European journal of pharmacology |
| Serotonin and sexual behaviour in female rats. Effects of hallucinogenic indolealkylamines and phenylethylamines | Everitt B.J.; Fuxe K. | 1977 | Neuroscience Letters |
| Comparative effects of hallucinogenic drugs on behavior of the cat. | Jacobs, B L; Trulson, M E; Stark, A D; Christoph, G R | 1977 | Communications in psychopharmacology |
| Behavioral effects of LSD in the cat: proposal of an animal behavior model for studying the actions of hallucinogenic drugs. | Jacobs, B L; Trulson, M E; Stern, W C | 1977 | Brain research |
| Psilocybin: biphasic dose-response effects on the acoustic startle reflex in the rat. | Davis, M; Walters, J K | 1977 | Pharmacology, biochemistry, and behavior |
| Psychoactivity of normacromerine in animals. | Bourn, W M; Keller, W J; Bonfiglio, J F | 1978 | Life sciences |
| Evaluation of psychotropic drugs with a modified open field test | Cunha J.M.; Masur J. | 1978 | Pharmacology |
| The effects of lysergic acid diethylamide and mescaline-derived hallucinogens on sensory-integrative function: tactile startle. | Geyer, M A; Petersen, L R; Rose, G J; Horwitt, D D; Light, R K; Adams, L M; Zook, J A; Hawkins, R L; Mandell, A J | 1978 | The Journal of pharmacology and experimental therapeutics |
| The identification of LSD-like hallucinogens using the chronic spinal dog. | Martin, W R; Vaupel, D B; Nozaki, M; Bright, L D | 1978 | Drug and alcohol dependence |

| | | | |
|---|---|---|---|
| EFFECTS OF MESCALINE AND PSILOCIN ON ACQUISITION, CONSOLIDATION, AND PERFORMANCE OF LIGHT-DARK DISCRIMINATION IN 2 INBRED STRAINS OF MICE | Castellano, C. | 1978 | Psychopharmacology |
| Screening hallucinogenic drugs. II. Systematic study of two behavioral tests | Calil H.M. | 1978 | Psychopharmacology |
| PSILOCYBIN NEURO-PSYCHO-PHARMACOLOGY IN RABBITS | Guest, J.; Consroe, P. | 1979 | Research Communications in Psychology Psychiatry and Behavior |
| Severe aggression in rats induced by mescaline but not other hallucinogens. | Sbordone, R J; Wingard, J A; Gorelick, D A; Elliott, M L | 1979 | Psychopharmacology |
| THE INHIBITION OF FOOD INTAKE IN THE DOG BY LSD MESCALINE PSILOCIN D AMPHETAMINE AND PHENYL ISO PROPYLAMINE DERIVATIVES | Vaupel, D. B.; Mozaki, M.; Martin, W. R.; Bright, L. D.; Morton, E. C. | 1979 | Life Sciences |
| A characteristic effect of hallucinogens on investigatory responding in rats. | Geyer, M A; Light, R K; Rose, G J; Petersen, L R; Horwitt, D D; Adams, L M; Hawkins, R L | 1979 | Psychopharmacology |
| Comparisons of mescal bean alkaloids with mescaline, DELTA9-THC and other psychotogens | Bourn W.M.; Keller W.J.; Bonfiglio J.F. | 1979 | Life Sciences |
| GROSS BEHAVIORAL AND PHYSIOLOGICAL EFFECTS OF HALLUCINOGENS IN CONSCIOUS RESTRAINED MONKEYS MACACA-FASCICULARIS | Wilson, M. C.; Bedford, J. A.; Davis, W. M. | 1981 | Research Communications in Substances of Abuse |

| | | | |
|---|---|---|---|
| STRUCTURE ACTIVITY RELATIONSHIPS AMONG HALLUCINOGENIC TRYPTAMINE DERIVATIVES EVALUATED BY SCHEDULE CONTROLLED BEHAVIOR | Harris, R. A.; Homfeld, E.; Campaigne, E. | 1981 | Journal of Pharmacy and Pharmacology |
| Induction of 5-hydroxytryptamine (5HT)-dependent myoclonus in guinea pigs by indole-containing, but not by piperazine-containing, 5HT-agonists suggests multiple cerebral 5HT receptors | Jenner P.; Luscombe G.; Marsden C.D. | 1981 | British Journal of Pharmacology |
| Dissociations between the effects of hallucinogenic drugs on behavior and raphe unit activity in freely moving cats. | Trulson, M E; Heym, J; Jacobs, B L | 1981 | Brain research |
| PSILOCYBIN AS A DISCRIMINATIVE STIMULUS - LACK OF SPECIFICITY IN AN ANIMAL BEHAVIOR MODEL FOR HALLUCINOGENS | KOERNER, J; APPEL, JB | 1982 | PSYCHOPHARMACOLOGY |
| DOSE DEPENDENT BEHAVIORAL-CHANGES INDUCED BY PSILOCYBIN IN SELECTED MEMBERS OF A PRIMATE SOCIAL COLONY | Sink, C. A.; Beluhan, L. J.; Schlemmer, R. F.; Heinze, W. J.; Davis, J. M. | 1983 | Federation Proceedings |
| Differential effects of hallucinogenic drugs on the activity of serotonin-containing neurons in the nucleus centralis superior and nucleus raphe pallidus in freely moving cats. | Trulson, M E; Preussler, D W; Trulson, V M | 1984 | The Journal of pharmacology and experimental therapeutics |

| | | | |
|---|---|---|---|
| 5-Hydroxytryptamine (5-HT)-dependent myoclonus in guinea pigs is induced through brainstem 5-HT-1 receptors | Luscombe G.; Jenner P.; Marsden C.D. | 1984 | Neuroscience Letters |
| A primate model for the study of hallucinogens | Schlemmer Jr. R.F.; Davis J.M. | 1986 | Pharmacology Biochemistry and Behavior |
| Comparison of stimulants and hallucinogens on shuttle avoidance in rats | Davis W.M.; Hatoum H.T. | 1987 | General Pharmacology: Vascular System |
| 3,4-methylenedioxymethapmhetamine(MDMA) and 4-OH-dimethyltryptamine (psilocin) interaction in rats: Behavioral study on prepulse inhibition of acoustic startle reaction and on locomotion | Palenicek, T.; Bubenikova, V.; Votava, M. | 2005 | Behavioural Pharmacology |
| Effects of Psilocybe argentipes on marble-burying behavior in mice. | Matsushima, Yoshihiro; Shirota, Osamu; Kikura-Hanajiri, Ruri; Goda, Yukihiro; Eguchi, Fumio | 2009 | Bioscience, biotechnology, and biochemistry |
| Acute toxicity of Psilocybe cubensis (Ear.) Sing., Strophariaceae, aqueous extract in mice | Kirsten, T. B.; Bernardi, M. M. | 2010 | Revista Brasileira De Farmacognosia-Brazilian Journal of Pharmacognosy |
| Differential contributions of serotonin receptors to the behavioral effects of indoleamine hallucinogens in mice. | Halberstadt, Adam L; Koedood, Liselore; Powell, Susan B; Geyer, Mark A | 2011 | Journal of psychopharmacology (Oxford, England) |
| THE EFFECT OF TRYPTAMINE HALLUCINOGENS ON QUANTITATIVE EEG AND BEHAVIOR IN RATS | Filip, Tyls; Tomas, Palenicek; Michaela, Fujakova; Anna, Kubesova; Martin, Brunovsky; Vladimir, Krajca; Jiri, Horacek | 2011 | Behavioural Pharmacology |

| | | | |
|---|---|---|---|
| Comparison of sensorimotor gating and quantitative EEG in serotonergic models of psychosis in the rat | Tyls, F.; Palenicek, T.; Fujakova, M.; Kaderabek, L.; Novakova, P.; Kubesova, A.; Horacek, J. | 2013 | European Neuropsychopharmacology |
| Effects of psilocybin on hippocampal neurogenesis and extinction of trace fear conditioning. | Catlow, Briony J; Song, Shijie; Paredes, Daniel A; Kirstein, Cheryl L; Sanchez-Ramos, Juan | 2013 | Experimental brain research |
| The impact of serotonin system modulation on behavior and quantitative EEG in an animal model of psychosis induced by psilocin | Tyls, F.; Palenicek, T.; Novakova, P.; Kaderabek, L.; Fujakova, M.; Kubesova, A.; Horacek, J. | 2014 | European Neuropsychopharmacology |
| The effect of psilocin on memory acquisition, retrieval, and consolidation in the rat. | Rambousek, Lukas; Palenicek, Tomas; Vales, Karel; Stuchlik, Ales | 2014 | Frontiers in behavioral neuroscience |
| Research on acute toxicity and the behavioral effects of methanolic extract from psilocybin mushrooms and psilocin in mice. | Zhuk, Olga; Jasicka-Misiak, Izabela; Poliwoda, Anna; Kazakova, Anastasia; Godovan, Vladlena V; Halama, Marek; Wieczorek, Piotr P | 2015 | Toxins |
| Effect of psilocin on extracellular dopamine and serotonin levels in the mesoaccumbens and mesocortical pathway in awake rats. | Sakashita, Yuichi; Abe, Kenji; Katagiri, Nobuyuki; Kambe, Toshie; Saitoh, Toshiaki; Utsunomiya, Iku; Horiguchi, Yoshie; Taguchi, Kyoji | 2015 | Biological & pharmaceutical bulletin |
| Sex differences and serotonergic mechanisms in the behavioural effects of psilocin. | Tyls, Filip; Palenicek, Tomas; Kaderabek, Lukas; Lipski, Michaela; Kubesova, Anna; Horacek, Jiri | 2016 | Behavioural pharmacology |

| | | | |
|---|---|---|---|
| Psilocin and ketamine microdosing: effects of subchronic intermittent microdoses in the elevated plus-maze in male Wistar rats. | Horsley, Rachel R; Palenicek, Tomas; Kolin, Jan; Vales, Karel | 2018 | Behavioural pharmacology |
| Alteration of Depressive-like Behaviors by Psilocybe cubensis Alkaloid Extract in Mice: the Role of Glutamate Pathway | Mahmoudi, E.; Faizi, M.; Hajiaghaee, R.; Razmi, A. | 2018 | Research Journal of Pharmacognosy |
| Harnessing psilocybin: Synaptic mechanisms underlying the antidepressant response | Thompson S. | 2019 | Neuropsychopharmacology |
| The polypharmacological profile of psilocybin and potential behavioural effects of very low doses | Kiilerich, K.; Speth, N.; Lorenz, J.; Casado-Sainz, A.; Shalgunov, V.; Lange, D.; Xiong, M.; Herth, M. M.; Overgaard, A.; Hansen, H. D.; Palner, M. | 2019 | European Neuropsychopharmacology |
| Psilocybin lacks antidepressant-like effect in the Flinders Sensitive Line rat. | Jefsen, Oskar; Hojgaard, Kristoffer; Christiansen, Sofie Laage; Elfving, Betina; Nutt, David John; Wegener, Gregers; Muller, Heidi Kaastrup | 2019 | Acta neuropsychiatrica |
| Effects of acute and repeated treatment with serotonin 5-$HT_{2A}$ receptor agonist hallucinogens on intracranial self-stimulation in rats. | Sakloth, Farhana; Leggett, Elizabeth; Moerke, Megan J; Townsend, E Andrew; Banks, Matthew L; Negus, S Stevens | 2019 | Experimental and clinical psychopharmacology |

| | | | |
|---|---|---|---|
| The Effects of Psilocybin on Binge-Like Feeding Behaviour in Rats | Hurley, S.; Gilmour, G.; Soula, A.; Medhurst, L.; Hickey, M.; Whelan, T.; Selimbeyoglu, A. | 2020 | Neuropsychopharmacology |
| Effects of a single dose of psilocybin on behaviour, brain 5-$HT_{2A}$ receptor occupancy and gene expression in the pig. | Donovan, Lene Lundgaard; Johansen, Jens Vilstrup; Ros, Nidia Fernandez; Jaberi, Elham; Linnet, Kristian; Johansen, Sys Stybe; Ozenne, Brice; Issazadeh-Navikas, Shohreh; Hansen, Hanne Demant; Knudsen, Gitte Moos | 2020 | European Neuropsychopharmacology |
| Psychedelics, but Not Ketamine, Produce Persistent Antidepressant-like Effects in a Rodent Experimental System for the Study of Depression. | Hibicke, Meghan; Landry, Alexus N; Kramer, Hannah M; Talman, Zoe K; Nichols, Charles D | 2020 | ACS chemical neuroscience |
| One Dose of Psilocybin in Late Adolescence Mitigates Deleterious Effects of Developmental Stress on Cognition and Behavioral Despair in Adult Female Rats | Hibicke, M.; Nichols, C. | 2020 | Faseb Journal |
| Harnessing Psilocybin: Antidepressant-Like Behavioral and Synaptic Actions of Psilocybin are Independent of 5-HT2R Activation in Mice | Hesselgrave, N.; Troppoli, T.; Wulff, A.; Cole, A.; Thompson, S. | 2020 | Neuropsychopharmacology |

| | | | |
|---|---|---|---|
| Psilocybin and Ketamine Acutely Promote Wakefulness, Suppress REM Sleep but Differentially Modulate High Frequency EEG Oscillatory Power in Wistar Kyoto Rats - a Preliminary Analysis | Thomas, C.; Gilmour, G.; Medhurst, L.; Soula, A.; Hickey, M.; Hurley, S.; Whelan, T.; Selimbeyoglu, A. | 2020 | Neuropsychopharmacology |
| Psilocybin, After Only a Single Treatment, has Persistent Antidepressant-Like Effects in a Rat Experimental System for the Study of Mood Disorders | Nichols, C.; Hibicke, M. | 2020 | Neuropsychopharmacology |
| Low Doses of Psilocybin and Ketamine Enhance Motivation and Attention in Poor Performing Rats: Evidence for an Antidepressant Property. | Higgins, Guy A; Carroll, Nicole K; Brown, Matt; MacMillan, Cam; Silenieks, Leo B; Thevarkunnel, Sandy; Izhakova, Julia; Magomedova, Lilia; DeLannoy, Ines; Sellers, Edward M | 2021 | Frontiers in pharmacology |
| A Complex Impact of Systemically Administered 5-$HT_{2A}$ Receptor Ligands on Conditioned Fear. | Hagsater, Sven Melker; Pettersson, Robert; Pettersson, Christopher; Atanasovski, Daniela; Naslund, Jakob; Eriksson, Elias | 2021 | The international journal of neuropsychopharmacology |
| Investigation of the Structure-Activity Relationships of Psilocybin Analogues | Klein A.K.; Chatha M.; Laskowski L.J.; Anderson E.I.; Brandt S.D.; Chapman S.J.; McCorvy J.D.; Halberstadt A.L. | 2021 | ACS Pharmacology and Translational Science |
| Harnessing psilocybin: antidepressant-like behavioral and synaptic actions of psilocybin are independent of 5-HT2R activation in mice. | Hesselgrave, Natalie; Troppoli, Timothy A; Wulff, Andreas B; Cole, Anthony B; Thompson, Scott M | 2021 | Proceedings of the National Academy of Sciences of the United States of America |

List of Excluded Studies by Date, BIPA

| Title | Authors | Pub Year | Journal | Notes |
|---|---|---|---|---|
| Pharmacology of psilocybin, a drug from psilocybe mexicana heim | Weidmann H.; Taeschler M.; Konzett H. | 1958 | Experientia | Exclusion reason: not available in English; |
| The effect of the action of psilocybin on the rabbit brain | Monnier M. | 1959 | Experientia | Exclusion reason: not available in English |
| Studies on psilocybin and related compounds. I. Communication. Structure/activity relationship of oxyindole-derivatives with regard to their effect on the knee jerk of spinal cats. | WEIDMANN, H; CERLETTI, A | 1960 | Helvetica physiologica et pharmacologica acta | Exclusion reason: No behaviour measured |
| The relationship between the metabolic fate and pharmacological actions of serotonin, bufotenine and psilocybin | Gessner, P. K.; Khairallah, P. A.; McIsaac, W. M.; Page, L. H. | 1960 | Jour Pharmacol and Exptl Therap | Exclusion reason: No behaviour measured |
| EFFECT OF PSILOCYBIN UPON SYSTEMIC, PULMONARY, AND CORONARY CIRCULATION OF INTACT DOG | MAXWELL, GM; KNEEBONE, GM; ELLIOTT, RB | 1962 | ARCHIVES INTERNATIONALES DE PHARMACODYNAMIE ET DE THERAPIE | Exclusion reason: Unavailable within timeframe of study |

| | | | | |
|---|---|---|---|---|
| DEPHOSPHORYLATION OF PSILOCYBIN IN INTACT MOUSE | HORITA, A; WEBER, LJ | 1962 | TOXICOLOGY AND APPLIED PHARMACOLOGY | Exclusion reason: No behaviour measured |
| FATE OF PSILOCIN IN RAT | KALBERER, F; RUTSCHMANN, J; KREIS, W | 1962 | BIOCHEMICAL PHARMACOLOGY | Exclusion reason: No behavioural/neurological outcome |
| EFFECTS OF LSD-25, PSILOCYBIN, AND PSILOCIN ON TEMPORAL LOBE EEG PATTERNS AND LEARNED BEHAVIOR IN CAT | ADEY, WR; BELL, FR; DENNIS, BJ | 1962 | NEUROLOGY | Exclusion reason: Unavailable within timeframe of study |
| Effects of LSD-25, psiolocybin, and psilocin on temporal lobe EEG patterns and learned behavior in the cat. | ADEY, W R; BELL, F R; DENNIS, B J | 1962 | Neurology | Exclusion reason: Could not be found after ILLs |
| Search for experimental equivalents of hallucinogenic activity. Effects of hallucinogens on rectal temperature in rabbit and on morphine effects in mouse | Jacob J.; Lafille C.; Loiseau G.; Echinard-Garin P.; Barthelemy C. | 1962 | | Exclusion reason: Could not be found after ILLs |
| The effect of psilocybin upon the systemic, pulmonary and coronary circulation of the intact dog | Maxwell G.M.; Kneebone G.M.; Elliott R.B. | 1962 | Archives internationales de pharmacodynamie et de therapie | Exclusion reason: No behaviour measured |
| The late of Psilocin in the rat | Kalberer, F.; Kreis, W.; Rutschmann, J. | 1962 | Biochem Pharmacol | Exclusion reason: No behaviour measured |

| | | | | |
|---|---|---|---|---|
| SIMULTANEOUS STUDIES OF BLOOD SUGAR, BEHAVIOURAL CHANGES AND EEG ON WAKE RABBIT AFTER ADMININISTRATION OF PSILOCYBIN | STEINER, JE; SULMAN, FG | 1963 | ARCHIVES INTERNATIONALES DE PHARMACODYNAMIE ET DE THERAPIE | Exclusion reason: Unavailable within timeframe of study |
| SIMULTANEOUS STUDIES OF BLOOD SUGAR, BEHAVIOURAL CHANGES AND EEG ON WAKE RABBIT AFTER ADMININISTRATION OF PSILOCYBIN | Steiner, J. E.; Sulman, F. G. | 1963 | Archives Internationales De Pharmacodynamie Et De Therapie | Exclusion reason: Could not be found after ILLs |
| Some biochemical studies on psilocybin and psilocin | Horita, A. | 1963 | Jour Neuropsychiat | Exclusion reason: No behaviour measured |
| PSYCHOTOMIMETICS--A NEUROPHARMACOLOGICAL STUDY. | CURTIS, D R | 1963 | Proceedings of the Australian Association of Neurologists | Exclusion reason: not original research |
| PHARMACOLOGICAL DETECTION and CHARACTERIZATION of HALLUCINOGENS. I. HYPERTHERMIZING ACTIVITIES in THE RABBIT | Jacob J.; Lafille C. | 1963 | Archives internationales de pharmacodynamie et de therapie | Exclusion reason: not available in English |
| An electrographic study of psilxcin and 4-methyl-alpha-methyl tryptamine (mp-809) | Brodey J.F.; Steiner W.G.; Himwich H.E. | 1963 | | Exclusion reason: No behaviour measured |

| | | | | |
|---|---|---|---|---|
| Research on the pharmacological characterization and differentiation of hallucinogenic drugs (indole derivatives and mescaline, nalorphine, central anticholinergics, and phencyclidine). | Jacob, J; Lafille, C; Loiseau, G; Echinard-Garin, P; Barthelemy, C | 1964 | L'Encephale: Revue de psychiatrie clinique biologique et therapeutique | Exclusion reason: not available in English |
| DISINHIBITION OF CONDITIONED BEHAVIOR BY CEREBRAL SYNAPTIC INHIBITORS | Halasz, M. F.; Marrazzi, A. S. | 1965 | Pharmacologist | Exclusion reason: No behaviour measured |
| The effects of tryptamine and alpha-methyl-tryptamine on the general and coronary haemodynamics and metabolism of the dog | Maxwell, G. M.; Burnell, R. H.; Kneebone, G. M. | 1965 | Arch Int Pharmacodyn Therap | Exclusion reason: No behaviour measured |
| EFFECTS OF STRESS AND PSYCHOTROPIC DRUGS ON RAT LIVER TRYPTOPHAN PYRROLASE. | NOMURA, J | 1965 | Endocrinology | Exclusion reason: No behaviour measured |
| The influence of hallucinogenic drugs upon in vivo brain levels of adenine nucleotides, phosphocreatine and inorganic phosphate in the rat. | Lewis, J J; Ritchie, A P; van Petten, G R | 1965 | British journal of pharmacology and chemotherapy | Exclusion reason: No behaviour measured |
| PSILOCIN - EFFECTS ON BEHAVIOUR AND BRAIN SEROTONIN IN MICE | COLLINS, RL; ORDY, JM; SAMORAJS.T | 1966 | NATURE | Exclusion reason: duplicate |

| | | | | |
|---|---|---|---|---|
| EFFECT OF PSILOCYBINE ON VISUAL EVOKED POTENTIALS IN A CERCOPITHECINAE PAPIO PAPIO | BERMOND, F; BERT, J; AYATS, J | 1966 | COMPTES RENDUS DES SEANCES DE LA SOCIETE DE BIOLOGIE ET DE SES FILIALES | Exclusion reason: not available in English |
| Inhibition of isolation-induced attack behavior of mice by drugs. | Uyeno, E T | 1966 | Proceedings of the Western Pharmacology Society | Exclusion reason: not original research |
| The modification of the EEG in the reserpinized rabbit caused by psychoanaleptics and the precursors of biogenic amines Ger., and Engl. summ. | Nakajima, H.; Thuillier, J. | 1966 | Med Pharmacol Exp | Exclusion reason: No behaviour measured |
| Eeg effects of psychoanaleptics and biogenic amine precursors in reserpinized rabbits | Kakajirna H.; Thuillier J. | 1966 | | Exclusion reason: No behaviour measured |
| EFFECT OF PSILOCYBINE ON VISUAL EVOKED POTENTIALS IN A CERCOPITHECINAE PAPIO PAPIO | Bermond, F.; Bert, J.; Ayats, J. | 1966 | Comptes Rendus Des Seances De La Societe De Biologie Et De Ses Filiales | Exclusion reason: not available in English |
| Effect of certain psychopharmacologic substances on S35-methionine cytoplasm incorporation (cyrillic) | Petkov V.; Shoumkov G.; Koushev V. | 1966 | Savremenna Medicina | Exclusion reason: No behaviour measured |
| EFFECTS OF CHLORPROMAZINE AND PSILOCIN ON PREGNANCY OF C57BL/10 MICE AND THEIR OFFSPRING AT BIRTH | ROLSTEN, C | 1967 | ANATOMICAL RECORD | Exclusion reason: Unavailable within timeframe of study |

| | | | | |
|---|---|---|---|---|
| EFFECTS OF MESCALINE AND PSILOCYBIN ON DOMINANCE BEHAVIOR OF RAT | UYENO, ET | 1967 | ARCHIVES INTERNATIONALES DE PHARMACODYNAMIE ET DE THERAPIE | Exclusion reason: Unavailable within timeframe of study; |
| EFFECT OF PSILOCYBIN ON EYE MOVEMENTS AND ON ELECTRORETINOGRAM IN A CERCOPITHECINAE PAPIO PAPIO | BERMOND, F; BERT, J; AYATS, H | 1967 | COMPTES RENDUS DES SEANCES DE LA SOCIETE DE BIOLOGIE ET DE SES FILIALES | Exclusion reason: not available in English |
| EFFECTS OF CHLORPROMAZINE AND PSILOCIN ON PREGNANCY OF C57BL/10 MICE AND THEIR OFFSPRING AT BIRTH | Rolsten, C. | 1967 | Anatomical Record | Exclusion reason: No behaviour measured |
| Hallucinogens and dominance behavior of the rat. | Uyeno, E T | 1967 | Proceedings of the Western Pharmacology Society | Exclusion reason: not original research |
| A possible correlation between drug-induced hallucinations in man and a behavioural response in mice. | Corne, S J; Pickering, R W | 1967 | Psychopharmacologia | Exclusion reason: Competitive |
| Inhibitory effects of lysergic acid derivatives and reserpine on 5-HT binding to nerve ending particles. | Marchbanks, R M | 1967 | Biochemical pharmacology | Exclusion reason: No behaviour measured |

| | | | | |
|---|---|---|---|---|
| EFFECT OF MONO AMINO OXIDASE INHIBITORS ON EXPERIMENTAL PSYCHOSES FOLLOWING THE ADMINISTRATION OF PSILOCYBIN | Vojtechovsky, M.; Hort, V.; Safratove, V. | 1968 | Activitas Nervosa Superior | Exclusion reason: Could not be found after ILLs |
| TRYPTAMINE RECEPTORS IN DOG SPINAL CORD AND THEIR RELATIONSHIP TO THE AGONISTIC ACTIONS OF LYSERGIC-ACID DI ETHYLAMIDE LIKE PSYCHOTOGENS | Martin, W. R.; Eades, C. G. | 1968 | Pharmacologist | Exclusion reason: No behaviour measured |
| DIFFERENTIATION OF INST ELECTRO ENCEPHALOGRAM DEACTIVATION NICOTINE CENT STIM LYSERGIC-ACID DI ETHYLAMIDE CENT STIM PSILOCYBIN CENT STIM RABBIT | Kakolewski, J. W. | 1968 | Physiology and Behavior | Exclusion reason: No behaviour measured |
| Interactions of norepinephrine with subcellular fractions of rat brain. L characteristics of norepinephrine uptake | Herblin W.F.; O'Brien R.D. | 1968 | Brain Research | Exclusion reason: No behaviour measured |
| Autoradiographic studies on the distribution of psychoactive drugs in the rat brain: III. 14C psilocin. | Hopf, A; Eckert, H | 1969 | Psychopharmacologia | Exclusion reason: No behaviour measured |
| EFFECTS OF PSILOCYBIN, DIMETHYLTRYPTAMINE AND VARIOUS LYSERGIC ACID DERIVATIVES ON PHOTICALLY- | MELDRUM, BS; NAQUET, R | 1970 | BRITISH JOURNAL OF PHARMACOLOGY | Exclusion reason: duplicate |

| | | | | |
|---|---|---|---|---|
| INDUCED EPILEPSY IN BABOON (PAPIO-PAPIO) | | | | |
| EFFECT OF LSD-25 AND PSILOCYBIN ON NOREPINEPHRINE AND 3H-NOREPINEPHRINE METABOLITES IN RAT BRAIN | GOLDSTEI.ML; LOVELL, RA; BOGGAN, WO | 1970 | FEDERATION PROCEEDINGS | Exclusion reason: Unavailable within timeframe of study |
| ACTION OF SOME NEURO DRUGS AND PSYCHO PHARMACA DRUGS ON MEMBRANE ATPASE AND ACETYL CHOLIN ESTERASE OF CORTICAL SYNAPTOSOMES | Waser, P. G.; Schaub, E. | 1970 | Heilbronn, Edith and Anders Winter | Exclusion reason: No behaviour measured |
| EFFECTS OF PSILOCYBIN, DIMETHYLTRYPTAMINE AND VARIOUS LYSERGIC ACID DERIVATIVES ON PHOTICALLY-INDUCED EPILEPSY IN BABOON (PAPIO-PAPIO) | Meldrum, B. S.; Naquet, R. | 1970 | British Journal of Pharmacology | Exclusion reason: duplicate |
| The action of tryptamine on the dog spinal cord and its relationship to the agonistic actions of LSD-like psychotogens. | Martin, W R; Eades, C G | 1970 | Psychopharmacologia | Exclusion reason: Competitive; |

| | | | | |
|---|---|---|---|---|
| EFFECT OF LSD-25 AND PSILOCYBIN ON NOREPINEPHRINE AND 3H-NOREPINEPHRINE METABOLITES IN RAT BRAIN | Goldstei.Ml,; Lovell, R. A.; Boggan, W. O. | 1970 | Federation Proceedings | Exclusion reason: No behaviour measured |
| EFFECTS OF HALLUCINOGENS ON RAT BRAIN MONO AMINE OXIDASE ACTIVITY | Collins, B. J.; Lovell, R. A.; Boggan, W. O.; Freedman, D. X. | 1970 | Pharmacologist | Exclusion reason: No behaviour measured |
| PSYCHOTOMIMETIC DRUGS AND BRAIN 5 HYDROXY TRYPTAMINE METABOLISM | Freedman, D. X.; Gottlieb, R.; Lovell, R. A. | 1970 | Biochemical Pharmacology | Exclusion reason: No behaviour measured |
| EFFECTS OF PSILOCYBIN, DIMETHYLTRYPTAMINE, MESCALINE AND VARIOUS LYSERGIC ACID DERIVATIVES ON EEG AND ON PHOTICALLY INDUCED EPILEPSY IN BABOON (PAPIO-PAPIO) | MELDRUM, BS; NAQUET, R | 1971 | ELECTROENCEPHALOGRAPHY AND CLINICAL NEUROPHYSIOLOGY | Exclusion reason: duplicate |
| THE ACTION OF SOME NEURO PHARMACOLOGICAL AND PSYCHO PHARMACOLOGICAL AGENTS ON THE MEMBRANE ATPASE OF CORTICAL SYNAPTOSOMES | Waser, P. G.; Schaub, E. | 1971 | Clementi, F. And B. Ceccarelli | Exclusion reason: No behaviour measured |

| | | | | |
|---|---|---|---|---|
| EFFECT OF PSYCHOTOMIMETIC DRUGS ON THE CYCLIC 3 5 AMP SYSTEM OF RAT BRAIN | Uzunov, P.; Weiss, B. | 1971 | Pharmacologist | Exclusion reason: No behaviour measured |
| Effects of psilocybin, dimethytaryptamine, mescaline and various lysergic acid derivates on the eeg and on photically induced epilepsy in the baboon (papio papio) | Meldrum B.S.; Naquet R. | 1971 | | Exclusion reason: duplicate |
| Stimulation of [14C]serotonin synthesis from [ 14C]tryptophan by mescaline in rat pineal organ cultures | Shein H.M.; Wilson S.; Larin F.; Wurtman R.J. | 1971 | Life Sciences | Exclusion reason: duplicate |
| Effects of psychoactive agents on the conditioning of the microcirculation in the rat. | Kato, L; Gozsy, B; Ban, T A; Sterlin, C | 1971 | Conditional reflex | Exclusion reason: No behaviour measured |
| Hallucinogenic drugs of the indolealkylamine type and central monoamine neurons. | Anden, N E; Corrodi, H; Fuxe, K | 1971 | The Journal of pharmacology and experimental therapeutics | Exclusion reason: No non-drug control |
| Stimulation of 14clerotonin synthesis from 14chryptophan by mescaline in rat pineal organ cultures | Shein H.M.; Wilson S.; Larin F.; Wurtman R.J. | 1971 | Life Sciences | Exclusion reason: No behaviour measured |
| EFFECTS OF SEROTONIN 5 HYDROXY TRYPTAMINE AND SOME RELATED INDOLE COMPOUNDS IN A MAMMALIAN SYMPATHETIC GANGLION | Haefely, W. | 1972 | Experientia (Basel) | Exclusion reason: No behaviour measured |

| | | | | |
|---|---|---|---|---|
| THE EFFECTS OF VARIOUS ERGOT DERIVATIVES ON THE ELECTRO ENCEPHALOGRAM AND PHOTO SENSITIVITY IN PAPIO-PAPIO | Balzano, E.; Meldrum, B. S.; Naquet, R.; Vuillon-Cacciuttolo, G. | 1972 | Electroencephalography and Clinical Neurophysiology | Exclusion reason: No psilocybin/psilocin |
| EFFECTS OF LSD, PSILOCYBIN, HARMALINE AND AMPHETAMINE ON BODY-TEMPERATURE OF PARA-CHLOROPHENYLALANINE PRETREATED RATS | LADEFOGED, O | 1973 | ARCHIVES INTERNATIONALES DE PHARMACODYNAMIE ET DE THERAPIE | Exclusion reason: Unavailable within timeframe of study |
| EFFECTS OF AMPHETAMINE (A), LSD, PSILOCYBIN (P), AND DOM ON SCHEDULE-CONTROLLED BEHAVIOR IN RAT | MARQUIS, WJ; TILSON, HA; RECH, RH | 1973 | FEDERATION PROCEEDINGS | Exclusion reason: Unavailable within timeframe of study;; |
| The effects of d-amphetamine, 2,5-dimethoxy-4-methyl-amphetamine (DOM), and psilocybin on fixed-interval responding in the rat. | Tilson, Hugh A; Marquis, William J; Rech, R. H | 1973 | Proceedings of the Annual Convention of the American Psychological Association | Exclusion reason: Could not be found after ILLs |
| The effects of LSD, psilocybin, harmaline and amphetamine on the body temperature of para-chlorophenylalanine pretreated rats. | Ladefoged, O | 1973 | Archives internationales de pharmacodynamie et de therapie | Exclusion reason: No behaviour measured |

| | | | | |
|---|---|---|---|---|
| EFFECT OF LSD, PSILOCYBIN, HARMALINE AND AMPHETAMINE ON BODY-TEMPERATURE OF PARA-CHLOROPHENYLALANINE PRETREATED RABBITS | LADEFOGED, O | 1974 | ARCHIVES INTERNATIONALES DE PHARMACODYNAMIE ET DE THERAPIE | Exclusion reason: duplicate |
| EFFECTS OF PSILOCYBIN ON EEG AND BEHAVIOR IN MONKEYS | HORIBE, M | 1974 | ACTIVITAS NERVOSA SUPERIOR | Exclusion reason: Unavailable within timeframe of study |
| EFFECTS OF LSD MESCALINE AND PSILOCYBIN ON SYMPATHETIC PREGANGLIONIC NEURONS | McCloskey, K. L.; Franz, D. N. | 1974 | Pharmacologist | Exclusion reason: No behaviour measured |
| DIFFERENT EFFECTS OF PSILOCYBIN ON THE ELECTRO ENCEPHALOGRAM OF VARIOUS TYPES OF EPILEPSY | Kolarik, J. | 1974 | Electroencephalography and Clinical Neurophysiology | Exclusion reason: No behaviour measured |
| Distribution patterns of -sup-1-sup-4-psilocybin in the brains of various animals. | Hopf, A; Eckert, H | 1974 | Activitas Nervosa Superior | Exclusion reason: No behaviour measured |
| A comparison of psychotomimetic drug effects on rat brain norepinephrine metabolism. | Stolk, J M; Barchas, J D; Goldstein, M; Boggan, W; Freedman, D X | 1974 | The Journal of pharmacology and experimental therapeutics | Exclusion reason: No behaviour measured |

| | | | | |
|---|---|---|---|---|
| The possible role of tryptamine in brain function and its relationship to the actions of LSD-like hallucinogens. | Martin, W R; Sloan, J W | 1974 | The Mount Sinai journal of medicine, New York | Exclusion reason: No psilocybin/psilocin |
| The effect of LSD, psilocybin, harmaline and amphetamine on the body temperature of para-chlorophenylalanine pretreated rabbits. | Ladefoged, O | 1974 | Archives internationales de pharmacodynamie et de therapie | Exclusion reason: No behaviour measured |
| CORTICAL AND OR SUBCORTICAL EFFECTS AS A FUNCTION OF HALLUCINOGENIC DRUG STRUCTURE | Goldman, H.; Fischer, R. | 1974 | Pharmacologist | Exclusion reason: No psilocybin/psilocin; |
| The effects of 5-hydroxytryptamine and some related compounds on the cat superior cervical ganglion in situ. | Haefely, W | 1974 | Naunyn-Schmiedeberg's archives of pharmacology | Exclusion reason: No behaviour measured |
| Comparison of the discriminative stimulus properties of delta9-THC and psilocybin in rats. | Greenberg, I; Kuhn, D; Appel, J B | 1975 | Pharmacology, biochemistry, and behavior | Exclusion reason: Drug discrimination |
| The dopamine receptor: differential binding of d LSD and related agents to agonist and antagonist states | Creese I.; Burt D.R.; Snyder S.H. | 1975 | Life Sciences | Exclusion reason: No behaviour measured |
| Serotonin sensitive adenylate cyclase activity in immature rat brain | Von Hungen K.; Roberts S.; Hill D.F. | 1975 | Brain Research | Exclusion reason: No behaviour measured |

| | | | | |
|---|---|---|---|---|
| Interactions between lysergic acid diethylamide and dopamine-sensitive adenylate cyclase systems in rat brain. | Hungen, K V; Roberts, S; Hill, D F | 1975 | Brain research | Exclusion reason: No behaviour measured |
| Hallucinogenic indoleamines: Preferential action upon presynaptic serotonin receptors. | Aghajanian, G K; Hailgler, H J | 1975 | Psychopharmacology communications | Exclusion reason: No behaviour measured |
| DISCRIMINATIVE RESPONSE CONTROL BY PSYCHO MOTOR STIMULANTS | Silverman, P. B.; Ho, B. T. | 1976 | Psychopharmacology Communications | Exclusion reason: Drug discrimination |
| THE BABOON PAPIO-PAPIO HUMAN EPILEPSY MODEL PHARMACOLOGICAL ASPECTS | Balzamo, E. | 1976 | Stal | Exclusion reason: Could not be found after ILLs |
| DRUGS AND PONTO GENICULO OCCIPITAL WAVES IN THE LATERAL GENICULATE BODY OF THE CURARIZED CAT PART 2 PONTO GENICULO OCCIPITAL WAVE ACTIVITY AND BRAIN 5 HYDROXY TRYPTAMINE | Ruch-Monachon, M. A.; Jalfre, M.; Haefely, W. | 1976 | Archives Internationales De Pharmacodynamie Et De Therapie | Exclusion reason: No behaviour measured |
| Binding interactions of lysergic acid diethylamide and related agents with dopamine receptors in the brain | Burt D.R.; Creese I.; Snyder S.H. | 1976 | Molecular Pharmacology | Exclusion reason: No behaviour measured |
| PSILOCYBIN - BIPHASIC DOSE-RESPONSE EFFECTS ON ACOUSTIC STARTLE REFLEX IN RAT | DAVIS, M; WALTERS, JK | 1977 | PHARMACOLOGY BIOCHEMISTRY AND BEHAVIOR | Exclusion reason: duplicate |

| | | | | |
|---|---|---|---|---|
| Effect of indole hallucinogens, mescaline and DMPEA on rat plasma prolactin | Meltzer H.Y.; Fessler R.G.; Simonovic M.; Fang V.S. | 1977 | Federation Proceedings | Exclusion reason: No behaviour measured |
| EFFECTS OF MESCALINE AND PSILOCIN ON ACQUISITION, CONSOLIDATION, AND PERFORMANCE OF LIGHT-DARK DISCRIMINATION IN 2 INBRED STRAINS OF MICE | CASTELLANO, C | 1978 | PSYCHOPHARMACOLOGY | Exclusion reason: duplicate ; |
| LSD-LIKE HALLUCINOGENS IN DOG - VALIDATION STUDIES WITH MESCALINE (MES), PSILOCIN (PSI) AND DIMETHYLTRYPTAMINE (DMT) | VAUPEL, DB; MARTIN, WR | 1978 | FEDERATION PROCEEDINGS | Exclusion reason: Drug discrimination |
| LSD-like hallucinogens in the dog: validation studies with mescaline (MES), psilocin (PSI) and dimethyltryptamine (DMT) | Vaupel D.B.; Martin W.R. | 1978 | Federation Proceedings | Exclusion reason: not original research |
| STIMULUS PROPERTIES OF COMMONALITY WITH OTHER HALLUCINOGENS 2 5 DI METHOXY-4-METHYL AMPHETAMINE | Silverman, P. B.; Ho, B. T. | 1978 | Psychopharmacology | Exclusion reason: Could not be found after ILLs |
| EFFECTS OF PSYCHO DYSLEPTICS ON AGGRESSIVE BEHAVIOR OF ANIMALS | Uyeno, E. T. | 1978 | Valzelli, Luigi | Exclusion reason: not original research |
| Defining the histamine H2-receptor in brain: the interaction with LSD. | Green, J P; Weinstein, H; Maayani, S | 1978 | NIDA research monograph | Exclusion reason: No behaviour measured |

| | | | | |
|---|---|---|---|---|
| LSD and related drugs as DA antagonists: Receptor-mediated effects on the synthesis and turnover of DA. | Persson, Sven-Ake | 1978 | Life Sciences | Exclusion reason: No behaviour measured |
| LSD AND RELATED DRUGS AS DOPAMINE ANTAGONISTS RECEPTOR MEDIATED EFFECTS ON THE SYNTHESIS AND TURNOVER OF DOPAMINE | Persson, S. A. | 1978 | Life Sciences | Exclusion reason: No behaviour measured |
| HIGH AFFINITY TRITIATED SEROTONIN BINDING TO CAUDATE INHIBITION BY HALLUCINOGENS AND SEROTONINERGIC DRUGS | Whitaker, P. M.; Seeman, P. | 1978 | Psychopharmacology | Exclusion reason: No behaviour measured |
| Selective labeling of serotonin receptors by d-[3H]lysergic acid diethylamide in calf caudate | Whitaker P.M.; Seeman P. | 1978 | Proceedings of the National Academy of Sciences of the United States of America | Exclusion reason: No behaviour measured |
| Stimulation of rat prolactin secretion by indolealkylamine hallucinogens. | Meltzer, H Y; Fessler, R G; Simonovic, M; Fang, V S | 1978 | Psychopharmacology | Exclusion reason: No behaviour measured |
| Stimulation of adenylate cyclase activity in monkey anterior limbic cortex by serotonin | Ahn H.S.; Makman M.H. | 1978 | Brain Research | Exclusion reason: No behaviour measured |

| | | | | |
|---|---|---|---|---|
| INHIBITION OF FOOD-INTAKE IN THE DOG BY LSD, MESCALINE, PSILOCIN, D-AMPHETAMINE AND PHENYLISOPROPYLAMINE DERIVATIVES | VAUPEL, DB; NOZAKI, M; MARTIN, WR; BRIGHT, LD; MORTON, EC | 1979 | LIFE SCIENCES | Exclusion reason: duplicate |
| COMPARISONS OF THE EFFECTS OF MESCAL BEAN ALKALOIDS WITH MESCALINE, N,N-DIMETHYLTRYPTAMINE (DMT), PSILOCYBIN, AMPHETAMINE, DELTA-9-TETRAHYDROCANNABINOL (THC), AND PENTOBARBITAL IN RATS | BOURN, WM; KELLER, WJ; BONFIGLIO, JF | 1979 | FEDERATION PROCEEDINGS | Exclusion reason: Unavailable within timeframe of study |
| PSILOCYBIN NEURO-PSYCHO-PHARMACOLOGY IN RABBITS | GUEST, J; CONSROE, P | 1979 | RESEARCH COMMUNICATIONS IN PSYCHOLOGY PSYCHIATRY AND BEHAVIOR | Exclusion reason: Unavailable within timeframe of study |
| COMPARISONS OF THE EFFECTS OF MESCAL BEAN ALKALOIDS WITH MESCALINE, N,N-DIMETHYLTRYPTAMINE (DMT), PSILOCYBIN, AMPHETAMINE, DELTA-9-TETRAHYDROCANNABINOL (THC), AND PENTOBARBITAL IN RATS | Bourn, W. M.; Keller, W. J.; Bonfiglio, J. F. | 1979 | Federation Proceedings | Exclusion reason: not original research |

| | | | | |
|---|---|---|---|---|
| EXPERIENCE OF INDUCED VISUAL HALLUCINATIONS DEPENDS ON PRE TREATMENT ELECTRO ENCEPHALOGRAM SPECTRA | Koukkou, M.; Lehmann, D. | 1979 | Neuroscience Letters | Exclusion reason: No mammals |
| HALLUCINOGENS ANTAGONIZE HISTAMINE H-1 RECEPTORS OF CULTURED MOUSE NEURO BLASTOMA CELLS | Fredrickson, P. A.; Richelson, E. | 1979 | European Journal of Pharmacology | Exclusion reason: No behaviour measured |
| Shape change of blood platelets: A model for cerebral 5-hydroxytryptamine receptors? | Graf M.; Pletscher A. | 1979 | British Journal of Pharmacology | Exclusion reason: No behaviour measured |
| INCREASE OF CYCLIC GMP IN BLOOD PLATELETS BY BIOGENIC AMINES A RECEPTOR MEDIATED EFFECT | Laubscher, A.; Pletscher, A. | 1980 | Journal of Pharmacy and Pharmacology | Exclusion reason: No behaviour measured |
| Pharmacological characteristics of tetrahydrocannabinol seizure susceptible rabbits | Consroe P.; Fish B.S. | 1980 | Federation Proceedings | Exclusion reason: Could not be found after ILLs |
| Hallucinogenic agents as discriminative stimuli: A correlation with serotonin receptor affinities | Glennon R.A.; Young R.; Rosecrans J.A.; Kallman M.J. | 1980 | Psychopharmacology | Exclusion reason: Drug discrimination |
| Serotonin receptors in hippocampus and frontal cortex | Seeman P.; Westman K.; Coscina D.; Warsh J.J. | 1980 | European Journal of Pharmacology | Exclusion reason: No behaviour measured |

| | | | | |
|---|---|---|---|---|
| Hallucinogens potentiate responses to serotonin and norepinephrine in the facial motor nucleus. | McCall, R B; Aghajanian, G K | 1980 | Life sciences | Exclusion reason: No behaviour measured |
| ELECTRO ENCEPHALOGRAPHIC EFFECTS OF HALLUCINOGENS AND CANNABINOIDS USING SLEEP WAKING BEHAVIOR AS A BASELINE | Fairchild, M. D.; Jenden, D. J.; Mickey, M. R.; Yale, C. | 1980 | Pharmacology Biochemistry and Behavior | Exclusion reason: No behaviour measured |
| 5 HYDROXY TRYPTAMINE DEPENDENT MYO CLONUS IN GUINEA-PIGS IS INDUCED BY 5 HYDROXY TRYPTAMINE AGONISTS CONTAINING AN INDOLE NUCLEUS BUT NOT BY THOSE POSSESSING A PIPERAZINE MOIETY | Luscombe, G.; Jenner, P.; Marsden, C. D. | 1981 | Neuroscience Letters | Exclusion reason: not original research |
| 5HT dependent myoclonus in guinea pigs is induced by 5HT agonists containing an indole nucleus, but not by those possessing a piperazine moiety | Luscombe G.; Jenner P.; Marsden C.D. | 1981 | Neuroscience Letters | Exclusion reason: not original research |
| Structure-activity relationships among hallucinogenic tryptamine derivatives evaluated by schedule-controlled behaviour | Adron Harris R.; Homfeld E.; Campaigne E. | 1981 | Journal of Pharmacy and Pharmacology | Exclusion reason: duplicate |

| | | | | |
|---|---|---|---|---|
| Psilocybin as a discriminative stimulus: lack of specificity in an animal behavior model for 'hallucinogens'. | Koerner, J; Appel, J B | 1982 | Psychopharmacology | Exclusion reason: Drug discrimination |
| Depolarizing responses recorded from nodose ganglion cells of the rabbit evoked by 5-hydroxytryptamine and other substances | Wallis D.I.; Stansfeld C.E.; Nash H.L. | 1982 | Neuropharmacology | Exclusion reason: No behaviour measured |
| Nerve terminal effects of indoleamine psychotomimetics on 5-hydroxytryptamine. | Halaris, A E | 1982 | Neuroscience and biobehavioral reviews | Exclusion reason: No behaviour measured |
| DOSE DEPENDENT BEHAVIORAL-CHANGES INDUCED BY PSILOCYBIN IN SELECTED MEMBERS OF A PRIMATE SOCIAL COLONY | SINK, CA; BELUHAN, LJ; SCHLEMMER, RF; HEINZE, WJ; DAVIS, JM | 1983 | FEDERATION PROCEEDINGS | Exclusion reason: Unavailable within timeframe of study |
| DIFFERENTIAL EFFECTS OF INDOLEAMINE HALLUCINOGENS ON SEROTONIN CONTAINING NEURONS IN THE NUCLEUS CENTRALIS SUPERIOR AND NUCLEUS RAPHE PALLIDUS IN FREELY MOVING CATS | Trulson, V. M.; Trulson, M. E. | 1983 | Federation Proceedings | Exclusion reason: No behaviour measured |

| | | | | |
|---|---|---|---|---|
| INTERACTION OF 5 HYDROXY TRYPTAMINE AGONISTS WITH GUINEA-PIG BRAIN STEM 5 HYDROXY TRYPTAMINE RECEPTORS AND THE INDUCTION OF MYO CLONUS | Jenner, P.; Luscombe, G.; Marsden, C. D. | 1983 | British Journal of Pharmacology | Exclusion reason: No behaviour measured |
| Inhibition of synaptosomal neurotransmitter uptake by hallucinogens | Whipple M.R.; Reinecke M.G.; Gage F.H. | 1983 | Journal of Neurochemistry | Exclusion reason: No behaviour measured |
| Correlation of [3H]5-hydroxytryptamine (5HT) binding to brain stem preparations and the production and prevention of myoclonus in guinea pig by 5HT agonists and antagonists | Luscombe G.; Jenner P.; Marsden C.D. | 1984 | European Journal of Pharmacology | Exclusion reason: No behaviour measured |
| Pharmacological characterization of solubilized 5-HT1 serotonin binding sites from bovine brain | Allgren R.L.; Kyncl M.M.A.; Ciaranello R.D. | 1985 | Brain Research | Exclusion reason: No behaviour measured |
| RELATIONSHIP OF CENTRAL NERVOUS SYSTEM TRYPTAMINERGIC PROCESSES AND THE ACTION OF LSD-LIKE HALLUCINOGENS | Martin, W. R.; Sloan, J. W. | 1986 | Pharmacology Biochemistry and Behavior | Exclusion reason: not original research |
| Relationship of CNS tryptaminergic processes and the action of LSD-like hallucinogens. | Martin, W R; Sloan, J W | 1986 | Pharmacology, biochemistry, and behavior | Exclusion reason: No non-drug control |

| | | | | |
|---|---|---|---|---|
| Differences in the stimulus properties of 3,4-methylenedioxyamphetamine and 3,4-methylenedioxymethamphetamine in animals trained to discriminate hallucinogens from saline. | Callahan, P M; Appel, J B | 1988 | The Journal of pharmacology and experimental therapeutics | Exclusion reason: No non-drug control |
| DIFFERENTIAL INTERACTIONS OF INDOLEALKYLAMINES WITH 5 HYDROXYTRYPTAMINE RECEPTOR SUBTYPES | McKenna, D. J.; Repke, D. B.; Lo, L.; Peroutka, S. J. | 1990 | Neuropharmacology | Exclusion reason: No behaviour measured |
| Lysergic acid diethylamide (LSD) administration selectively downregulates serotonin2 receptors in rat brain. | Buckholtz, N S; Zhou, D F; Freedman, D X; Potter, W Z | 1990 | Neuropsychopharmacology : official publication of the American College of Neuropsychopharmacology | Exclusion reason: No behaviour measured |
| Effect of ring fluorination on the pharmacology of hallucinogenic tryptamines. | Blair, J B; Kurrasch-Orbaugh, D; Marona-Lewicka, D; Cumbay, M G; Watts, V J; Barker, E L; Nichols, D E | 2000 | Journal of medicinal chemistry | Exclusion reason: No psilocybin/psilocin |

| | | | | |
|---|---|---|---|---|
| The effect of Psilocybe cubensis extract on hippocampal neurons in vitro. | Moldavan, M G; Grodzinskaya, A A; Solomko, E F; Lomberh, M L; Wasser, S P; Storozhuk, V M | 2001 | Fiziolohichnyi zhurnal (Kiev, Ukraine : 1994) | Exclusion reason: No behaviour measured |
| 5-$HT_{2A}$ receptor-stimulated phosphoinositide hydrolysis in the stimulus effects of hallucinogens. | Rabin, Richard A; Regina, Meredith; Doat, Mireille; Winter, J C | 2002 | Pharmacology, biochemistry, and behavior | Exclusion reason: No behaviour measured |
| The indoleamine hallucinogens, psilocin and 5MeODMT, increase prepulse inhibition in mice: role of 5 - HT1A and 5 - HT2A receptors | Powell, S. B.; Risbrough, V. B.; Lehmann-Masten, V. D.; Krebs-Thomson, K.; Hen, R.; Gingrich, J. A.; Geyer, M. A. | 2003 | Society for Neuroscience Abstract Viewer and Itinerary Planner | Exclusion reason: Could not be found after ILLs |
| Cognitive and psychopathological aspects of the 5-$HT_{2A}$ model of experimental psychosis | Hasler, F.; Dobricki, M.; Grimber, U.; Vollenweider, F. X. | 2003 | Journal of Psychopharmacology | Exclusion reason: Human trial |

| | | | | |
|---|---|---|---|---|
| Serotonin 5-hydroxytryptamine2A receptor-coupled phospholipase C and phospholipase A2 signaling pathways have different receptor reserves | Kurrasch-Orbaugh D.M.; Watts V.J.; Barker E.L.; Nichols D.E. | 2003 | Journal of Pharmacology and Experimental Therapeutics | Exclusion reason: No behaviour measured |
| The influence of psilocin and phenylethylamine on the energy metabolism in the rat heart | Machoy-Mokrzynska A.; Safranow K.; Borowiak K.S.; Majdanik S.; Dziedziejko V.; Bialecka M.; Chlubek D. | 2003 | Acta Toxicologica | Exclusion reason: No behaviour measured |
| Transient reinforcing effects of phenylisopropylamine and indolealkylamine hallucinogens in rhesus monkeys. | Fantegrossi, W E; Woods, J H; Winger, G | 2004 | Behavioural pharmacology | Exclusion reason: Drug discrimination; |
| Effects of systemic psilocin on serotonin release in rat brain | Katagiri, N.; Abe, K.; Yamaguchi, M.; Saitoh, T.; Horiguchi, Y.; Utsunomiya, I.; Hoshi, K.; Taguchi, K. | 2006 | Journal of Pharmacological Sciences | Exclusion reason: No behaviour measured |

| | | | | |
|---|---|---|---|---|
| Modeling of psychotic-like behavior: Comparison of MK-801 with psilocin, LSD, mescaline and 2C-B models. behavioral study on prepulse inhibition of acoustic startle and on locomotion | Palenicek, T.; Bubenikova, V.; Horacek, J. | 2006 | Schizophrenia Research | Exclusion reason: No non-drug control |
| Hallucinogens recruit specific cortical 5-HT(2A) receptor-mediated signaling pathways to affect behavior. | Gonzalez-Maeso, Javier; Weisstaub, Noelia V; Zhou, Mingming; Chan, Pokman; Ivic, Lidija; Ang, Rosalind; Lira, Alena; Bradley-Moore, Maria; Ge, Yongchao; Zhou, Qiang; Sealfon, Stuart C; Gingrich, Jay A | 2007 | Neuron | Exclusion reason: No psilocybin/psilocin |
| CONCENTRATION OF SELECTED MICROELEMENTS IN BLOOD SERUM OF RATS EXPOSED TO THE ACTION OF PSILOCIN AND PHENYLETHYLAMINE | Majdanik, Slawomir; Borowiak, Krzysztof; Brzezinska, Maria; Machoy- | 2007 | Roczniki Pomorskiej Akademii Medycznej w Szczecinie | Exclusion reason: No behaviour measured |

| | | | | |
|---|---|---|---|---|
| | Mokrzynska, Anna | | | |
| Influence of Assays Conditions to Glucuronidation Activity in Vitro: Psilocin Study | Manevski, Nenad; Kurkela, Mika; Hoglund, Camilla; Mauriala, Timo; Court, Michael H.; Yli-Kauhaluoma, Jari; Finel, Moshe | 2010 | Drug Metabolism Reviews | Exclusion reason: No behaviour measured |
| A comparision of the effects of two hallucinogens, psilocin and meskaline, in quantitative EEG and in senzorimotor information processing in the animal model of psychosis | | 2011 | Psychiatrie | Exclusion reason: Unavailable within timeframe of study |
| A comparision of the effects of two hallucinogens, psilocin and meskaline, in quantitative EEG and in senzorimotor information processing in the animal model of psychosis | Palenieek T.; Fujakova M.; Tyls F.; Kubesova A.; Brunovsky M.; Horacek J. | 2011 | Psychiatrie | Exclusion reason: not available in English |

| | | | | |
|---|---|---|---|---|
| Quantitative EEG in animal models of psychosis: the impact of behaviour | Palenicek, T.; Fujakova, M.; Tyls, E.; Brunovsky, M.; Kubesova, A.; Horacek, J.; Krajca, V. | 2011 | European Neuropsychopharmacology | Exclusion reason: No behaviour measured |
| Development of a discrete trials task to assess serotonergic modulation of interval timing in mice | Halberstadt A.L.; Young J.W.; Geyer M.A. | 2011 | Neuropsychopharmacology | Exclusion reason: No psilocybin/psilocin |
| Determining the pharmacokinetics of psilocin in rat plasma using ultra-performance liquid chromatography coupled with a photodiode array detector after orally administering an extract of Gymnopilus spectabilis. | Chen, Jianbo; Li, Meijia; Yan, Xitao; Wu, Enqi; Zhu, Hongmei; Lee, Kwan Jun; Chu, Van Men; Zhan, Lifeng; Lee, Wonjae; Kang, Jong Seong | 2011 | Journal of chromatography. B, Analytical technologies in the biomedical and life sciences | Exclusion reason: No behaviour measured |
| A comparison of electroencephalographic activity in serotonergic and gultamatergic models of psychosis | Tyls F.; Palenicek T.; Fujakova M.; Kubesova A.; Brunovsky M.; Horacek J.; Krajca V. | 2012 | International Journal of Neuropsychopharmacology | Exclusion reason: No behaviour measured |

| | | | | |
|---|---|---|---|---|
| Flavone Glycoside Antagonizes Psilocybin-Induced Toxicity by Reducing Oxidative Stress in Rats Model | Huang, Hong-yan; Huang, Zhe; Li, Yu-bai | 2012 | Journal of China Medical University | Exclusion reason: No behaviour measured |
| Neurovascular and neuroimaging effects of the hallucinogenic serotonin receptor agonist psilocin in the rat brain. | Spain, Aisling; Howarth, Clare; Khrapitchev, Alexandre A; Sharp, Trevor; Sibson, Nicola R; Martin, Chris | 2015 | Neuropharmacology | Exclusion reason: No behaviour measured |
| Quantitative EEG study of serotonergic hallucinogens in rats-the relationship of brain activity and behavior | Vejmola C.; Tyls F.; Kaderabek L.; Lipski M.; Palenicek T. | 2016 | European Neuropsychopharmacology | Exclusion reason: No behaviour measured |
| Psilocybin-induced psychosis in humans and in rats - translational quantitative EEG study | Tyls, F.; Vejmola, C.; Viktorinova, M.; Kaderabek, L.; Palenicek, T. | 2016 | European Neuropsychopharmacology | Exclusion reason: No behaviour measured |
| Fully Automated System to Explore the Differential Effects of Psychedelic and Non-Psychedelic Serotonin 2 A (5-HT2A) Receptor Agonists on Behavioral Tolerance | Revenga, Mario de la Fuente; Shah, Urjita; Gonzalez-Maeso, Javier | 2018 | Neuropsychopharmacology | Exclusion reason: No psilocybin/psilocin |

| | | | | |
|---|---|---|---|---|
| Psilocybin modulated expression of plasticityrelated genes and proteins in rat prefrontal cortex and hippocampus | Jefsen O.; Hojgaard K.; Elfving B.; Wegener G.; Muller H.K. | 2018 | Acta Neuropsychiatrica | Exclusion reason: No behaviour measured |
| Sex differences in serotonergic and dopaminergic mediation of LSD discrimination in rats. | Herr, Keli A | 2018 | Dissertation Abstracts International: Section B: The Sciences and Engineering | Exclusion reason: Drug discrimination |
| EEG correlates of the serotonergic hallucinogens as a parameter of assessing translational validity of the serotonergic model of psychosis in rats | Vejmola C.; Tyls F.; Lipski M.; Palenicek T. | 2018 | Clinical EEG and Neuroscience | Exclusion reason: No behaviour measured |
| Time course of quantitative EEG changes in an animal model of psilocin-induced psychosis | Tyls F.; Vejmola C.; Piorecka V.; Koudelka V.; Novak T.; Palenicek T. | 2018 | Clinical EEG and Neuroscience | Exclusion reason: No behaviour measured |
| Functional connectivity embedding for electrophysiological models of induced psychosis | Koudelka V.; Tyls F.; Vejmola C.; Brunovsky M.; Palenicek T.; Horacek J. | 2018 | Clinical EEG and Neuroscience | Exclusion reason: No behaviour measured |

| | | | | |
|---|---|---|---|---|
| Significant probability mapping on animal EEG | Piorecka V.; Tyls F.; Krajca V.; Palenicek T. | 2018 | Clinical EEG and Neuroscience | Exclusion reason: No behaviour measured |
| P.228 The polypharmacological profile of psilocybin and potential behavioural effects of very low doses | Kiilerich K.; Speth N.; Lorenz J.; Casado-Sainz A.; Shalgunov V.; Lange D.; Xiong M.; Herth M.M.; Overgaard A.; Hansen H.D.; Palner M. | 2019 | European Neuropsychopharmacology | Exclusion reason: duplicate |
| Psychedelics Improve the Mental Health of Rats | Hibicke, Meghan; Landry, Alexus N.; Talman, Zoe K.; Nichols, Charles | 2019 | Faseb Journal | Exclusion reason: duplicate |
| Psilocybin-Assisted Treatment of Major Depressive Disorder: Results From a Randomized Trial | Griffiths, Roland; Barrett, Frederick; Darrick, May; Johnson, Matthew; Mary, Cosimano; | 2019 | Neuropsychopharmacology | Exclusion reason: Human study |

| | | | | |
|---|---|---|---|---|
| | Patrick, Finan; Alan, Davis | | | |
| EEG correlates of the effect of psychedelics in rat - Spectral maps and coherence | Vejmola C.; Tyls F.; Kaderabek L.; Piorecka V.; Palenicek T. | 2019 | Neuropsychobiology | Exclusion reason: No behaviour measured |
| The effect of psilocybin on plasticity-related genes and proteins in the rat brain | Jefsen, O.; Hojgaard, K.; Elfving, B.; Wegener, G.; Muller, H. K. | 2019 | European Psychiatry | Exclusion reason: No behaviour measured |
| The MATLAB toolbox for animal 3D brain mapping and significant probability mapping | Piorecka V.; Tyls F.; Vejmola C.; Piorecky M.; Palenicek T.; Krajca V. | 2019 | Neuropsychobiology | Exclusion reason: No behaviour measured |
| The effect of psilocybin on EEG activity: Comparison of recent human findings with animal data | Palenicek T.; Tyls F.; Viktorinova M.; Androvicova R.; Brunovsky M.; Horacek J. | 2020 | Clinical EEG and Neuroscience | Exclusion reason: No behavioural/neurological outcome |

| | | | | |
|---|---|---|---|---|
| Correlation between the potency of hallucinogens in the mouse head-twitch response assay and their behavioral and subjective effects in other species. | Halberstadt, Adam L; Chatha, Muhammad; Klein, Adam K; Wallach, Jason; Brandt, Simon D | 2020 | Neuropharmacology | Exclusion reason: No psilocybin/psilocin |
| Psilocybin and LSD have no long-lasting effects in an animal model of alcohol relapse. | Meinhardt, Marcus W; Gungor, Cansu; Skorodumov, Ivan; Mertens, Lea J; Spanagel, Rainer | 2020 | Neuropsychopharmacology : official publication of the American College of Neuropsychopharmacology | Exclusion reason: Competitive |
| Sustained Reductions in Headache Burden After the Limited Administration of low Dose Psilocybin in Migraine and Cluster Headache: Results From Two Preliminary Studies | Schindler, Emmanuelle | 2020 | Neuropsychopharmacology | Exclusion reason: Human study |
| What Does the Sucrose Preference Test Tell Us About Reward Behavior? An Analysis of Licking Behavior in the SPT | Wulff, Andreas; Thompson, Scott | 2020 | Neuropsychopharmacology | Exclusion reason: No non-drug control |

| | | | | |
|---|---|---|---|---|
| Automated detection of the head-twitch response using wavelet scalograms and a deep convolutional neural network | Halberstadt, A. L. | 2020 | Sci Rep | Exclusion reason: wrong study design |
| The blue of mushrooms: Psilocybin serves as building block for protective oligomers | Anonymous. | 2020 | Deutsche Apotheker Zeitung | Exclusion reason: Could not be found after ILLs; |
| Transcriptional regulation in the rat prefrontal cortex and hippocampus after a single administration of psilocybin. | Jefsen, Oskar Hougaard; Elfving, Betina; Wegener, Gregers; Muller, Heidi Kaastrup | 2020 | Journal of psychopharmacology (Oxford, England) | Exclusion reason: No behaviour measured |
| Synthesis and Biological Evaluation of Tryptamines Found in Hallucinogenic Mushrooms: Norbaeocystin, Baeocystin, Norpsilocin, and Aeruginascin. | Sherwood, Alexander M; Halberstadt, Adam L; Klein, Adam K; McCorvy, John D; Kaylo, Kristi W; Kargbo, Robert B; Meisenheimer, Poncho | 2020 | Journal of natural products | Exclusion reason: No behaviour measured |

| | | | | |
|---|---|---|---|---|
| Integrity and Segregation of Macroscale Cerebral Functional Networks Correlate With Plasma Psilocin Level and Psychedelic Experience | Knudsen, Gitte; Madsen, Martin K.; Stenbaek, Dea S.; Arvidsson, Albin; Armand, Sophia; Marstrand-Joergensen, Maja; Johansen, Sys S.; Linnet, Kristian; Ozenne, Brice; Fisher, Patrick | 2020 | Neuropsychopharmacology | Exclusion reason: Human study |
| Examining the Acute Effect of Psilocybin in Treatment-Resistant Obsessive Compulsive Disorder | Kelmendi, Benjamin; Adams, Thomas; Depalmer, Giuliana; Kichuk, Stephen; Forte, Jennifer; Grazioplene, Rachael; Pittenger, Christopher | 2020 | Neuropsychopharmacology | Exclusion reason: Human study |

| | | | | |
|---|---|---|---|---|
| Chemoenzymatic Synthesis of 5-Methylpsilocybin: A Tryptamine with Potential Psychedelic Activity. | Fricke, Janis; Sherwood, Alexander M; Halberstadt, Adam L; Kargbo, Robert B; Hoffmeister, Dirk | 2021 | Journal of natural products | Exclusion reason: No psilocybin/psilocin |
| Structure-Activity Relationship Analysis of Psychedelics in a Rat Model of Asthma Reveals the Anti-Inflammatory Pharmacophore | Flanagan, Thomas W.; Billac, Gerald B.; Landry, Alexus N.; Sebastian, Melaine N.; Nichols, Charles D.; Cormier, Stephania A. | 2021 | ACS Pharmacology and Translational Science | Exclusion reason: No behaviour measured |
| Investigating the role of 5-$HT_{2A}$ and 5-$HT_{2C}$ receptor activation in the effects of psilocybin, DOI, and citalopram on marble burying in mice. | Odland, Anna U; Kristensen, Jesper L; Andreasen, Jesper T | 2021 | Behavioural brain research | Exclusion reason: No non-drug control |

| | | | | |
|---|---|---|---|---|
| Psilocybin exerts distinct effects on resting state networks associated with serotonin and dopamine in mice. | Grandjean, Joanes; Buehlmann, David; Buerge, Michaela; Sigrist, Hannes; Seifritz, Erich; Vollenweider, Franz X; Pryce, Christopher R; Rudin, Markus | 2021 | NeuroImage | Exclusion reason: No behaviour measured |
| Discriminative stimulus effects of substituted tryptamines in rats | Gatch M.B.; Hoch A.; Carbonaro T.M. | 2021 | ACS Pharmacology and Translational Science | Exclusion reason: Drug discrimination |
| Effects of a single dose of psilocybin on behaviour, brain 5-$HT_{2A}$ receptor occupancy and gene expression in the pig. | Donovan, Lene Lundgaard; Johansen, Jens Vilstrup; Ros, Nidia Fernandez; Jaberi, Elham; Linnet, Kristian; Johansen, Sys Stybe; Ozenne, Brice; Issazadeh-Navikas, | 2021 | European neuropsychopharmacology : the journal of the European College of Neuropsychopharmacology | Exclusion reason: No behaviour measured |

| | | | | |
|---|---|---|---|---|
| | Shohreh; Hansen, Hanne Demant; Knudsen, Gitte Moos | | | |
| Transcriptional regulation in the rat prefrontal cortex and hippocampus after a single administration of psilocybin. | Jefsen, Oskar Hougaard; Elfving, Betina; Wegener, Gregers; Muller, Heidi Kaastrup | 2021 | Journal of psychopharmacology (Oxford, England) | Exclusion reason: No behaviour measured |
| Preclinical screening for antidepressant activity - shifting focus away from the Forced Swim Test to the use of translational biomarkers | Sewell, Fiona; Waterson, Ian; Jones, David; Tricklebank, Mark David; Ragan, Ian | 2021 | Regulatory Toxicology and Pharmacology | Exclusion reason: No psilocybin/psilocin |
| A Single Dose of Psilocybin Increases Synaptic Density and Decreases 5-$HT_{2A}$ Receptor Density in the Pig Brain. | Raval, Nakul Ravi; Johansen, Annette; Donovan, Lene Lundgaard; Ros, Nidia Fernandez; Ozenne, Brice; | 2021 | International journal of molecular sciences | Exclusion reason: No behaviour measured |

| | |
|---|---|
| | Hansen, Hanne Demant; Knudsen, Gitte Moos |

**List of Excluded Studies (Full-text Review): BIPA**

| Number | Source | Reason for exclusion post full-text review |
|---|---|---|
| 1. | Adey, W. R., Bell, F. R. & Dennis, B. J. Effects of LSD-25, psilocybin, and psilocin on temporal lobe EEG patterns and learned behavior in the cat. *Neurology*. **12**, 591-602 (1962). | Could not be found after ILLs (inter-library loans). |
| 2. | Adron Harris, R., Homfeld, E. & Campaigne, E. Structure-activity relationships among hallucinogenic tryptamine derivatives evaluated by schedule-controlled behaviour. *J. Pharm. Pharmacol.* **33**, 320–322 (1981). | Duplicate |
| 3. | Aghajanian, G. K. & Hailgler, H. J. Hallucinogenic indoleamines: preferential action upon presynaptic serotonin receptors. *Psychopharmacol. Commun.* **1**, 619–629 (1975). | No behavior measured – "new criterion" |
| 4. | Ahn, H. S. & Makman, M. H. Stimulation of adenylate cyclase activity in monkey anterior limbic cortex by serotonin. *Brain Res.* **153**, 636 | No behavior measured – "new criterion" |
| 5. | Allgren, R. L., Kyncl, M. M. A. & Ciaranello, R. D. Pharmacological characterization of solubilized 5-HT1 serotonin binding sites from bovine brain. *Brain Res.* **348**, 77–85 (1985). | No behavior measured – "new criterion" |
| 6. | Anden, N. E., Corrodi, H. & Fuxe, K. Hallucinogenic drugs of the indolealkylamine type and central monoamine neurons. *J. Pharmacol. Exp. Ther.* **179**, 236–249 (1971). | No non-drug conrol |
| 7. | Balzamo, E. The baboon Papio papio human epilepsy model pharmacological aspects. *Stal* **1**, 317–323 (1976). | Could not be found after ILLs |
| 8. | Balzano, E., Meldrum, B. S., Naquet, R. & Vuillon-Cacciuttolo, G. The effects of various ergot derivatives on the electro encephalogram and photo-sensitivity in Papio papio. *Electroencephalogr. Clin. Neurophysiol.* **32**, 578 (1972). | No psilocybin/psilocin |
| 9. | Bermond, F., Bert, J. & Ayats, J. Effects of psilocybine on visual evoked potentials in a cercopithecinae Papio papio. *C. R. Seances Soc. Biol. Fil.* **160**, 2405–2409 (1966). | Not available in English |
| 10. | Blair, J. B. *et al.* Effect of ring fluorination on the pharmacology of hallucinogenic tryptamines. *J. Med. Chem.* **43**, 4701–4710 (2000). | No psilocybin/psilocin |
| 11. | Bourn, W. ., Keller, W. J. & Bonfiglio, J. F. Comparisons of the effects of mescal bean alkaloids with mescaline, N,N-Dimethyltryptamine (DMT), psilocybin, amphetamine, delta-9-tetrahydrocannabinol (THC), and pentobarbital in rats. *Fed. Proc.* **38**, 590 (1979). | Not original research (source study) |

| | | |
|---|---|---|
| 12. | Brodey, J. F., Steiner, W. G. & Himwich, H. E. An electrographic study of psilixcin and 4-methyl-alpha-methyl-tryptamine (MP-809). *J. Pharmacol. Exp. Ther.* **140**, 8–18 (1963). | No behavior measured – “new criterion” |
| 13. | Buckholtz, N. S., Zhou, D., Freedman, D. X. & Potter, W. Z. Lysergic acid diethylamide (LSD) administration selectively downregulates serotonin2 receptors in rat brain. *Neuropsychopharmacology* **3**, 137–148 (1990). | No behavior measured – “new criterion” |
| 14. | Burt, D. R., Creese, I. & Snyder, S. H. Binding interactions of lysergic acid diethylamide and related agents with dopamine receptors in the brain. *Mol. Pharmacol.* **12**, 631–638 (1976). | No behavior measured – “new criterion” |
| 15. | Callahan, P. M. & Appel, J. B. Differences in the stimulus properties of 3,4-Methylenedioxyamphetamine and 3,4-Methylenedioxymethamphetamine in animals trained to discriminate hallucinogens from saline. *J. Pharmacol. Exp. Ther.* **246**, 866–870 (1988). | No non-drug control |
| 16. | Chen, J. *et al.* Determining the pharmacokinetics of psilocin in rat plasma using ultra-performance liquid chromatography coupled with a photodiode array detector after orally administering an extract of Gymnopilus spectabilis. *J. Chromatogr. B Anal. Technol. Biomed. Life Sci.* **879**, 2669–2672 (2011). | No behavior measured – “new criterion” |
| 17. | Collins, B. J., Lovell, R. A., Boggan, W. O. & Freedman, D. X. Effects of hallucinogens on rat brain monoamine oxidase activity. *Pharmacologist* **12**, 256 (1970). | No behavior measured – “new criterion” |
| 18. | Consroe, P. & Fish, B. S. Pharmacological characteristics of tetrahydrocannabinol seizure susceptible rabbits. *Fed. Proc.* **39**, 3065 (1980). | Could not be found after ILLs |
| 19. | Corne, S. J. & Pickering, R. W. A possible correlation between drug-induced hallucinations in man and a behavioural response in mice. *Psychopharmacologia* **11**, 65–78 (1967). | Competitive |
| 20. | Creese, I., Burt, D. R. & Snyder, S. H. The dopamine receptor: differential binding of d-LSD and related agents to agonist and antagonist states. *Life Sci.* **17**, 1715–1720 (1975). | No behavior measured – “new criterion” |
| 21. | Curtis, D. R. Psychotomimetics - a neuropharmacological study. *Proc. Aust. Assoc. Neurol.* **1**, 43–45 (1963). | Not original research (source study) |
| 22. | Fairchild, M. D., Jenden, D. J., Mickey, M. R. & Yale, C. EEG effects of hallucinogens and cannabinoids using sleep-waking behavior as baseline. *Pharmacol. Biochem. Behav.* **12**, 99–105 (1980). | No behavior measured – “new criterion” |
| 23. | Fantegrossi, W. E., Woods, J. H. & Winger, G. Transient reinforcing effects of phenylisopropylamine and indolealkylamine hallucinogens in rhesus monkeys. *Behav. Pharmacol.* **15**, 149–157 (2004). | Drug discrimination |
| 24. | Fredrickson, P. A. & Richelson, E. Hallucinogens antagonize histamine H-1 receptors of cultured mouse neuro blastoma cells. *Eur. J. Pharmacol.* **56**, 261–264 (1979). | No behavior measured – “new criterion” |
| 25. | Freedman, D. X., Gottlieb, R. & Lovell, R. A. Psychotomimetic drugs and brain 5-hydroxytryptamine metabolism. *Biochem. Pharmacol.* **19**, 1181–1188 (1970). | No behavior measured – “new criterion” |
| 26. | Gessner, P. K., Khairallah, P. A., McIsaac, W. M. & Page, I. H. The relationship between the metabolic fate and pharmacological actions of serotonin, bufotenine and psilocybin. *J. Pharmacol. Exp. Ther.* **130**, 126–133 (1960). | No behavior measured – “new criterion” |

| | | |
|---|---|---|
| 27. | Glennon, R. A., Young, R., Rosecrans, J. A. & Kallman, M. J. Hallucinogenic agents as discriminative stimuli: A correlation with serotonin receptor affinities. *Psychopharmacology (Berl).* **68**, 155–158 (1980). | Drug discrimination |
| 28. | Goldman, H. & Fischer, H. Cortical and/or subcortical effects as a function of hallucinogenic drug structure. *Pharmacologist* **16**, 237 (1974). | No psilocybin/psilocin |
| 29. | Goldstein, M. L., Lovell, R. A., Boggan, W. O. & Freedman, D. X. Effect of LSD-25 and psilocybin on norepinephrine and 3H-norepinephrine metabolites in rat brain. *Fed. Proc.* **29**, A650 (1970). | No behavior measured – "new criterion" |
| 30. | González-Maeso, J. *et al.* Hallucinogens Recruit Specific Cortical 5-$HT_{2A}$ Receptor-Mediated Signaling Pathways to Affect Behavior. *Neuron* **53**, 439–452 (2007). | No psilocybin/psilocin |
| 31. | Graf, M. & Pietscher, A. Shape change of blood platelets: a model for cerebral 5-hydroxytryptamine receptors? *Br. J. Pharmacol.* **65**, 601–608 (1979). | No behavior measured – "new criterion" |
| 32. | Green, J. P., Weingartner, H. & Maayani, S. Defining the histamine H2-receptor in brain: the interaction with LSD. *NIDA Res. Monogr.* 38–59 (1978). | No behavior measured – "new criterion" |
| 33. | Greenberg, I., Kuhn, D. & Appel, J. B. Comparison of the discriminative stimulus properties of clonidine and amphetamine in rats. *Pharmacol. Biochem. Behav.* **3**, 931–934 (1975). | Drug discrimination |
| 34. | Haefely, W. The effects of 5-hydroxytryptamine and some related compounds on the cat superior cervical ganglion in situ. *Naunyn. Schmiedebergs. Arch. Pharmacol.* **281**, 145–165 (1974). | No behavior measured – "new criterion" |
| 35. | Haefely, W. Effects of serotonin (5-HT) and some related indole compounds in a mammalian sympathetic ganglion. *Exp.* **27**, 1112 (1972). | No behavior measured – "new criterion" |
| 36. | Halaris, A. E. Nerve terminal effects of indoleamine psychotomimetics on 5-hydroxytryptamine. *Neurosci. Biobehav. Rev.* **6**, 483–487 (1982). | No behavior measured – "new criterion" |
| 37. | Halasz, M. F. & Marrazzi, A. S. Disinhibition of conditioned behavior by cerebral synaptic inhibitors. *Pharmacologist* **7**, 173 (1965). | No behavior measured – "new criterion" |
| 38. | Halberstadt, A. L., Young, J. W. & Geyer, M. A. Development of a discrete trials task to assess serotonergic modulation of interval timing in mice. *Neuropsychopharmacology.* **36**, S210–S211 (2011). | No psilocybin/psilocin |
| 39. | Hasler, F., Dobricki, M., Grimber, U. & Vollenweider, F. X. Cognitive and psychopathological aspects of the 5-$HT_{2A}$ model of experimental psychosis. *J. Psychopharmacol.* **17**, A44 (2003). | No mammals |
| 40. | Herblin, W. F. & O'Brien, R. D. Interactions of norepinephrine with subcellular fractions of rat brain I. Characteristics of norepinephrine uptake. *Brain Res.* **8**, 298–309 (1968). | No behavior measured – "new criterion" |
| 41. | Herr, K. A. Sex differences in serotonergic and dopaminergic mediation of LSD discrimination in rats. *Diss. Abstr. Int. Sect. B Sci. Eng.* **79**, No-Specified (2018). | Drug discrimination |
| 42. | Hopf, A. & Eckert, H. Distribution patterns of 14-C-psilocin in the brains of various animals. *Act. Nerve Super.* **16**, 64–66 (1974). | No behavior measured – "new criterion" |

| | | |
|---|---|---|
| 43. | Hopf, A. & Eckert, H. Autoradiographic studies on the distribution of psychoactive drugs in the rat brain. *Psychopharmacologia* **16**, 201–222 (1969). | No behavior measured – “new criterion” |
| 44. | Horita, A. Some biochemical studies on psilocybin and psilocin. *J. Neuropsychiatry* **4**, 270–273 (1963). | No behavior measured – “new criterion” |
| 45. | Huang, H., Huange, Z. & Li, Y. Flavone glycoside antagonizes psilocybin-induced toxicity by reducing oxidative stress in rat models. *J. China Med. Univ.* **41**, 48–51 (2012). | No behavior measured – “new criterion” |
| 46. | Hungen, K. V., Roberts, S. & Hill, D. F. Interactions between lysergic acid diethylamide and dopamine sensitive adenylate cyclase systems in rat brain. *Brain Res.* **94**, 57–66 (1975). | No behavior measured – “new criterion” |
| 47. | Jacob, J. & Lafille, C. Pharmacological detection and characterization of hallucinogens. I. Hyperthermizing activities in the rabbit. *Arch. Int. Pharmacodyn. Ther.* **145**, 528–545 (1963). | Not available in English |
| 48. | Jacob, J., Lafille, C., Loiseau, G., Echinard-Garin, P. & Barthelemy, C. Search for experimental equivalents of hallucinogenic activity. Effects of hallucinogens on rectal temperature in rabbit and on morphine effects in mouse. **7**, 296–304 (1962). | Could not be found after ILLS |
| 49. | Jacob, J., Lafille, C., Loiseau, G., Echinard-Garin, P. & Barthelemy, C. Research on the pharmacological characterization and differentiation of hallucinogenic drugs (indole derivatives and mescaline, nalorphine, central anticholinergics, and phencyclidine). *L’Encephale Rev. Psychiatr. Clin. Biol. Ther.* **53**, 520–535 (1964). | Not available in English |
| 50. | Jefsen, O., Hojgaard, K., Elfving, B., Wegener, G. & Muller, H. K. The effect of psilocybin on plasticity-related genes and proteins in the rat brain. *Eur. Psychiatry* **56**, S166 (2019). | No behavior measured – “new criterion” |
| 51. | Jefsen, O., Hojgaard, K., Elfving, B., Wegener, G. & Muller, H. K. Psilocybin modulated expression of plasticity related genes and proteins in rat prefrontal cortex and hippocampus. *Acta Neuropschiatrica* **30**, 13–14 (2018). | No behavior measured – “new criterion” |
| 52. | Jenner, P., Luscombe, G. & Marsden, C. D. Interaction of 5-HT agonists with guinea-pig brain stem 5-HT receptors and the induction of myoclonus. *Br. J. Pharmacol.* **80**, 667P (1983). | No behavior measured – “new criterion” |
| 53. | Kakajima, H. & Thuillier, J. EEG effects of psychoanaleptics and biogenic amine precursors in reserpinized rabbits. **14**, 161–168 (1966). | No behavior measured – “new criterion” |
| 54. | Kakolewski, J. W. Differentiation of EEG ‘deactivation’. *Physiol. Behav.* **3**, 503–506 (1968). | No behavior measured – “new criterion” |
| 55. | Kalberer, F., Kreis, W. & Rutschmann, J. The fate of psilocin in the rat. *Biochem. Pharmacol.* **11**, 261–269 (1962). | No behavior measured – “new criterion” |
| 56. | Katagiri, N. *et al.* Effects of systemic psilocin on serotonin release in rat brain. *J. Pharmacol. Sci.* **100**, 141P (2006). | No behavior measured – “new criterion” |
| 57. | Kato, L., Gözsy, B., Ban, T. A. & Sterlin, C. Effects of psychoactive agents on the conditioning of the microcirculation in the rat. *Cond. Reflex* **6**, 67–77 (1971). | No behavior measured – “new criterion” |

| | | |
|---|---|---|
| 58. | Koerner, J. & Appel, J. B. Psilocybin as a discriminative stimulus: Lack of specificity in an animal behavior model for 'hallucinogens'. *Psychopharmacology (Berl).* **76**, 130–135 (1982). | Drug discrimination |
| 59. | Kolarik, J. Different effects of psilocybin on the electroencephalogram of various types of epilepsy. *Electroencephalogr. Clin. Neurophysiol.* **36**, 80 (1974). | No behavior measured – "new criterion" |
| 60. | Koudelka, V. *et al.* Functional connectivity embedding for electrophysiological models of induced psychosis. *Clin. EEG Neurosci.* **49**, NP20–NP21 (2018). | No behavior measured – "new criterion" |
| 61. | Koukkou, M. & Lehmann, D. Experience of induced visual hallucinations depends on pre-treatment EEG spectra. *Neurosci. Lett.* S190 (1979). | No mammals |
| 62. | Kurrasch-Orbaugh, D., Watts, V. J., Barker, E. L. & Nichols, D. E. Serotonin 5-hydroxytryptamine2A receptor-coupled phospholipase C and phospholipase A2 signalling pathways have different receptor reserves. *J. Pharmacol. Exp. Ther.* **304**, 229–237 (2003). | No behavior measured – "new criterion" |
| 63. | Ladefoged, O. The effects of LSD, psilocybin, harmaline and amphetamine on the body temperature of para chlorophenylalanine pretreated rats. *Archives Internationales de Pharmacodynamie et de Therapie* vol. 204 326–332 (1973). | No behavior measured – "new criterion" |
| 64. | Ladefoged, O. The effect of LSD, psilocybin, harmaline and amphetamine on the body temperature of para-chlorophenylalanine pretreated rabbits. *Arch. Int. Pharmacodyn. Ther.* **208**, 251–254 (1974). | No behavior measured – "new criterion" |
| 65. | Laubscher, A. & Pletscher, A. Increase of cyclic GMP in blood platelets by biogenic amines a receptor mediated effect. *J. Pharm. Pharmacol.* **32**, 601–602 (1980). | No behavior measured – "new criterion" |
| 66. | Lewis, J. J., Ritchie, A. P. & Van Petten, G. R. The influence of hallucinogenic drugs upon in vivo brain levels of adenine nucleotides, phosphocreatine and inorganic phosphate in the rat. *Br. J. Pharmacol. Chemother.* **25**, 631–637 (1965). | No behavior measured – "new criterion" |
| 67. | Luscombe, G., Jenner, P. & Mardsen, C. D. 5HT dependent myoclonus in guinea pigs is induced by 5HT agonists containing an indole nucleus, but not by those possessing a piperazine moiety. *Neurosci. Lett.* **24**, S23 (1981). | No behavior measured – "new criterion" |
| 68. | Luscombe, G., Jenner, P. & Mardsen, C. D. 5 hydroxytryptamine dependent myoclonus in guinea-pigs is induced by 5 hydroxytryptamine agonists containing an indole nucleus but not by those possessing a piperazine moiety. *Neurosci. Lett.* S23 (1981). | Not original research (source study) |
| 69. | Luscombe, G., Jenner, P. & Marsden, C. D. University Department of Neurology, Institute of Psychiatry and King's College Hospital Medical School, Denmark Hill, London SE5, U.K. *Eur. J. Pharmacol.* **104**, 235–244 (1984). | Not original research (source study) |
| 70. | Machoy-Mokrzynska, A. *et al.* The influence of psilocin and phenylethylamine on the energy metabolism in the rat heart. *Acta Toxicol.* **11**, 13–19 (2003). | No behavior measured – "new criterion" |
| 71. | Majdanik, S., Borowiak, K., Brzezinska, M. & Machoy-Mokrzynska, A. Concentration of selected microelements in blood serum or rats exposed to the action of psilocin and phenylethylamine. *Rocz. Pomor. Akad. Med. w Szczecinie* **53**, 153–158 (2007). | No behavior measured – "new criterion" |

| | | |
|---|---|---|
| 72. | Manevski, N. *et al.* Influence of assays conditions to glucuronidation activity in vitro: psilocin study. *Drug Metab. Rev.* **42**, 64 (2010). | No behavior measured – "new criterion" |
| 73. | Marchbanks, R. M. Inhibitory effects of lysergic acid derivatives and reserpine on 5-HT binding to nerve ending particles. *Biochem. Pharmacol.* **16**, 1971–1979 (1967). | No behavior measured – "new criterion" |
| 74. | Martin, W. R. & Eades, C. G. The action of tryptamine on the dog spinal cord and its relationship to the agonistic actions of LSD-like psychotogens. *Psychopharmacologia* **17**, 242–257 (1970). | Competitive |
| 75. | Martin, W. R. & Sloan, J. W. Relationship of CNS tryptaminergic processes and the action of LSD-like hallucinogens. *Pharmacol. Biochem. Behav.* **24**, 393–399 (1986). | No non-drug control |
| 76. | Martin, W. R. & Eades, C. G. Tryptamine receptors in dog spinal cord and their relationship to the agonistic actions of lysergic-acid diethylamide like psychotogens. *Pharmacologist* **10**, 196 (1968). | No behavior measured – "new criterion" |
| 77. | Martin, W. R. & Sloan, J. W. The possible role of tryptamine in brain function and its relationship to the actions of LSD-like hallucinogens. *Mt. Sinai J. Med.* **41**, 276–282 (1974). | No psilocybin/psilocin |
| 78. | Martin, W. R. & Sloan, J. W. Relationship of central nervous system tryptaminergic processes and the action of LSD-like hallucinogens. *Pharmacol. Biochem. Behav.* **24**, 393–400 (1986). | Not original research (source study) |
| 79. | Maxwell, G. M., Burnell, R. H. & Kneebone, G. M. The effects of tryptamine and alpha-methyl-tryptamine on the general and coronary haemodynamics and metabolism of the dog. *Arch Int Pharmacodyn Ther.* **153**, 79–86 (1965). | No behavior measured – "new criterion" |
| 80. | Maxwell, G. M., Kneebone, G. M. & Elliot, R. B. The effect of psilocybin upon the systemic, pulmonary and coronary circulation of the intact dog. *Arch. Int. Pharmacodyn. Ther.* **137**, 108–115 (1962). | No behavior measured – "new criterion" |
| 81. | McCall, R. B. & Aghajanian, G. K. Hallucinogens potentiate responses to serotonin and norepinephrine in the facial motor nucleus. *Life Sci.* **26**, 1149–1156 (1980). | No behavior measured – "new criterion" |
| 82. | McCloskey, K. L. & Franz, D. N. Effects of LSD mescaline and psilocybin on sympathetic preganglioinic neurons. *Pharmacologist* **16**, 237 (1974). | No behavior measured – "new criterion" |
| 83. | McKenna, D. J., Repke, D. B., Lo, L. & Peroutka, S. J. Differential interactions of indolealkylamines with 5-hydroxytryptamine receptor subtypes. *Neuropharmacology* **29**, 193–198 (1990). | No behavior measured – "new criterion" |
| 84. | Meldrum, B. S. & Naquet, R. Effects of psilocybin, dimethyltryptamine, mescaline and various lysergic acid derivatives on the EEG and on photically induced epilepsy in the baboon (Papio papio). *Electroencephalogr. Clin. Neurophysiol.* **31**, 563–572 (1971). | Duplicate |
| 85. | Meldrum, B. S. & Naquet, R. Effects of psilocybin, dimethyltryptamine and various lysergic acid derivatives on photically-induced epilepsy in baboon (Papio papio). *Br. J. Pharmacol.* **40**, 144–145 (1970). | Duplicate |
| 86. | Meltzer, H. Y., Fessler, R. G., Simonovic, M. & Fang, V. S. Stimulation of rat prolactin secretion by indolealkylamine hallucinogens. *Psychopharmacology (Berl).* **56**, 255–259 (1978). | No behavior measured – "new criterion" |

| | | |
|---|---|---|
| 87. | Meltzer, H. Y., Fessler, R. G., Simonovic, M. & Fang, V. S. Effect of indole hallucinogens, mesacline and DMPEA on rat plasma protein. *Fed. Proc.* **36**, (1977). | No behavior measured – "new criterion" |
| 88. | Moldavan, M. G. *et al.* The effect of psilocybe cubensis extract on hippocampal neurons in vitro. *Fiziol. Zh.* **47**, 15–23 (2001). | No behavior measured – "new criterion" |
| 89. | Monnier, M. The effect of the action of psilocybin on the rabbit brain. *Experentia* **15**, 321–323 (1959). | Not available in English |
| 90. | Nakajima, H. & Thuillier, J. Changes in the EEG in the reserpinized rabbit caused by psychoanaleptics and the precursors of biogenic amines. *Med Pharmacol Exp Int J Exp Med* **14**, 161–168 (1966). | No behavior measured – "new criterion" |
| 91. | Nomura, J. Effects of stress and psychotropic drugs on rat liver tryptophan pyrrolase. *Endocrinology* **76**, 1190–1194 (1965). | No behavior measured – "new criterion" |
| 92. | Palenicek, T., Bubenikova, V. & Horacek, J. Modelling of psychotic-like behavior: comparison of MK-801 with psilocin, LSD, mescaline and 2C-B models. | No non-drug control |
| 93. | Palenicek, T. *et al.* Quantitative EEG in animal models of psychosis: the impact of behaviour. *Eur. Neuropsychopharmacol.* **21**, S317–S3 | No behavior measured – "new criterion" |
| 94. | Palenicek, T. *et al.* A comparison of the effects of two hallucinogens, psilocin and meskaline, in quantitative EEG and in sensorimotor information | Not available in English |
| 95. | Persson, S. A. LSD and related drugs as dopamine antagonists receptor mediated effects on the synthesis and turnover of dopamine. *Life Sci.* **23**, 523–526 (1978). | No behavior measured – "new criterion" |
| 96. | Persson, S.-A. LSD and related drugs as DA antagonists: receptor-mediated effects on teh synthesis and turnover of DA. *Life Sci.* **23**, 523–526 (1978). | No behavior measured – "new criterion" |
| 97. | Petkov, V., Shoumkov, G. & Koushev, V. Effect of certain psychopharmacologic substances on S35-methionine cytoplasm incorporation (cyrillic). *Savrem. Med.* **17**, 461–470 (1966). | No behavior measured – "new criterion" |
| 98. | Piorecka, V., Tyls, F., Krajca, V. & Palenicek, T. Significant probability mapping on animal EEG. *Clin. EEG Neurosci.* **49**, NP22 (2018). | No behavior measured – "new criterion" |
| 99. | Piorecka, V. *et al.* The MATLAB toolbox for animal 3D brain mapping and significant probability mapping. *Neuropsychobiology* **77**, 149–150 (2019). | No behavior measured – "new criterion" |
| 100. | Powell, S. B. *et al.* The indoleamine hallucinogens, psilocin and 5MeODMT, increase prepulse inhibition in mice: rol eof 5-HT1A and 5-HT$_{2A}$ receptors. *Soc. Neurosci.* Abstract No. 315.7 (2003). | Could not be found after ILLs |
| 101. | Rabin, R. A., Regina, M., Doat, M. & Winter, J. C. 5-HT$_{2A}$ receptor-stimulated phosphoinositide hydrolysis in the stimulus effects of hallucinogens. *Pharmacol. Biochem. Behav.* **72**, 29–37 (2002). | No behavior measured – "new criterion" |
| 102. | Rolsten, C. Effects of chlorpromazine and psilocin on pregnancy of C57BL/10 mice and their offspring at birth. *Anat. Rec.* **157**, 311 (1967). | No behavior measured – "new criterion" |
| 103. | Ruch-Monachon, M. A., Jalfre, M. & Haefely, W. Drugs and PGO waves in the lateral geniculate body of the curarized cat. II. PGO wave activity and brain 5-hydroxytryptamine. *Arch Int Pharmacodyn Ther* **219**, 269–286 (1976). | No behavior measured – "new criterion" |

| | | |
|---|---|---|
| 104. | Seeman, P., Westman, K., Coscina, D. & Warsh, J. J. Serotonin receptors in hippocampus and frontal cortex. *Eur. J. Pharmacol.* **66**, 179–191 (1980). | No behavior measured – “new criterion” |
| 105. | Shein, H. M., Wilson, S., Larin, F. & Wurtman, R. J. Stimulation of [14C] serotonin synthesis from [14C] tryptophan by mescaline in rat pineal organ cultures. *Life Sci.* **10**, 273–282 (1971). | No behavior measured – “new criterion” |
| 106. | Shein, H. M., Wilson, S., Larin, F. & Wurtman, R. J. Stimulation of 14C serotonin synthesis from 14C tryptophan by mescaline in rat pineal organ cultures. *Life Sci.* **10**, 273–282 (1971). | Duplicate |
| 107. | Silverman, P. B. & Ho, B. T. Stimulus properties of commonality with other hallucinogens 2,5-dimethoxy-4-methyl amphetamine. *Psychopharmacology (Berl).* **58**, 10 (1978). | Could not be found after ILLs |
| 108. | Silverman, P. B. & Ho, B. T. Discriminative response control by psychomotor stimulants. *Psychopharmacol. Commun.* **2**, 331–337 (1976). | Drug discrimination |
| 109. | Spain, A. *et al.* Neurovascular and neuroimaging effects of the hallucinogenic serotonin receptor agonist psilocin in the rat brain. *Neuropharmacology* **99**, 210–220 (2015). | No behavior measured – “new criterion” |
| 110. | Steiner, J. E. & Sulman, F. G. Simultaneous studies of blood sugar, behavioural changes and EEG on wake rabbit after administration of psilocybin. *Arch. Int. Pharmacodyn. Ther.* **145**, 301 (1963). | Could not be found after ILLs |
| 111. | Stolk, J. M., Barchas, J. D., Goldstein, M., Boggan, W. & Freedman, D. X. A comparison of psychotomimetic drug effects on rat brain norepinephrine metabolism. *J. Pharmacol. Exp. Ther.* **189**, 42–50 (1974). | No behavior measured – “new criterion” |
| 112. | Tilson, H. A., Marquis, W. J. & Rech, R. H. The effects of d-amphetamine, 2,5-dimethoxy-4-methyl-amphetamine (DOM), and psilocybin on fixed-interval responding in the rat. *Proc. Annu. Conv. Am. Pyschological Assoc.* 1007–1008 (1973). | Could not be found after ILLs |
| 113. | Trulson, V. M. & Trulson, M. E. Differential effects of indoleamine hallucinogens on serotonin containing neurons in the nucleus centralis superior and nucleus raphe pallidus in freely moving cats. *Fed. Proc.* **42**, 5036 (1983). | No behavior measured – “new criterion” |
| 114. | Tyls, F. *et al.* A comparison of electroencephalographic activity in serotonergic and glutamatergic models of psychosis. *Int. J. Neuropsychopharmacol.* **15**, 211 (2012). | No behavior measured – “new criterion” |
| 115. | Tyls, F. *et al.* Time course of quantitative EEG changes in an animal model of psilocin-induced psychosis. *Clin. EEG Neurosci.* **49**, NP | No behavior measured – “new criterion” |
| 116. | Tyls, F., Vejmola, C., Viktorinova, M., Kaderabek, L. & Palenicek, T. Psilocybin-induced psychosis in humans and in rats − translational quantitative EEG study. *Eur. Neuropsychopharmacol.* **26**, S257–S258 (2016). | No behavior measured – “new criterion” |
| 117. | Uyeno, E. T. Inhibition of isolation-induced attack behavior of mice by drugs. *Proc. West. Pharmacol. Soc.* **9**, (1966). | Not original research (source study) |
| 118. | Uyeno, E. T. Hallucinogens and dominance behavior in the rat. *Proc. West. Pharmacol. Soc.* **10**, (1967). | Not original research (source study) |
| 119. | Uyeno, E. T. Effects of psycho dysleptics on aggressive behavior of animals. *Mod. Probl. Pharmacopsych.* **13**, 103–113 (1978). | Not original research (source study) |

| | | |
|---|---|---|
| 120. | Uzunov, P. & Weiss, B. Effect of pyschotomimetic drugs on the cyclic 3 5 AMP system of rat brain. *Pharmacologist* **13**, 257 (1971). | No behavior measured – “new criterion” |
| 121. | Vaupel, D. B. & Martin, W. R. LSD-like hallucinogens in the dog: validation studies with mescaline (MES), psilocin (PSI) and dimethyltryptamine (DMT). *Fed. Proc.* **37**, (1978). | Not original research (source study) |
| 122. | Vejmola, C., Tylš, F., Kadeřábek, L., Lipski, M. & Páleníček, T. Quantitative EEG study of serotonergic hallucinogens in rats − the relationship of brain activity and behavior. *Eur. Neuropsychopharmacol.* **26**, S260–S261 (2016). | No behavior measured – “new criterion” |
| 123. | Vejmola, C., Tylš, F., Kaderabek, L., Piorecka, V. & Palenicek, T. EEG correlates of the effect of psychedelics in rat - spectral maps and coherence. *Neuropsychobiology* **77**, 142–143 (2019). | No behavior measured – “new criterion” |
| 124. | Vejmola, C., Tyls, F., Lipski, M. & Palenicek, T. EEG correlates of the serotonergic hallucinogens as a parameter of assessing translational validity of the serotonergic model of psychosis in rats. *Clin. EEG Neurosci.* **49**, NP22 (2018). | No behavior measured – “new criterion” |
| 125. | Vojtechovsky, M., Hort, V. & Safratove, V. Effect of monoamine oxidase inhibitors on experimental psychoses following the administration of psilocybin. *Act. Nerve Super.* **10**, 278–279 (1968). | Could not be found after ILLs |
| 126. | Von Hungen, K., Roberts, S. & Hill, D. F. Serotonin sensitive adenylate cyclase activity in immature rat brain. *Brain Res.* **84**, 257–267 (1975). | No behavior measured – “new criterion” |
| 127. | Wallis, D. I., Stansfeld, C. E. & Nash, H. L. Depolarizing responses recorded from nodose ganglion cells of the rabbit evoked by 5-hydroxytryptamine and other substances. *Neuropharmacology* **21**, 31–40 (1982). | No behavior measured – “new criterion” |
| 128. | Waser, P. G. & Schaub, E. The action of some neuro- and psychopharmacological agents on the membrane ATP-ase of cortical synaptosomes. *Adv Cytopharmacol.* **1**, 397–400 (1971). | No behavior measured – “new criterion” |
| 129. | Waser, P. G. & Schaub, E. Action of some neuro drugs and psycho pharmaca drugs on membrane ATPase and acetylcholine esterase of cortical synaptosomes. *Heilbronn, Ed. Anders Winter* 359–368 (1970). | No behavior measured – “new criterion” |
| 130. | Weidmann, H. & Cerletti, A. Studies on psilocybin and related compounds. I. Communication. Structure/activity relationship of oxyindole-derivatives with regard to their effect on the knee jerk of spinal cats. *Helv. Physiol. Pharmacol. Acta* **18**, 174–182 (1960). | No behavior measured – “new criterion” |
| 131. | Weidmann, H., Taeschler, M. & Konzett, H. Pharmacology of psilocybin, a drug from psilocybe mexicana heim. *Experientia* **14**, 378–379 (1958). | Not available in English |
| 132. | Whipple, M. R., Reinecke, M. G. & Gage, F. H. Inhibition of synaptosomal neurotransmitter uptake by hallucinogens. *J. Neurochem.* **40**, 1185–1188 (1983). | No behavior measured – “new criterion” |
| 133. | Whitaker, P. M. & Seeman, P. High affinity tritiated serotonin binding to caudate inhibition by hallucinogens and serotoninergic drugs. *Psychopharmacology (Berl).* **59**, 1–6 (1978). | No behavior measured – “new criterion” |
| 134. | Whitaker, P. M. & Seeman, P. Selective labeling of serotonin receptors by d-[3H]lysergic acid diethylamide in calf caudate. *Proc. Natl. Acad. Sci. United States Am.* **75**, 5783–5787 (1978). | No behavior measured – “new criterion” |
